# Selective pressure on membrane proteins drives the evolution of *Helicobacter pylori* Colombian subpopulations

**DOI:** 10.1101/2021.12.14.472690

**Authors:** Alix A. Guevara, Roberto C. Torres, Fabian L. Castro-Valencia, John J. Suárez, Angel Criollo-Rayo, Maria Mercedes Bravo, Luis Carvajal Carmona, M. Magdalena Echeverry de Polanco, Mabel E. Bohórquez, Javier Torres

## Abstract

*Helicobacter pylori* have coevolved with mankind since its origins, adapting to different human groups. In America *H. pylori* has evolved in several subpopulations specific for regions or even countries. In this study we analyzed the genome of 163 Colombian strains along with 1,113 strains that represent worldwide *H. pylori* populations to better discern the ancestry and adaption to Colombian people. Population structure was inferred with FineStructure and chromosome painting identifying the proportion of ancestries in Colombian isolates. Phylogenetic relationship was analyzed using the SNPs present in the core genome. Also, a Fst analysis was done to identify the gene variants with the strongest fixation in the identified Colombian subpopulations in relation to their parent population *hspSWEurope*. Worldwide, population structure analysis allowed the identification of two Colombian subpopulations, the previously described *hspSWEuropeColombia* and a novel subpopulation named *hspColombia*. In addition, three subgroups of *H. pylori* were identified within *hspColombia* that follow their geographic origin. The Colombian *H. pylori* subpopulations represent an admixture of European, African and Native indigenous ancestry; although some genomes showed a high proportion of self-identity, suggesting a strong adaption to these mestizo Colombian groups. The Fst analysis identified 82 SNPs significantly fixed in 26 genes of the *hspColombia* subpopulation that encode mainly for outer membrane proteins and proteins involved in central metabolism. The strongest fixation indices were identified in genes encoding the membrane proteins HofC, HopE, FrpB-4 and Sialidase A. These findings demonstrate that *H. pylori* has evolved in Colombia to give rise to subpopulations following a geographical structure, evolving to an autochthonous genetic pool, drive by a positive selective pressure especially on genes encoding for outer membrane proteins.

## Introduction

The association of *H. pylori* with humans dates back at least 58,000 years, most probably since the origin of our species and accompanied humans during its migrations out of Africa.^1–3^ Genome analyzes estimates that East Asians populated the Americas about 23 kya ago and dispersed across the continent after a settlement in Beringia 8,000 years ago. ^4^ Thus, the first Indigenous Americans carried bacterial strains with Asian ancestry, which later evolved in isolation through a process of bacterial recombination and/or mutation. However, genome diversity of *H. pylori* in the Americas showed a drastic change after the arrival of Europeans and the transatlantic trade of African slaves starting 500 years ago, leading to genomes with a mosaic of admixed ancestries that have followed unique evolutionary paths.^5–7^

The Colombian population has a complex and diverse genetic structure as a result of migrations of different human groups separated by origin, time, and space.^8,9^ Cultural, geographical, and climatic characteristics, delimit the country in 6 regions, Andean, Pacific, Caribbean, Orinoquia, Amazonian and Insular. The human groups with African ancestry are concentrated in the Caribbean and Pacific regions, whereas the Indigenous people with Amerindian ancestry are found in the Amazon. People from the Andean region are mostly mestizo but with a high European ancestry component.^8^ Initial MLST studies of *H. pylori* in areas of high and low risk of gastric cancer (GC) in the country, identified strains with either a strong European (*hpEurope*) or an African *(hpAfrica*) ancestry. ^10–11^ Recent comparative genomic studies based on whole genomes analysis have revealed the presence of Colombian *H. pylori* groups that have evolved to contain a strong genome self-identity, giving rise to clearly differentiated Colombian subpopulations, although with a defined European ancestry.^15–18^ In order to better understand the population structure and admixture of *H. pylori* in Colombia, we analyzed the complete genome of 163 Colombian strains together with 1,113 publicly available genomes representing the already described worldwide subpopulations. In addition, we aimed to identify the gene variants that distinguished the Colombian strains from their parent population *hspSWEurope* by studying the single nucleotide polymorphism fixed in Colombia.

## Materials and methods

### *H. pylori* genomes studied

The 166 Colombian *H. pylori* genomes publicly available at Enterobase, Genbank (NCBI) and BIGSdb were included in this study, corresponding to strains isolated from patients residing in 10 departments of the country (supplemental figure 1). In addition, 1,143 *H. pylori* genomes from 46 countries were included to represent the different populations of *H. pylori* reported worldwide (supplemental table 1, supplemental figure 1). The 1,309 genomes were annotated with Prokka v1.12^19^ and then filtered according to genome features, discarding 33 strains with a genome length greater than 1.84 Mbp, or with more than 474 contigs, or a gene content greater than 1,696 genes. The final number of selected strains was 1,276 that included 163 from Colombia and 1,113 from other parts of the world (supplemental figures 1, 2 and 3, supplemental table 1).

### Analysis of *H. pylori* populations structure in Colombia

The Core-SNPs were called from the genomes with the Snippy program^20^ using the -contig option. The population structure was inferred from the 195,217 SNPs present in the core genome using FineSTRUCTURE v2 and ChromoPainter v2^21^ as previously described Muñoz-Ramirez et al., 2021.^7^ Population assignment of each sample was based on the observed clusters in the co-ancestry matrix and the population of 957 samples reported in previous works.^7,15–18^ To infer the ancestry proportions and identify admixture in Colombian *H. pylori* strains we conducted chromosome painting using ChromoPainter v2^22^, designating a collection of 1,276 Colombian and non-Colombian genomes as donors to paint each of the 163 Colombian recipient genomes.

### Whole-genome phylogenetic analysis

Whole genome phylogenetic relationship between all samples was analyzed with a Kmer-based method using the VAMPhyRE software, as described in Muñoz-Ramirez et al., 2017^16^, using a 13-mer probe set, with 4 bp probe extension and allowing one mismatch. Additionally, we made a phylogenetic reconstruction based on SNPs from the core genome. Then, phylogenetic trees were built using the neighbor joining method in MEGA X^23^ and visualized with iTOL v4.^24^

### Genetic diversity in Colombian subpopulations

To identify the genetic differences between the two *H. pylori* Colombian subpopulations identified in this study and their parental population *hspSWEurope*; first, we estimated the intrapopulation diversity calling the Core-SNPs of each subpopulation as described above, using as reference a representative strain of each group with high self-identity. Then we studied the interpopulation diversity, calculating the core genome in the three subpopulations using the 26695 strain as reference. The *hspSWEurope* group was built selecting 54 European strains to include samples from Spain, Portugal, France and Belgium, countries that were involved in the conquest of America and Colombia.

### Gene variants fixed in Colombia subpopulation

To identify genetic variants that characterize the *hspColombia* subpopulation we analyzed changes in allele frequency between strains from the *hspColombia* subpopulation and its ancestral population *hspSWEurope*. First, a core genome alignment of *hspColombia* and *hspSWEurope* strains was estimated using the Pangea pipeline^25^, then SNPs were extracted with the Snippy software and used as input to compute a fixation index (Fst) as implemented in VCFtools^26^. Fst values of each position in the core genome was visualized in R v4.0.5. The sequences of 26 core genes with Fst values >0.05 (table 1) were extracted and aligned by population using the muscle software included in MEGA X^23^. Consensus nucleotide and protein sequences were then aligned to the 26695 reference strain and visualized with WebLog^27^ to identify positions corresponding to synonymous and non-synonymous substitutions. Finally, the structure of proteins with at least two non-synonymous mutations were obtained from the Protein Data Bank^28^ when available, or predicted in the ITASSER server^29^, and visualized in the UCSF Chimera v1.5^30^ program, using a gradient color to highlight Fst values.

**Table 1.** Genes with a high FST value a position between *hspColombia* and *hspSWEurope.*

| Gene | Description | Number of SNPs | Number of SNPs with a non-synonymous effect | Number of codons in change | Fst max value | Location Protein | Biological Function |
| --- | --- | --- | --- | --- | --- | --- | --- |
| HP0486 | Outer membrane protein HofC | 19 | 10 | 7 | 0.937532 | Membrane | Survival |
| HP0706 | Outer membrane protein HopE | 9 | 8 | 6 | 0.86328 | Membrane | Survival |
| HP0554 | Sialidase A | 6 | 5 | 3 | 0.687286 | Membrane | Survival |
| HP1055 | Hypothetical protein | 6 | 3 | 3 | 0.717047 | Membrane | Survival |
| HP1512 | Iron-regulated outer membrane protein FrpB4 | 5 | 3 | 1 | 0.799812 | Membrane | Survival |
| HP0252 | Outer membrane protein HopF | 4 | 2 | 2 | 0.635204 | Membrane | Survival |
| HP0377 | Thiol:disulfide interchange protein | 4 | 3 | 2 | 0.674541 | Membrane | Survival |
| HP0517 | GTP-binding protein Era | 4 | 4 | 3 | 0.609185 | Cytoplasm | Central Metabolism |
| HP1487 | ABC-2 type transport system permease protein | 3 | 2 | 2 | 0.540672 | Membrane | Survival |
| HP0127 | Outer membrane protein HorB | 2 | 2 | 1 | 0.622084 | Membrane | Survival |
| HP0175 | Putative peptidyl-prolyl cis-trans isomerase PpiC | 2 | 0 | 0 | 0.641939 | Cytoplasm | Central Metabolism |
| HP0686 | Iron(III) dicitrate transport protein FecA | 2 | 1 | 1 | 0.54699 | Membrane | Survival |
| HP1565 | Penicillin-binding protein 2 | 2 | 2 | 2 | 0.584235 | Membrane | Survival |
| HP0018 | Hypothetical protein | 1 | 1 | 1 | 0.678227 | Unkown | Survival |
| HP0019 | Chemotaxis protein CheV | 1 | 1 | 1 | 0.66627 | Membrane | Survival |
| HP0130 | Hypothetical protein | 1 | 1 | 1 | 0.635371 | Unkown | Survival |
| HP0181 | Membrane protein required for colicin V production | 1 | 0 | 0 | 0.500663 | Membrane | Survival |
| HP0269 | (dimethylallyl) Adenosine tRNA methylthiotransferase | 1 | 0 | 0 | 0.504857 | Cytoplasm | Central Metabolism |
| HP0415 | Potassium efflux system protein/Small-conductance mechanosensitive channel | 1 | 1 | 1 | 0.50346 | Membrane | Survival |
| HP0558 | 3-oxoacyl-(acyl carrier protein) synthase II | 1 | 0 | 0 | 0.506588 | Cytoplasm | Central Metabolism |
| HP0564 | Uncharacterized protein | 1 | 0 | 0 | 0.625088 | Cytoplasm | Central Metabolism |
| HP0788 | Outer membrane protein HofF | 1 | 1 | 1 | 0.623103 | Membrane | Survival |
| HP1012 | Putative zinc protease PqqE | 1 | 1 | 1 | 0.589325 | Membrane | Survival |
| HP1054 | Hypothetical protein | 1 | 1 | 1 | 0.535516 | Membrane | Survival |
| HP1056 | Hypothetical protein | 1 | 0 | 0 | 0.511819 | Membrane | Survival |
| HP1340 | Biopolymer transport protein ExbD | 1 | 1 | 1 | 0.63672 | Membrane | Survival |

## Results

### Global and Colombian population structure of *H. pylori*

To learn more in detail the structure of the *H. pylori* populations in Colombia, a large number of genomes from worldwide populations were included in the analyses, particularly genomes from Spain, Portugal, and South Africa (countries involved in the migrations of the colonial period). A large number of *H. pylori* genomes from Colombia are publicly available (163) and all were included in the study.

The analysis of the population structure with FineSTRUCTURE for the 1,276 strains allowed the identification of 19 populations and subpopulations (figure 1). The genomes included in the analysis came from 46 countries, some of them poorly represented in previous studies, such as Australia and Papua New Guinea (supplemental table 1), which allowed the identification of new subpopulations (*hspNEuropeAustralia* and *hspSWEuropeAustralia*). The resulted co-ancestry matrix showed three main clusters, one mostly composed by Asian strains and includes the *hpAsia2, hpEAsia, hspIndigenous,* and *hpSahul*; *hpAfrica2* also felt within this large group. The second cluster was composed by African populations, including *hspAfrica1SAfrica*, *hspAfrica1WAfrica*, *hspAfrica1NAmerica* and *hspAfrica1Nicaragua*. The third and biggest cluster in the co-ancestry matrix was defined by populations with European ancestry and it can be divided in two subclusters, one composed by *hspSEurope*, *hspNEurope* and the novel subpopulation *hspNEuropeAustralia* not described previously and that included exclusively *H. pylori* genomes from Australia. The second subcluster was composed by the *hspSWEurope*, *hspSWEuropeHonduras, hspSWEuropeColombia* and the novel Colombian cluster named *hspColombia*. Within this subcluster felt what we previously described as subpopulations *hspSWEuropeMexico* and *hspAfrica1MiscAmerica*^7,17^, but now reviewed after extending the study to include more strains from Latin America we renamed these two last groups as *hspSWEuropeNorthAmerica* and *hspSWEuropeSouthAmerica* (figure 1). Furthermore, within this subcluster we also identified a new Australian subpopulations not described before, that we named *hspSWEuropeAustralia* (figure 1).

**Figure 1.**
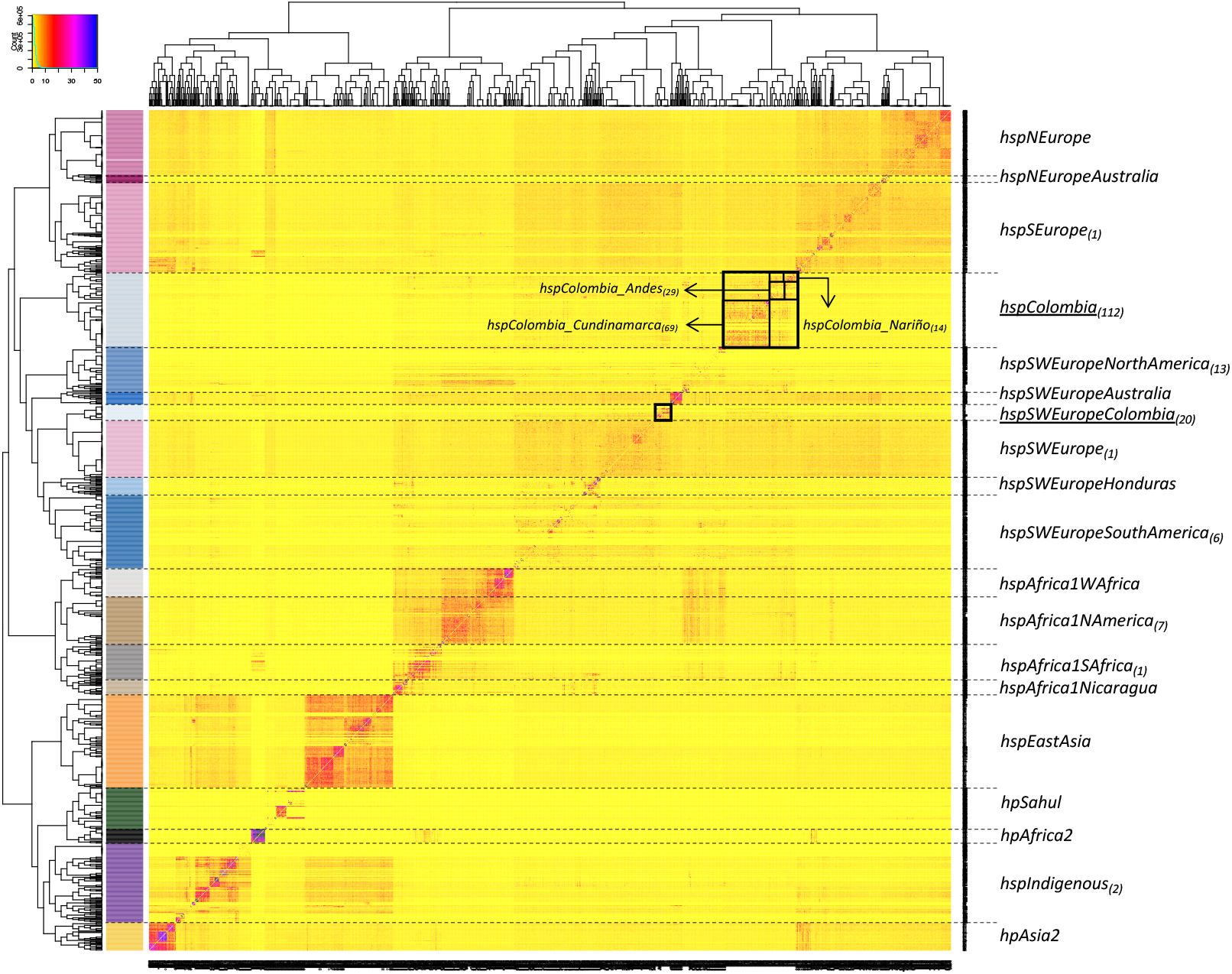
Co-ancestry matrix showing the population structure and genetic flow of the 1276 *H. pylori* strains analyzed. The colorimetric gradient in the heat map corresponds to the number of genomic motifs imported from a donor genome (column) to a recipient genome (row). The inferred tree is displayed at the top and left of the heat map, and *H. pylori* strain names are at the bottom and right. The allocation of the *H. pylori* population is provided in colors on the left side of the heat map and the names of each population group are located on the right; the subscripts indicate the number of Colombian strains identified in each population. The black boxes indicate the subpopulations identified in the majority in Colombia and in the case of *hspColombia*, the presence of three possible subpopulations within it is highlighted.

The two Colombian subpopulations were clearly separated (figure 1, arrows), the previously reported as *hspSWEuropeColombia* (because of its closeness with *hspSWEurope* strains) by Muñoz et al. (2017)^17^ and the one not previously described that we designated as *hspColombia*. This last population was further separated in three subclusters, which corresponded to their region of origin and were named as *hspColombia_Nariño* that included only isolates from the Nariño department; *hspColombia_Andes* composed mainly with strains from departments of the Andes Mountain region and *hspColombia_Cundinamarca* comprised mostly by strains isolated in the Cundinamarca department (supplemental figure 4). Most of the Colombian strains (81%) belonged to either *hspSWEuropeColombia* or to *hspColombia* and the remaining belonged to the other 16 subpopulations, particularly to *hspSWEuropeNAmerica*, *hspSWEuropeSAmerica* or *hspAfrica1NAmerica* (supplemental table 2).

### Phylogenetic analysis of all worldwide strains

To further support the above population structure results we made a whole genome phylogenetic analysis based on SNPs from the core genome of the 1,276 strains. The analysis showed clustering of *H. pylori*, which closely corresponded to the populations and subpopulations defined by FineStructure (figure 2A). Well defined clusters were identified for Asian population, for European from old-world and Latin America, and finally African old world and Latin America. (figure 2A). Also, it was noticed that Colombian subpopulations grouped in at least three different clusters, clearly separating the previously described *hspSWEuropeColombia* from the novel Colombian subpopulations identified in this study (figure 2). To further investigate the phylogenetic distribution within Colombian groups, we did a phylogenetic tree including only Colombian strains and identified at least tree clades, one composed by isolates from Cundinamarca department, another by isolates from Nariño and Andes departments, and the last made by isolates belonging to *hspSWEuropeColombia* (figure 2B).

**Figure 2.**
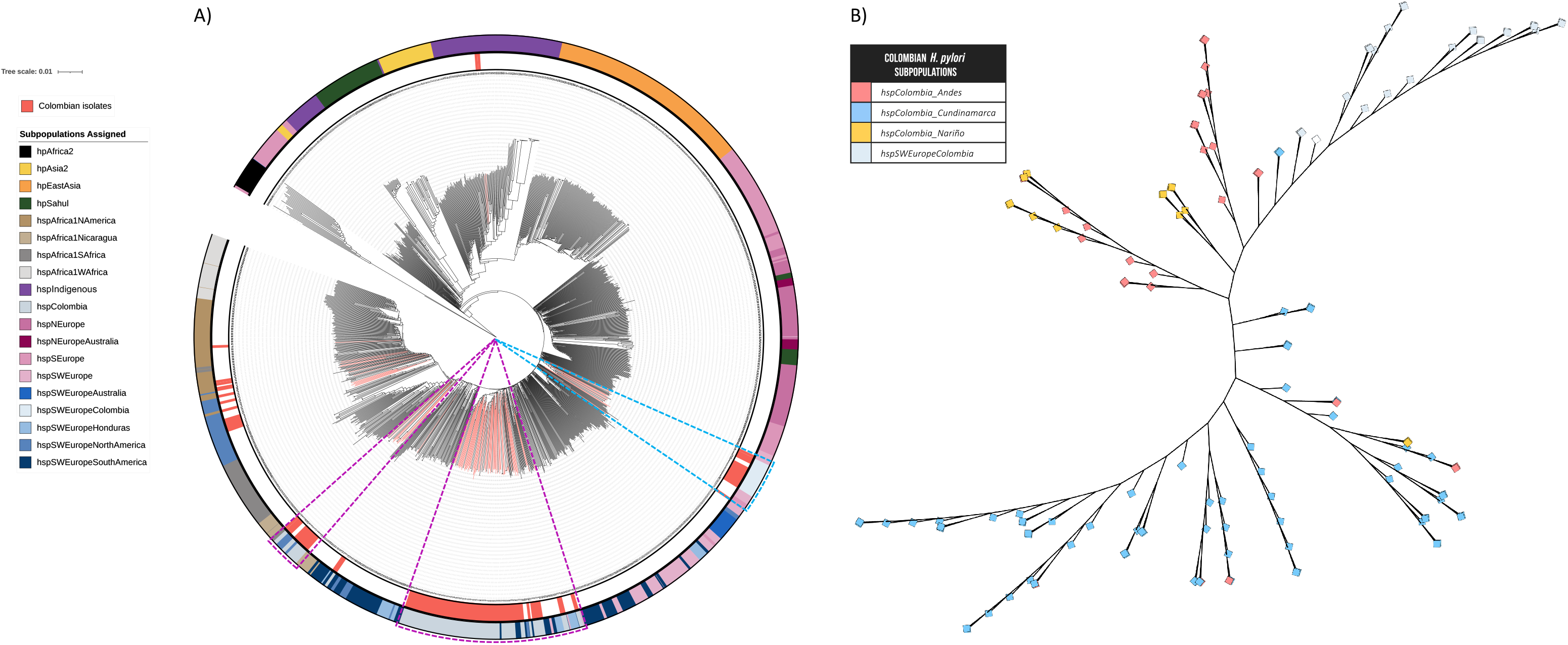
Phylogenetic analysis of the 1276 *H. pylori* strains analyzed based on. A) SNPS of the core genome. The phylogenetic analysis was performed with MEGA X, the phylogenetic tree was calculated using the Neighbor-joining model and it was edited in iTOL 4. The fuchsia color of the tree branches indicates the Colombian isolates included. The internal ring and colored branchs represent the *H. pylori* isolates from Colombian; the colors of the outer ring represent the *H. pylori* subpopulations assigned in the population structure analysis. B) Phylogenetic analyses including only the four subpopulation of Colombia.

### Admixture and diversity in Colombian subpopulations of *H. pylori*

A high proportion of the genomes of the old-world populations *hpAfrica2, hspEastAsia, hpSahul,* and *hspNEuropeAustralia* are painted by their own population (self-identity). In contrast, most populations with European ancestry (*hspNEurope, hspSEurope*, and *hspSWEurope*) present high admixture, contributed by different populations (figure 3). In the Colombian subpopulations, contribution from European, African and Amerindian donors was observed, but self-identity was the predominant ancestral component in some of the strains (supplemental figure 5A). Of note, a number of genomes of isolates from *hspColombia_Nariño, hspColombia_Andes*, and *hspColombia_Cundinamarca* were completely painted by their own subpopulations showing almost 100% of self-identity (supplemental figure 5B).

**Figure 3.**
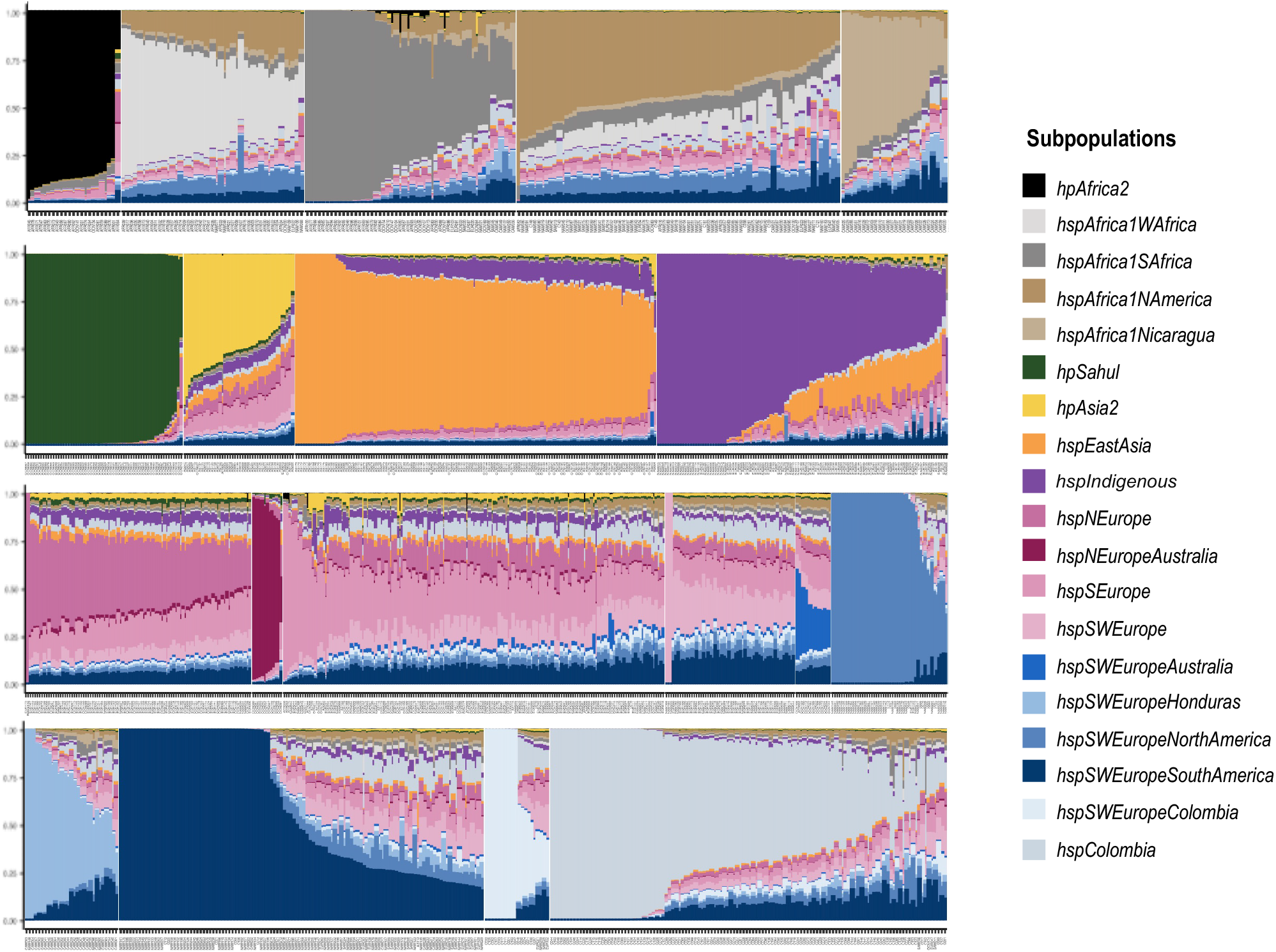
Representation by subpopulation of the mix of ancestry in *H. pylori* strains from populations analyzed using ChromoPainter version 2. From left to right, African, Asian, Amerind, European, and Colombian subpopulations. Each column represents a strain, and the color indicates the proportion of ancestry in that genome. The color code for each subpopulation is shown to the right of the figure.

To further analyze diversity in the genomes of the subpopulations we estimated the number or SNPs in relation to the number of analyzed genomes (supplemental figure 6). Genomes from *hspSWEurope* were more diverse, showing a larger number of SNPs followed by *hspSWEuropeColombia* and then by *hspColombia*.

### Genes coding for outer membrane proteins are significantly fixed in the *hspColombia* subpopulation

We analyzed 196,360 SNPs present in the core genome to identify nucleotide positions that have become significantly fixed in Colombian subpopulations when compared with their ancestral *hspSWEurope* subpopulation; the distribution of Fst values over the core genome is illustrated in figure 4. By selecting a Fst value over 0.5 we identified 26 genes significantly fixed in Colombian subpopulations, most of which encode for outer membrane proteins with a bacterial survival function (figure 4 and table 1). A total of 82 sites were identified in these 26 genes, 53 of these SNPs generated non-synonymous substitutions, some of these occuring in the same codon (supplemental table 3). The higher number of fixed positions and non-synonymous changes were identified in genes that encoded for the outer membrane proteins HofC (10 sites) and HopE (8 sites) as well as for Sialidase A (5 sites). In these 26 genes, there was a high number of outer membrane and transmembrane transport proteins involved in the interchange of ions and macromolecules, adhesion, bacterial adaptation to variation in the microenvironment, chemotaxis and homeostasis.^31,32^ We identified the position of each significant NS mutation in the seven genes with the highest Fst values, that encodes the membrane proteins HofF, Thiol:disulfide interchange protein, HofC, sialidase A, HopE, an hypothetical protein, and FrpB4 (figure 5).

**Figure 4.**
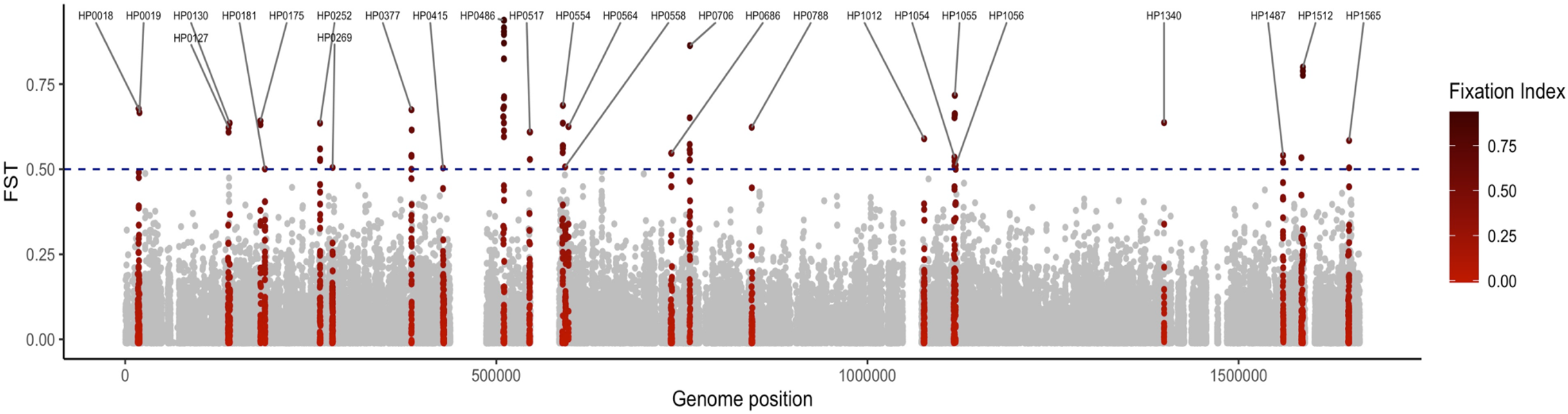
Core-genome Fst analysis to identify genetic variants that are significantly more comon in the *hspColombia* than in its parental subpopulation *hspSWEurope*. The *X*-axis indicates the nucleotide sites in the genome and the *Y-*axis shows the Fst value for each site. In red was show the 26 genes that present a site with an FST value of 0.5 or greater.

**Figure 5.**
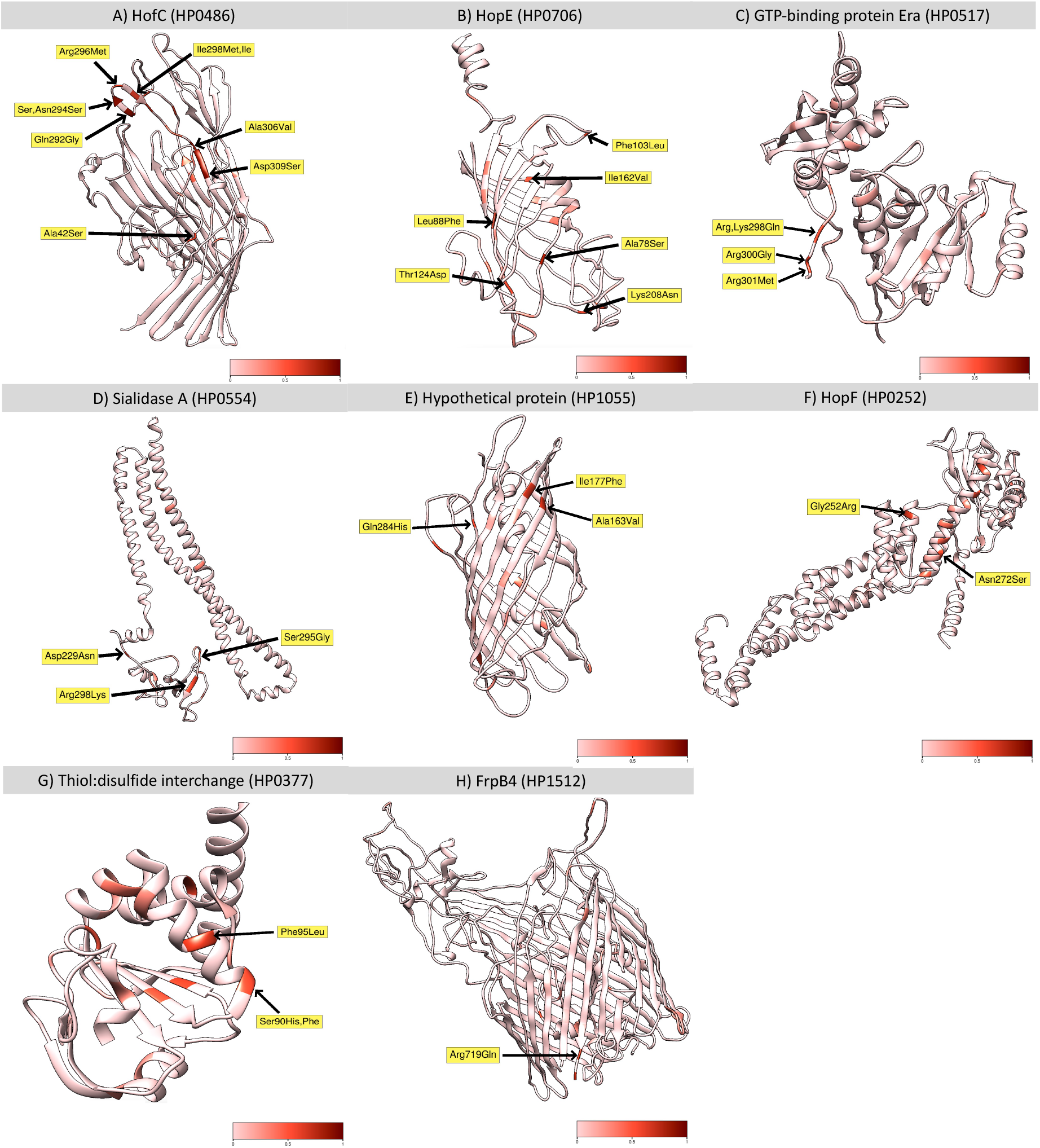
Location of non-synonymous mutations with a significative Fst value in the proteins with the highest number of variant positions. The 3D structure of each protein was inferred in TASSER server. A) HofC (HP0486). B) HopE (HP0706). C) GTP-binding protein Era (HP0517). D) Sialidase A (HP0554). E) Hypothetical protein (HP1055). F) HopF (HP0252). G) Thiol:disulfide interchange (HP0252). H) FrpB4 (HP1512). The color scale represents the Fst value for each position.

In the gene that encodes for HofC, an outer membrane protein involved in the adhesion of the bacteria to gastric epithelial cells,^31,32^ up to 19 fixed positions were identified, of which 10 were non-synonymous with Fst values higher than 0.94. Eight of these SNPs were concentrated in the 858–918 nucleotide positions, which correspond to the 291–309 aa residues of the protein (figure 5A); four of these variants generate changes in acid or basic aa to neutral leading to a change in net charge (figure 5A, supplemental figure 7A). The second gene with more changes was *hopE* that encodes for a porin involved in the adhesion to gastric epithelium, and which may induce an immune response.^33^ Highly significant nonsynonymous changes were identified in the 88–208 aa residues (figure 5B, supplemental figure 7B). Strong Fst values were found in the GTP-binding Era protein, an essential GTPase that binds both GDP and GTP, with 4 significant Fst values in the 298 – 301 aa residues (figure 5C, supplemental figure 7C). For the Sialidase A and the hypothetical protein HP1055, the nonsynonymous changes were also limited to a specific region of the protein (figure 5D and E, supplemental figure 7D and E), but little is known about the function of these proteins. We also found changes in metabolic proteins associated with protein folding, cell cycle regulation, and energy metabolism (figure 5G and H, supplemental figure 7G and H).

## Discussion

### A new Colombian *H. pylori* subpopulation

Previous studies have shown the rapid and ongoing evolution of *H. pylori* populations in the Americas since the encounter of European, African and Native Indigenous (Amerind) human populations 500 years ago.^4^ Studies have found an exquisite adaption of *H. pylori* to the different emerging mestizo human populations, distinguishing even between neighbor countries.^7,17,18^ In Colombia a subpopulation with European ancestry but yet a high proportion of self-identity have been confirmed by several groups (*hspSWEuropeColombia*) and recent reports suggest there might be more than one *H. pylori* subpopulation within Colombia.^8,11–13,15–18^ It has been reported that even within countries there might be different subpopulations driven by geographical separation, diet, costumes, among others. ^7,13–18^ In Colombia there have been important efforts to sequence the genome of a significant number of *H. pylori* isolates from different regions of the country. We took advantage of this to perform a comprehensive study of *H. pylor*i subpopulations by analyzing the genome of isolates from different regions of the country. The inclusion of a large number of genomes from all over the world allowed us an improved resolution to identify unknown subpopulations in Colombia but also in other countries (supplemental figures 1 and 2, supplemental table 1). We redefine the structure of isolates from the previously named *hspSWEuropeMexico* and *hspAfrica1MiscAmerica* and identified four new subpopulations (figure 1) composed of isolates from Latin America (*hspSWEuropeNorthAmerica* and *hspSWEuropeSouthAmerica*) and Australia (*hspNEuropeAustralia* and *hspSWEuropeAustralia*).

Recent studies have documented a detailed adaptive process of *H. pylori* to different mestizo populations in Latin American countries such as Mexico, Honduras, Nicaragua, and Peru.^5,7,15–18^ We show the presence of four subpopulations inside Colombia, supported by a comprehensive genomic analysis of a considerable number of *H. pylori* strains from different regions of the country; such a detailed geographic substructuring of subpopulations inside a country has not been reported.

The population structure analysis revealed the existence of two clearly separated subpopulation within the Colombian isolates (figure 1), one corresponded to the previously described as *hspSWEuropeColombia*^7^, grouping close to *hspSWEurope*, whereas the other was a distant cluster we designated as *hspColombia*. We were able to identify well-differentiated clades of Colombian strains, with a resolution in geographical regions not previously described. Our approach was to perform whole-genome analyses; whereas previous studies were limited to MLST analysis or included a reduced number of genomes.^11–13,15–18^ Our phylogenetic analysis identified different groups, suggesting the presence of more than one evolutionary lineage, as previously observed by Gutiérrez et al., (2017)^15^ in Colombian isolates from the Cundinamarca department. Furthermore, a detailed analyses indicated the presence of three subpopulations within *hspColombia*, which we named after their geographic origin as *hspColombia_Nariño*, *hspColombia_Andes* and *hspColombia_Cundinamarca*, all isolates from departments in the Andes Mountains such us Nariño, Boyacá, Santander, Tolima, and Cundinamarca. These four subpopulations were within a larger group including Latin American clusters with European ancestry, particularly close to *hspSWEurope* (figure 1 and 2B).

In addition to the above subpopulations, a few other Colombian strains felt into the African subpopulation *hspAfrica1NAmerica*, isolated from patients in the coastal region of the Nariño department that matches the African ancestry of the human population, descendent from the African slaves that established during 16th to the 19th century.^34,35^ Although only 2 strains from Amazonian inhabitants were included, they classified within the *hspIndigenous* subpopulation previously reported in native communities of Latin America.^7,36^ Thus, Colombia is a complex mosaic of *H. pylori* subpopulations with European, African or Indigenous ancestry that mirror the complex array of human groups distributed along the country.^8^ This study supports the existence of an evolutive differentiation process, with new subpopulations within the country that are evolving to adapt to the Colombian mestizo population from the different geographical regions.

High admixture is commonly observed in *H. pylori* genomes of European ancestry, even in the old-world regions (see figure 3) must probably because of the constant movement of people between countries. This high admixture is also observed in genomes of the Americas with either African or European ancestry, reflecting that these populations are the result of the “recent” encounter of European, African and Indigenous human races 500 years ago. Still, it is remarkable to observe that several *H. pylori* genomes of the American subpopulations with European ancestry (*hspSWEuropeNorthAmerica*, *hspSWEuropeSouthAmerica*, *hspSWEuropeColombia* and *hspColombia*) show a reduced admixture and are mostly *painted* by their own population (self-identity ancestry), suggesting an advance process of adaption to their human host. This “rapid” evolution may be the result of rampant genetic exchange in populations with high transmission rates like Latino America^18^. This study shows evidence documenting an advance process of adaptive evolution that distinguish between subpopulations in different regions of Colombia, partially explained by the presence of environmental and cultural conditions specific to each Andin department.^34,35,37,38^

The difference between Colombian *H. pylori* subpopulations was further documented with a genomic diversity analysis showing that *hspSWEuropeColombia* had significantly higher number of SNPs per genome than *hspColombia*, probably because diversification of *hspSWEuropeColombia* started earlier. This genomic diversity supports the existence of different evolutive lineages in Colombian subpopulations, as previously suggested by Gutierrez et al., (2017).^15^ Also, diversity of Colombian subpopulations was lower than *hspSWEurope*, which is expected considering that *hspSWEurope* is the parental population.

### The membrane proteins characterized a Colombian *H. pylori* subpopulation

The Fst analysis allowed the identification of the genes that contributed the most to the differentiation of the Colombian subpopulations and identified genes that encoded for proteins involved in bacterial survival and central metabolism. A large number of SNP positions were identified in membrane proteins associated with transport, structure, adhesion, and homeostasis, which play an essential role in the interaction and adaptation to the host.^31,32^ These results suggest that the changes observed in host-interaction proteins play a major role in differentiation of *H. pylori* populations in Colombia, which is in agreement with recent reports in other Latin American subpopulations.^7^ In fact, 10 of the proteins with significant Fst values coincide in both studies, including the outer membrane proteins *hofC*, *hopC*, the transport protein *fecA* and the isomerase *ppiC*. Still, there were outer membrane genes like *frpB4* and *hopF* with high Fst values for Colombian populations that were not reported in the Latin American study, suggesting the existence of specific adaption for some populations.

The *hofC* gene presented the largest number of positions with significant Fst values, most of them localized in the 858–918 nucleotide region, which agrees with the report by Thorell et al., (2017)^18^ that also found several SNPs in *hofC* in Latin American *H. pylori* subpopulations. Many of the identified SNPs cause amino acid changes modifying polarity, charge, or pH, which could modify the structure and the performance of the protein.

The outer membrane proteins HofC and HopC may function as adhesins recognizing receptors on the surface of the gastric epithelial cells and both human receptor and *H. pylori* adhesin may have co-evolved to have the right interaction. It is less clear for FecA, that transport ferric citrate or for the isomerase PpiC; probably they also interact with human proteins. We also found changes in proteins that regulate concentration of iron and potassium. The causal mechanism underlying this association is not clear, but it plausibly reflects a bacterial response to high concentrations of ions in the water and food due to Colombian geology, especially in the mountain areas of the country.^35^ The interaction of these proteins with proteins of the host or the environment of the gastric mucosa exert a strong positive selection to attain the right adaption of the bacterial-host interaction. In genes like those coding for outer membrane proteins HofC and HopE, fixed SNPs occur in more than one position (table 1 and figure 5) suggesting that in some genes mutations in multiple sites are needed to get the right changes in regions of the protein to adapt to the human population. In this study, the outer membrane protein genes *hofC*, *hopE* and *frpB4* had the highest Fst values identified, indicating that they play a major role in the adaptation of *H. pylori* to the Colombian human population, and are under strong selective pressure to drive the evolutive process during the differentiation of the populations.

## Conclusions

The findings described in this study demonstrate that *H. pylori* have evolved in Colombia to give rise to new subpopulations that follow a geographic structure. The evidence suggests that this evolutive process is in progress, with some strains having already a genome with almost 100% self-identity but others with gradients of admixture. The study also show that outer membrane protein genes present the highest fixation values and seem to drive the evolutive process as a result of high selective pressure by the host.

## Competing interests

None declared. The authors declare that they have no competing interests related to the subject matter or materials discussed in this article.

## Funding

Mabel Bohórquez and Magdalena Echeverry received research funding from Tolima University (Projects 30113, 350113, 160114, 10110, 40218 - contracts 398-2017 and 851-2020); COLCIENCIAS (grant number 110565843382; contract 204 −2015). Javier Torres received funding from the Infectious Diseases Research Unit of the Mexican Institute of Social Security. Luis Carvajal-Carmona receives funding from the National Cancer Institute (grants R01CA223978 and P30CA093373) of the US National Institutes of Health and The Auburn Community Cancer Endowed Chair in Basic Science. Alix A. Guevara-Tique and John J. Suaréz Received funding from the high level of human capital formation program of Tolima Department, Colciencias, and Tolima governorate (755-2016).

The content of this article is solely the responsibility of the authors and does not necessarily represent the official views of the National Institutes of Health or any of the funding agencies.

## Acknowledgments

To the researchers who participated in this study, to the Cytogenetics, Phylogeny, and Population Evolution research group from Tolima University and to the National Cancerological Institute, Bogotá, Colombia.

## Notes

### Competing Interest Statement

The authors have declared no competing interest.

