## Supplemental figures for "Selective pressure on membrane proteins drives the evolution of *Helicobacter pylori* Colombian subpopulations"

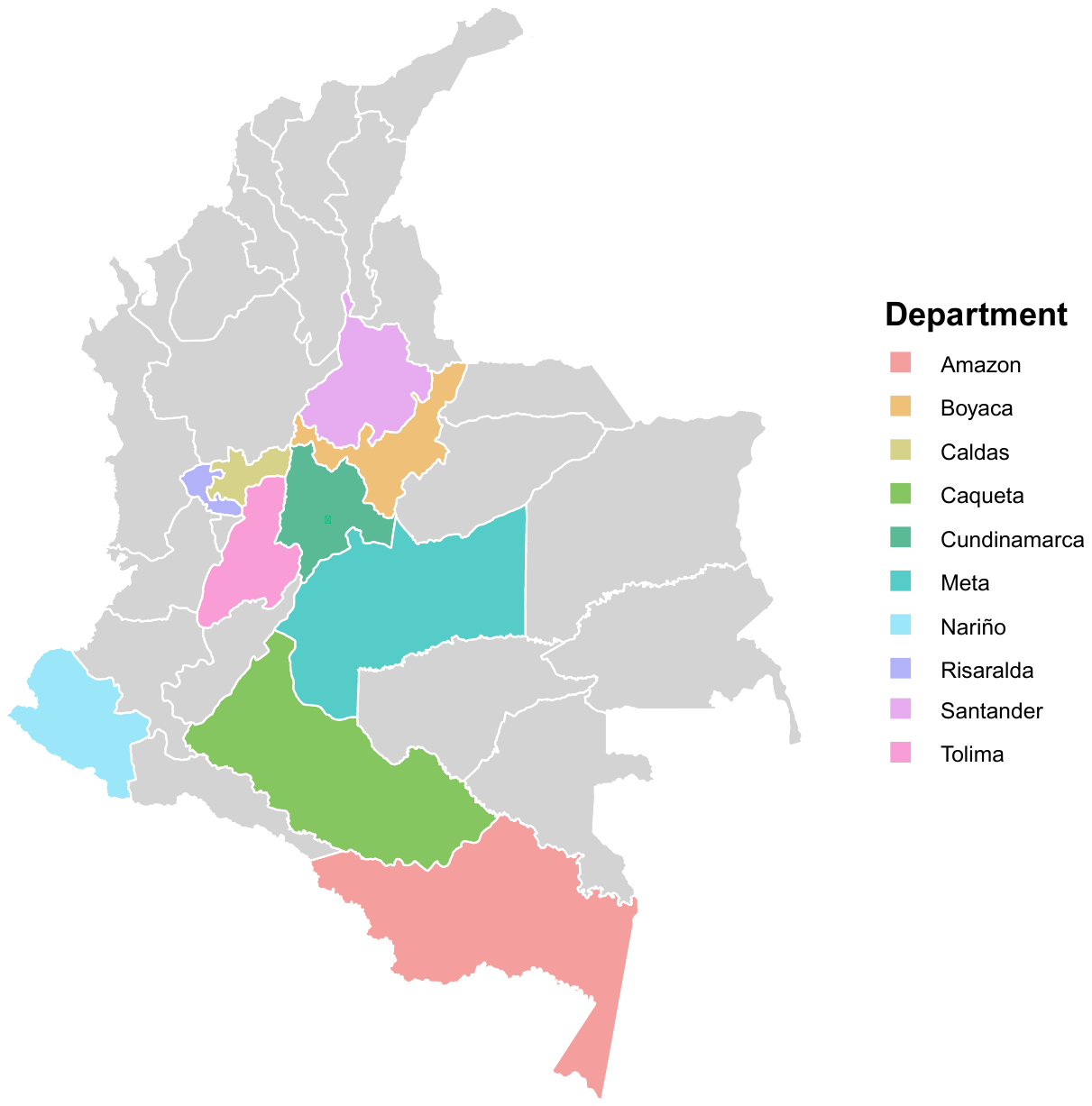

**Supplemental Figure 1.** **Geographical origin of Colombian *H. pylori* strains included in the analysis.**

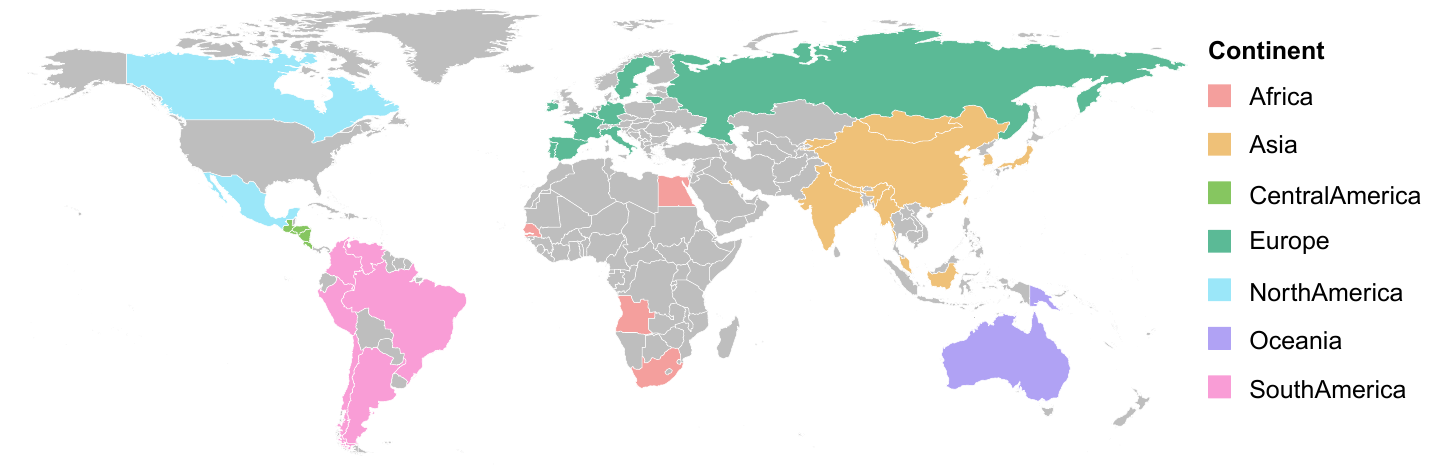

**Supplemental Figure 2. Geographical origin of the 1276 strains included in the analysis.**

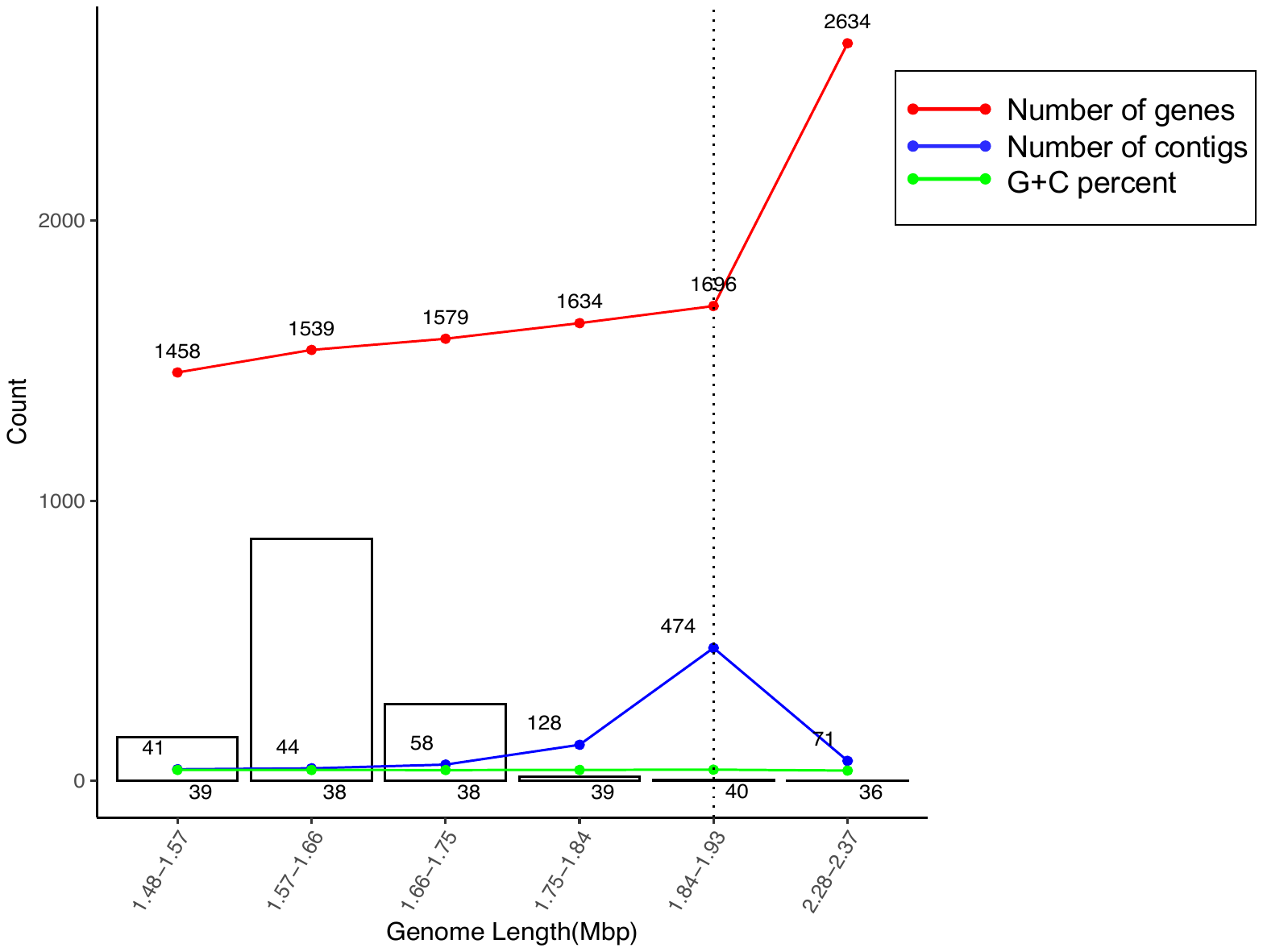

**Supplemental Figure 3. Quality control of the analyzed strains.** The X axis shows the size of the genome and the Y axis the number of strains analyzed; the red line indicates the number of genes identified in the genomic annotation; the blue, the number of contigs, and the green the percentage G + C of the strains analyzed. The figure was made in R program version 3.5. (Alix, yo quitaria esta figura; es medio confuse y no ayuda mucho)

**Supplemental Figure 4**. **Geographical origin and ancestry of the Colombian *H. pylori* subpopulations.** The background color of the map shows the majority ancestry component reported in the human population in those natural regions. The circles represent the places where there are *H. pylori* isolates that have had the complete genome sequenced, and the colors inside each circular graph reflect the subpopulations identified in the isolates from each department.

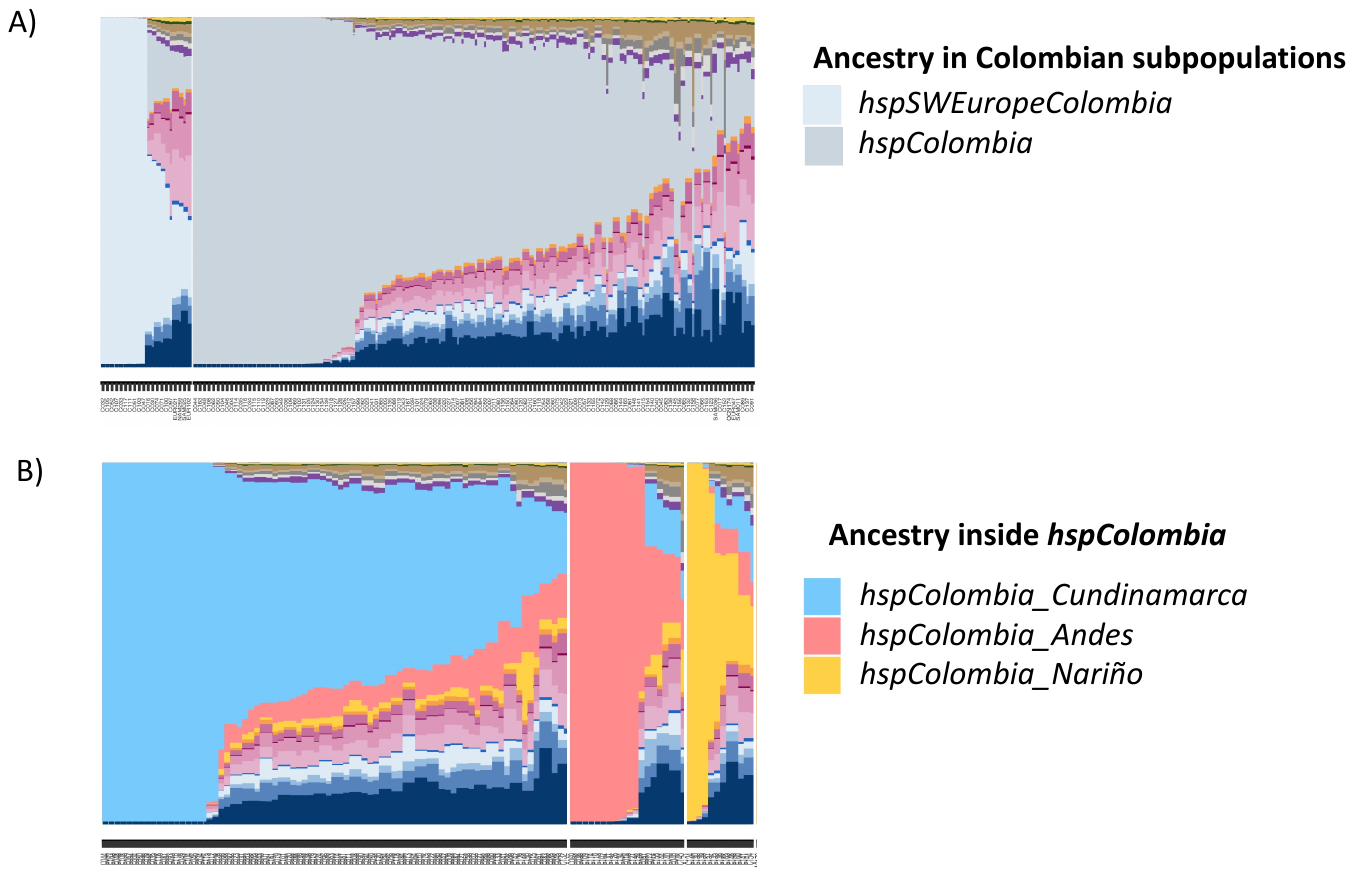

**Supplemental Figure 5.** **Representation of the admixture in Colombian subpopulations.** A). Admixture in the two large *H. pylori* subpopulations using all-world genomes as donors. B). Admixture of subpopulations within *hspColombia*. The samples were analyzed using ChromoPainter version 2. Each column represents a strain, and the color indicates the proportion of ancestry in that genome. The color code for each subpopulation is shown to the right of the figure.

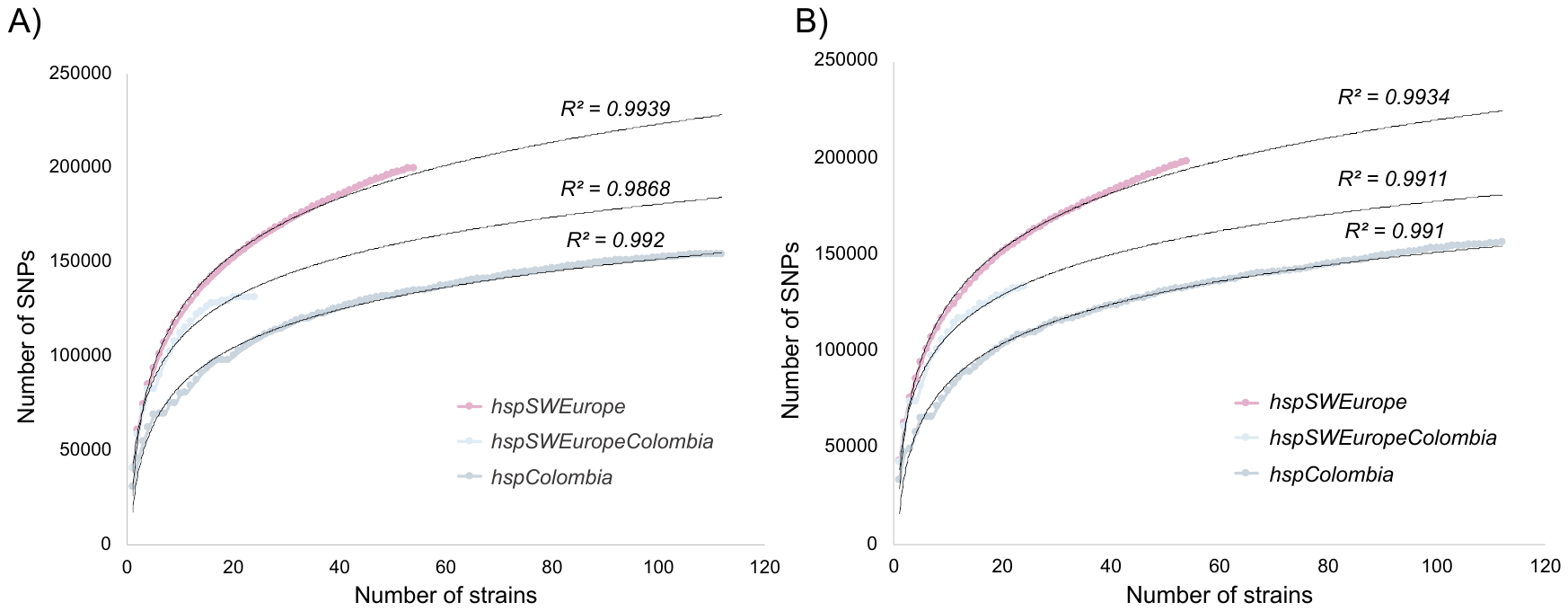

**Supplemental Figure 6. Genetic diversity in Colombian and *hspSWEurope* subpopulation.** A). Intraspecific diversify. B) Interspecific diversity. The X axis shows the number of strains analyzed by populations and the Y axis the number of accumulated SNPs; the pink line corresponds to isolates from *hspSWEurope*, the light blue to *hspSWEuropeColombia* and the light gray to *hspColombia*.

A) HofC (HP0486)

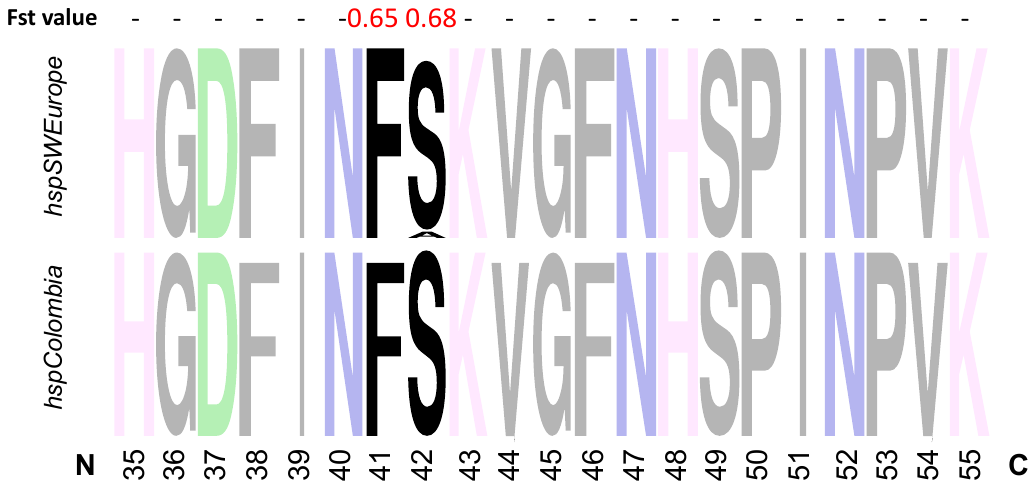

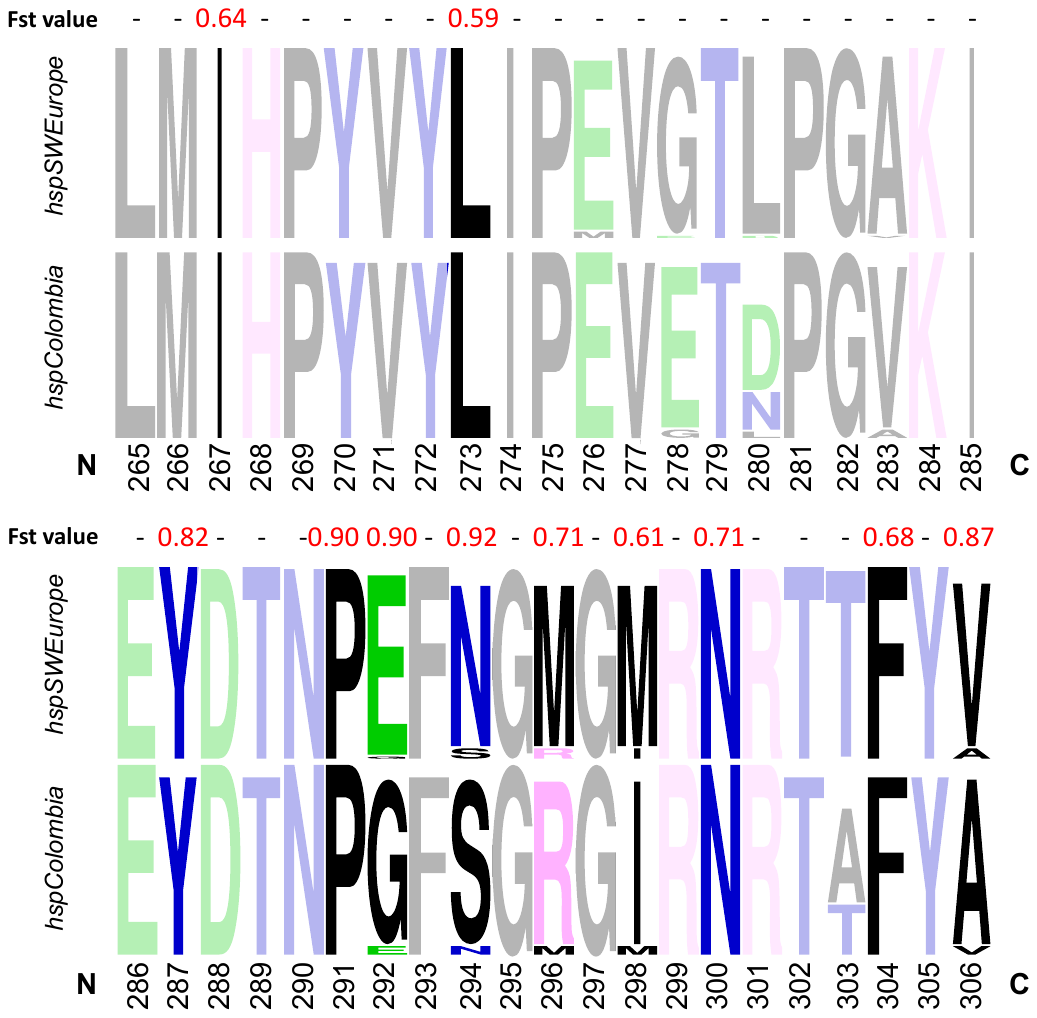

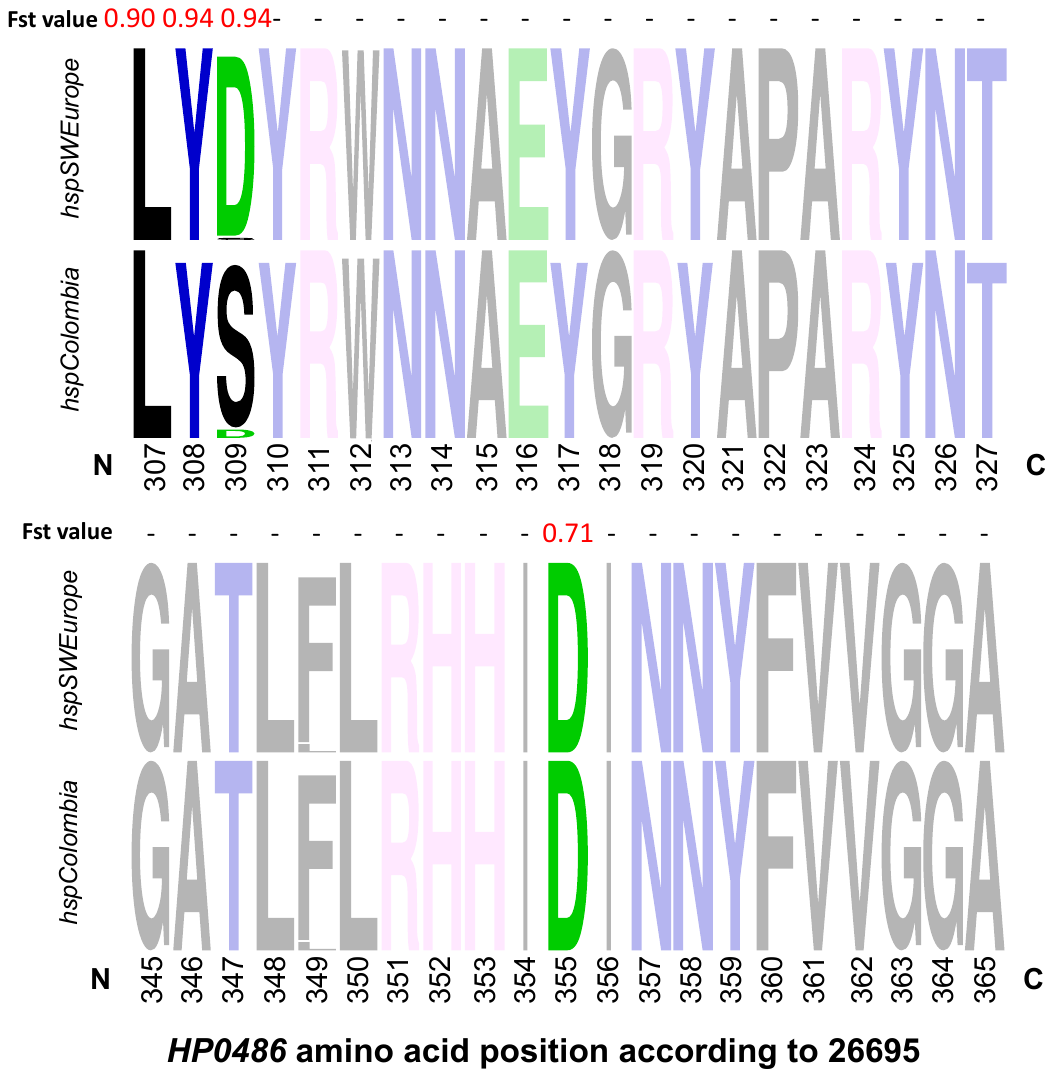

B) HopE (HP0706)

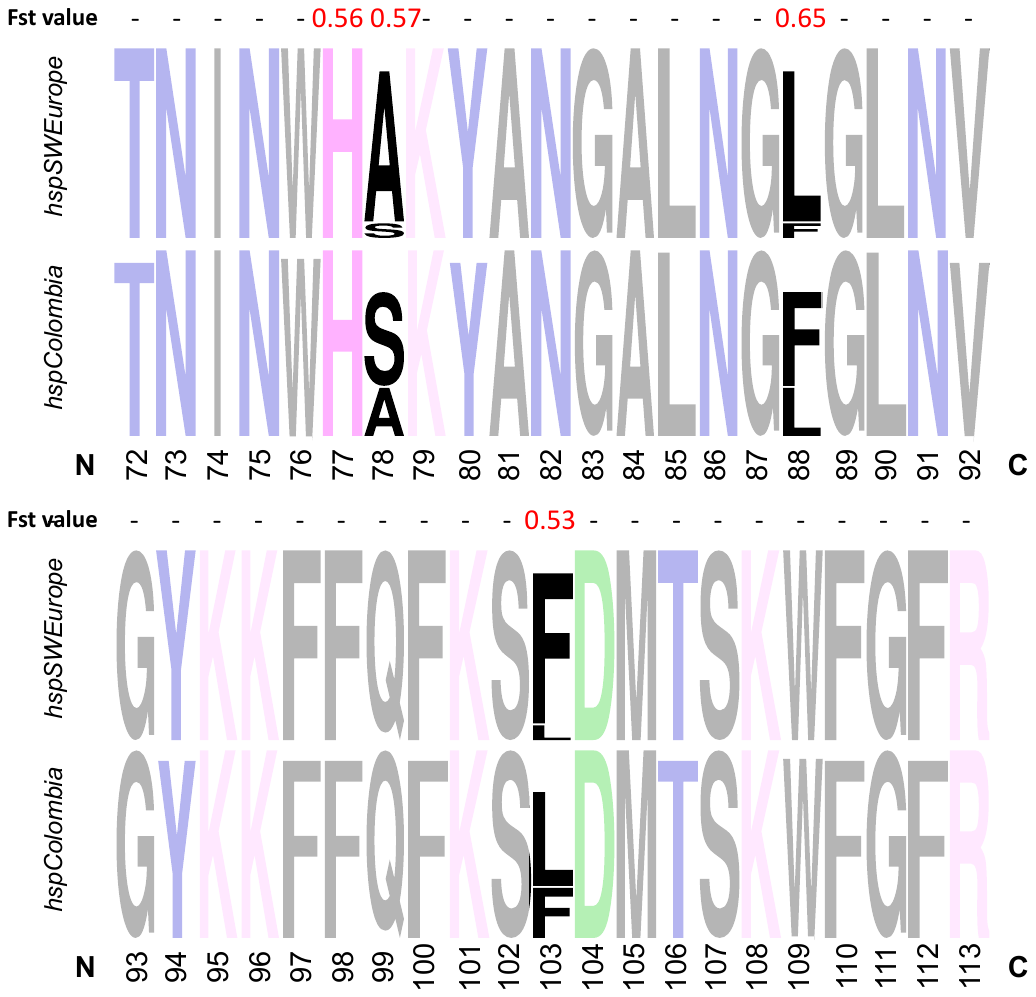

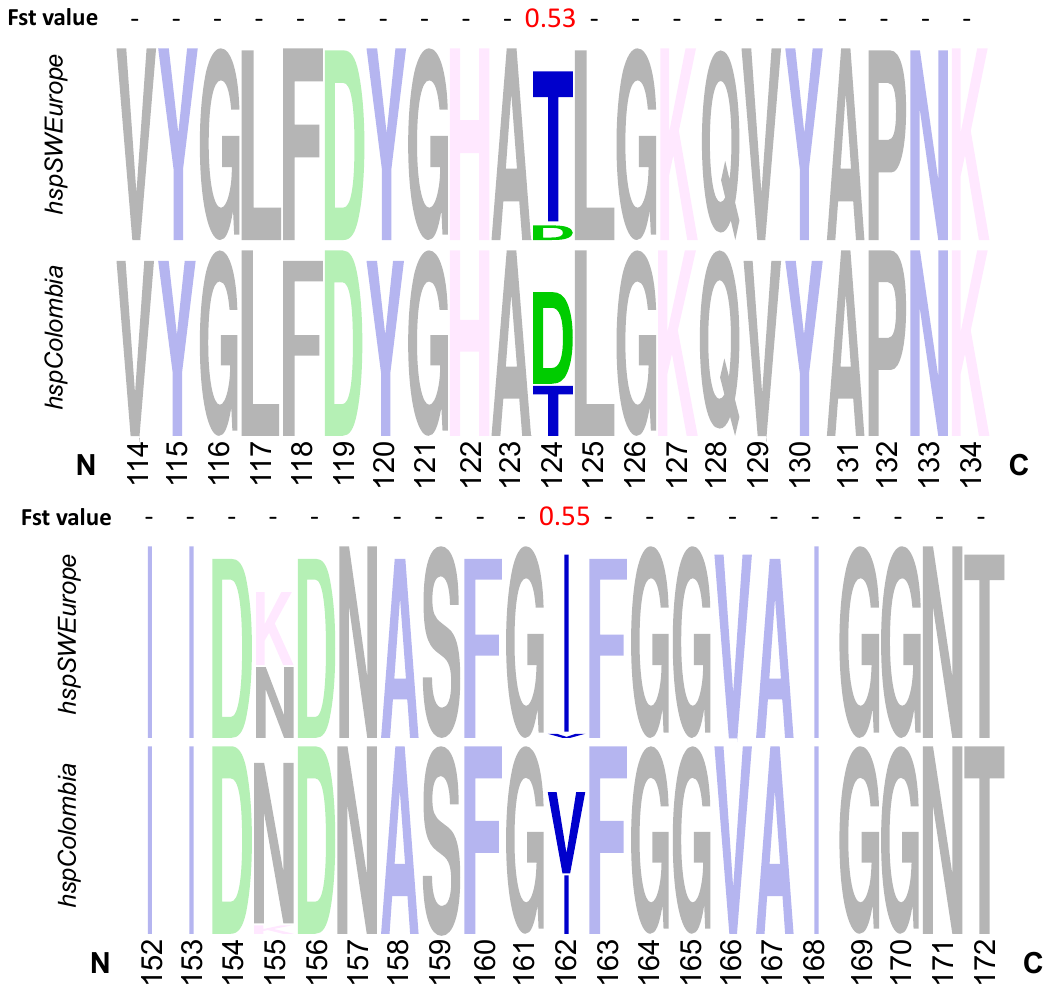

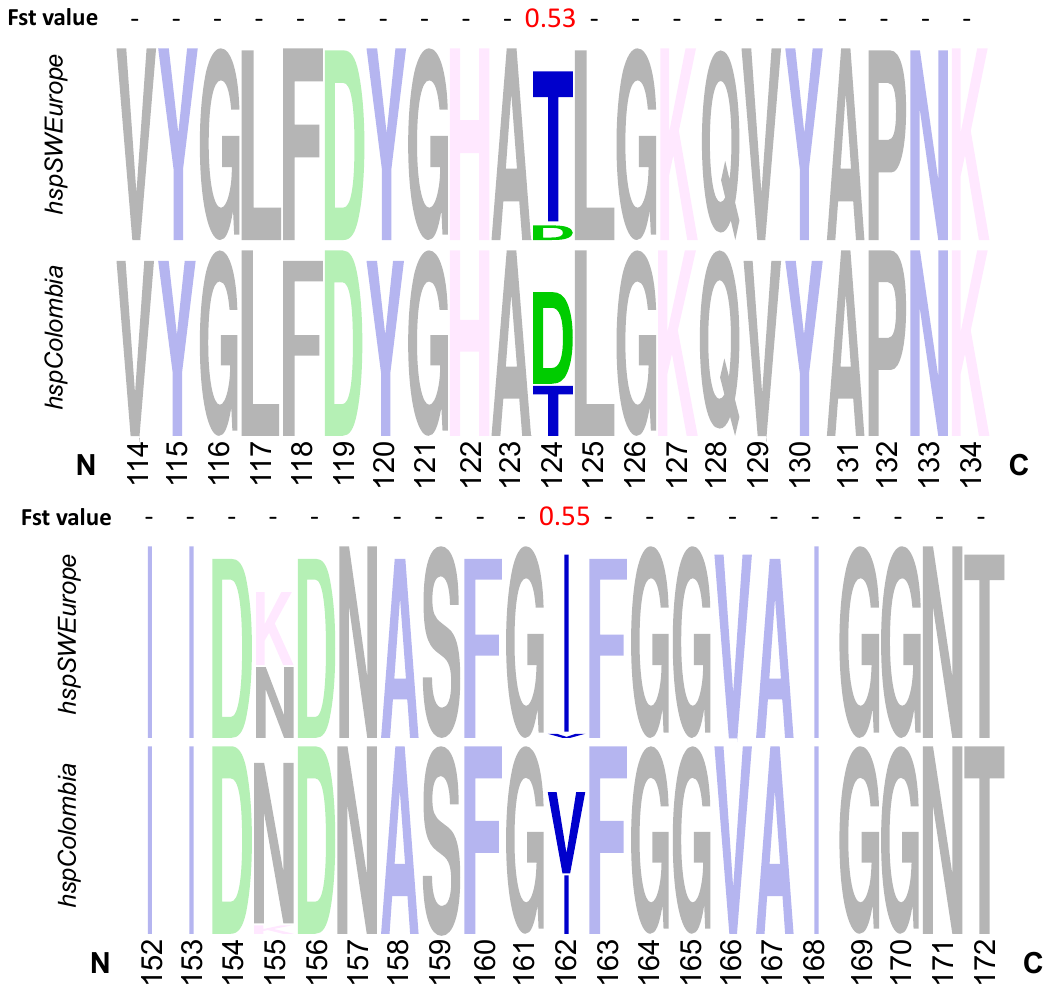

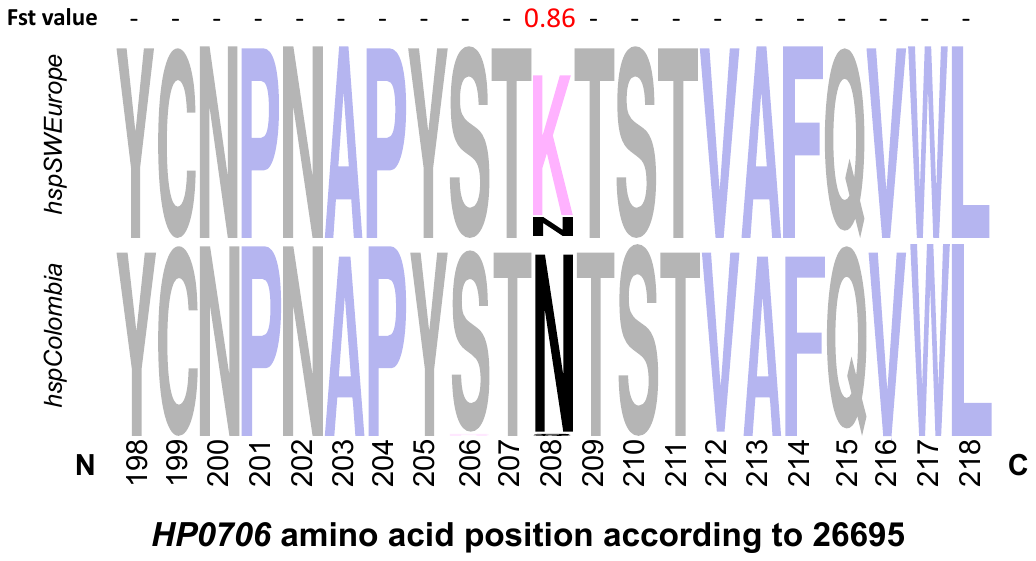

C) GTP-binding protein Era (HP0517)

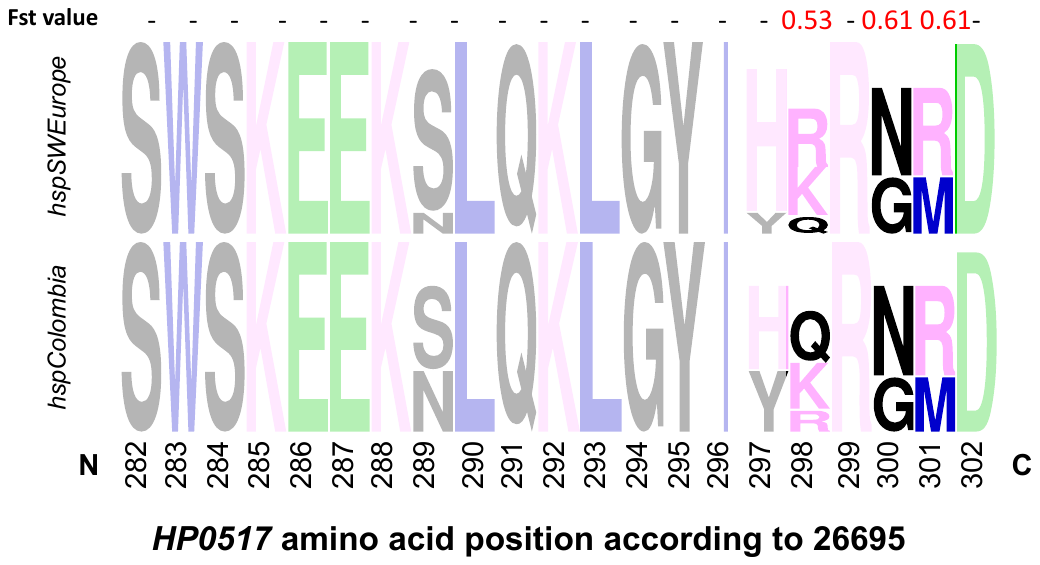

D) Sialidase A (HP0554)

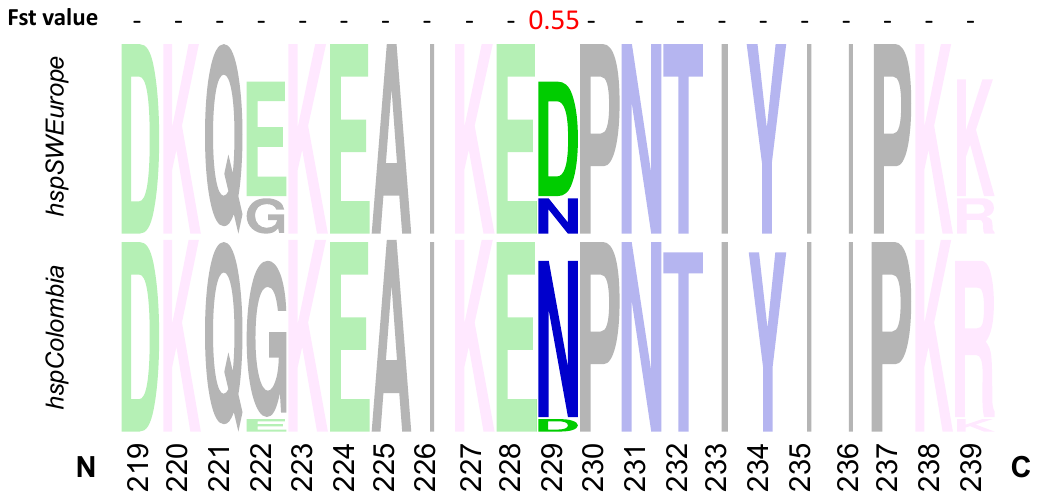

E) Hypothetical protein (HP1055)

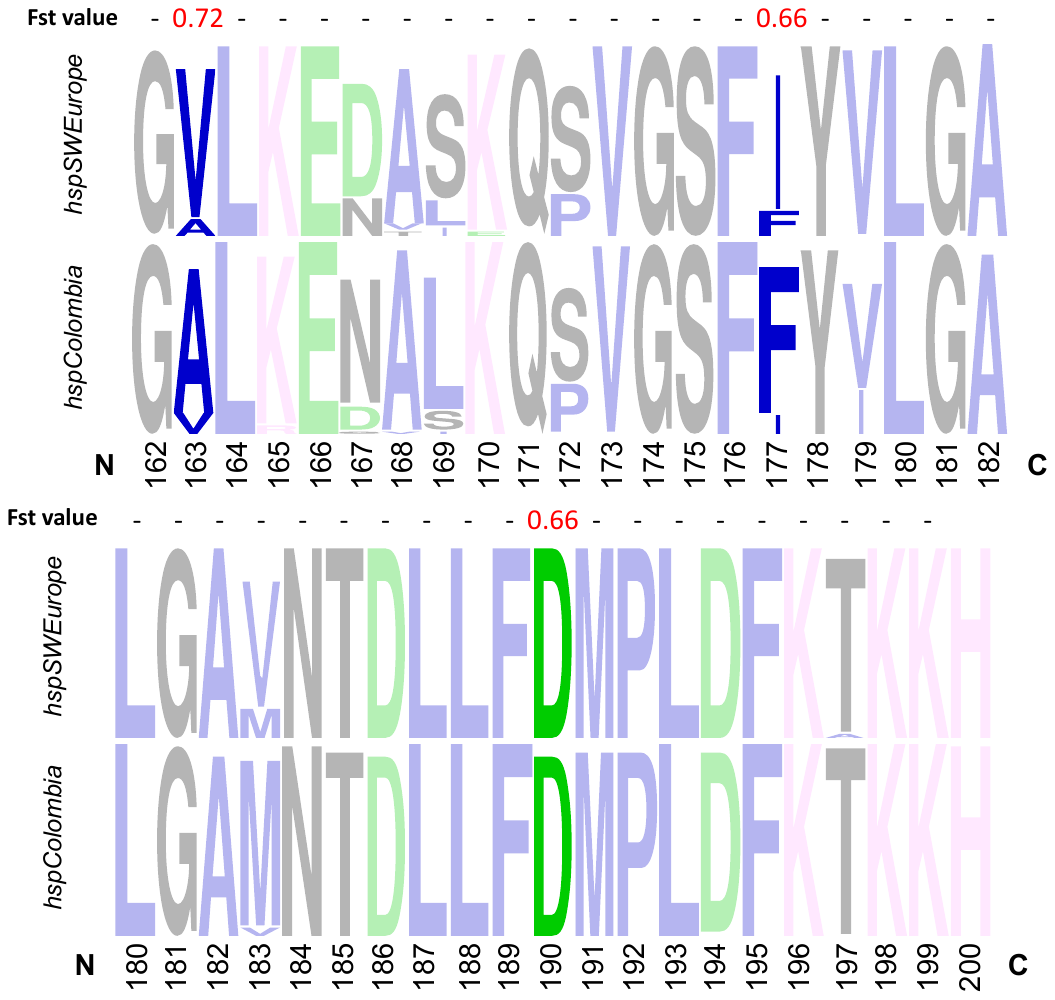

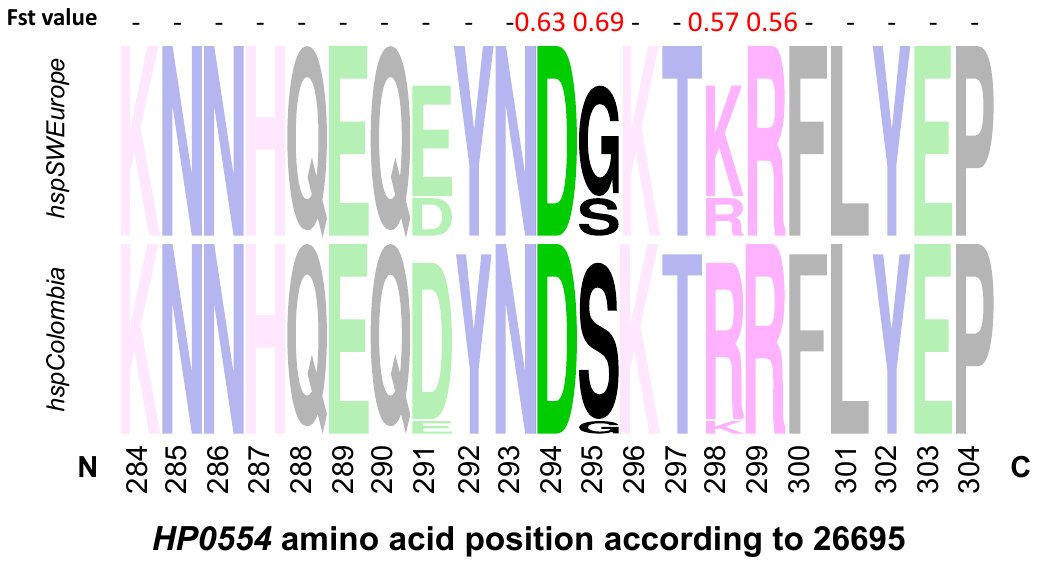

F) HopF (HP0252)

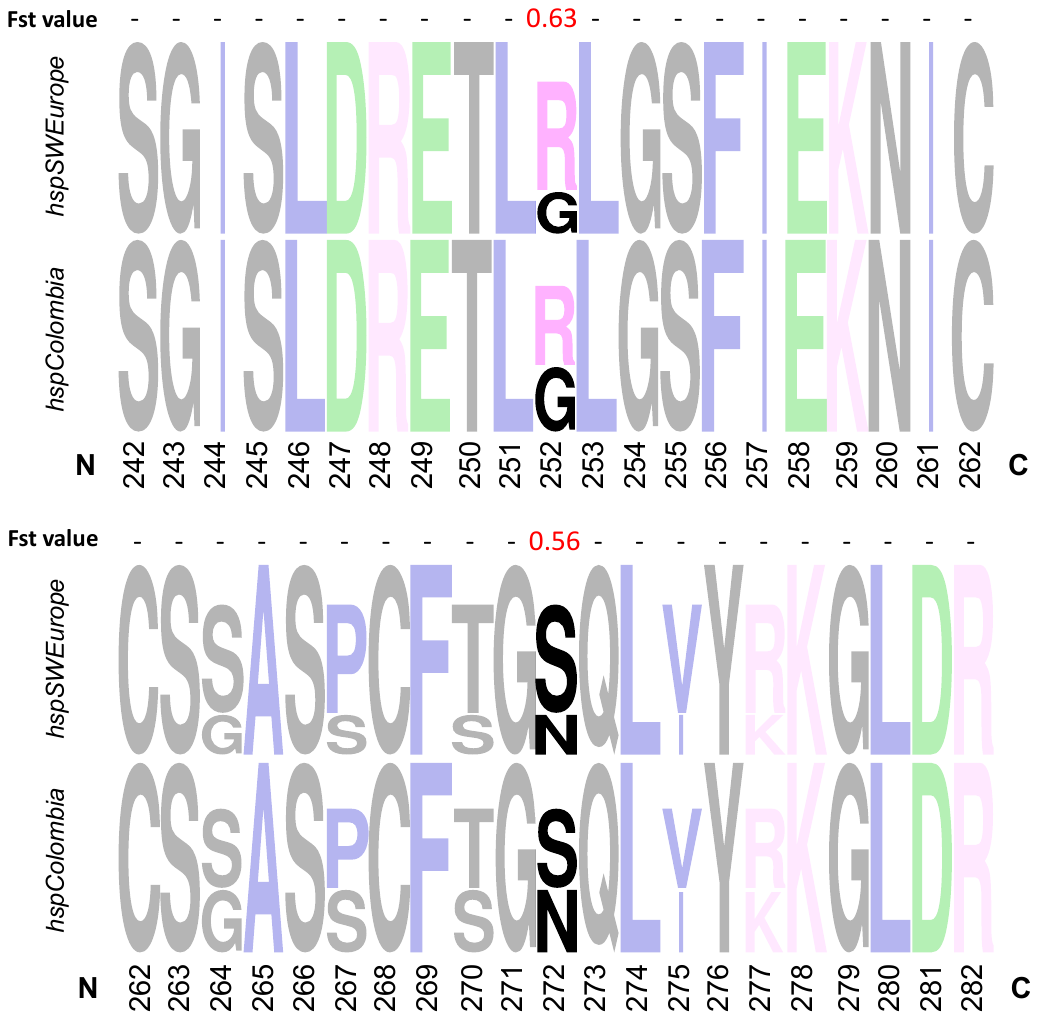

G) Thiol:disulfide interchange protein (HP0377)

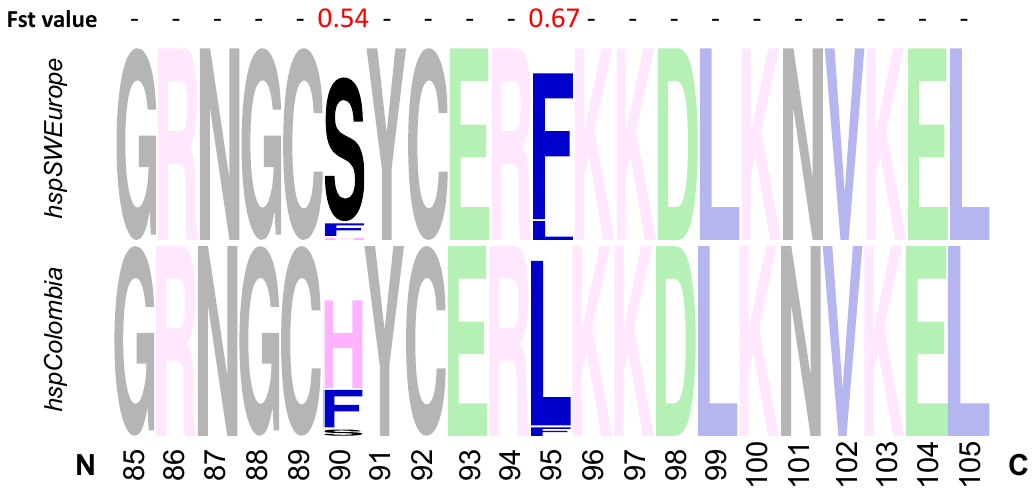

H) FrpB4 (HP1512)

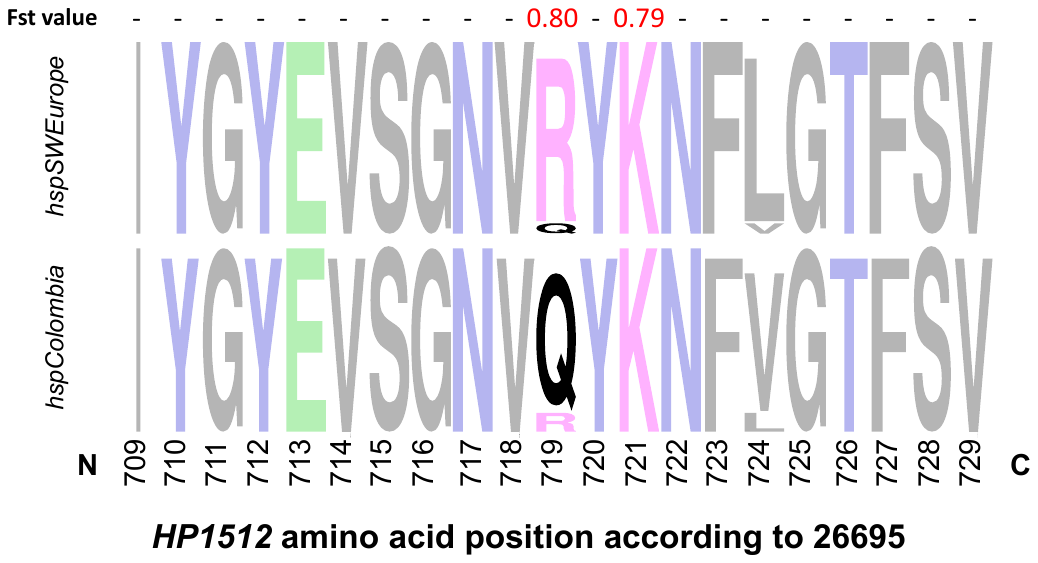

**Supplemental Figure 7. Fst values and WebLogo representations of non-synonymous changes in the 26 genes with a significative Fst value between *hspColombia* and *hspSWEurope* subpopulations.** The upper rows are Fst values calculated for isolates from *hspSWEurope* compared to isolates from the hspColombia subpopulations, and WebLogo representations show the region for which the Fst was calculate.
