## Supplemental tables for "Selective pressure on membrane proteins drives the evolution of *Helicobacter pylori* Colombian subpopulations"

### Table supplementary 1. Detailed information of isolates included in the study.

| **#** | **ID** | **Strain** | **Country** | **Region** | **Previous Population Assigned** | **Current Population Assigned** | **Genome Length** | **Number of Contigs** | **Average GC Content** | **Number Of Genes** |
| --- | --- | --- | --- | --- | --- | --- | --- | --- | --- | --- |
| 1 | AFR001 | K26A1 | Angola | Unknown | *hpAfrica2* | *hpAfrica2* | 1570310 | 1 | 38.7 | 1480 |
| 2 | AFR002 | EG_223 | Egypt | Unknown | *hspSWEurope* | *hspSWEuropeAustralia* | 1566053 | 27 | 38.4 | 1463 |
| 3 | AFR003 | GAM100Ai | Gambia | Unknown | *hspAfrica1WAfrica* | *hspAfrica1WAfrica* | 1633043 | 46 | 38.7 | 1529 |
| 4 | AFR004 | GAM101Biv | Gambia | Unknown | *hspAfrica1WAfrica* | *hspAfrica1WAfrica* | 1616343 | 53 | 38.5 | 1518 |
| 5 | AFR005 | GAM103Bi | Gambia | Unknown | *hspAfrica1WAfrica* | *hspAfrica1WAfrica* | 1624694 | 45 | 39.2 | 1508 |
| 6 | AFR006 | GAM105Ai | Gambia | Unknown | *hspAfrica1WAfrica* | *hspAfrica1WAfrica* | 1672031 | 47 | 39.1 | 1555 |
| 7 | AFR007 | GAM112Ai | Gambia | Unknown | *hspAfrica1WAfrica* | *hspAfrica1WAfrica* | 1626900 | 51 | 39.4 | 1522 |
| 8 | AFR008 | GAM114Ai | Gambia | Unknown | *hspAfrica1WAfrica* | *hspAfrica1WAfrica* | 1623315 | 31 | 40.1 | 1508 |
| 9 | AFR009 | GAM115Ai | Gambia | Unknown | *hspAfrica1WAfrica* | *hspAfrica1WAfrica* | 1685428 | 58 | 39.1 | 1574 |
| 10 | AFR010 | GAM117Ai | Gambia | Unknown | *hspAfrica1WAfrica* | *hspAfrica1WAfrica* | 1644727 | 60 | 38.7 | 1530 |
| 11 | AFR011 | GAM118Bi | Gambia | Unknown | *hspAfrica1WAfrica* | *hspAfrica1WAfrica* | 1684130 | 60 | 38.3 | 1558 |
| 12 | AFR012 | GAM119Bi | Gambia | Unknown | *hspAfrica1WAfrica* | *hspAfrica1WAfrica* | 1662596 | 47 | 39.0 | 1552 |
| 13 | AFR013 | GAM120Ai | Gambia | Unknown | *hspAfrica1WAfrica* | *hspAfrica1WAfrica* | 1688093 | 52 | 38.9 | 1564 |
| 14 | AFR014 | GAM121Aii | Gambia | Unknown | *hspAfrica1WAfrica* | *hspAfrica1WAfrica* | 1666274 | 51 | 38.7 | 1528 |
| 15 | AFR015 | GAM201Ai | Gambia | Unknown | *hspAfrica1WAfrica* | *hspAfrica1WAfrica* | 1615922 | 49 | 38.6 | 1490 |
| 16 | AFR016 | GAM210Bi | Gambia | Unknown | *hspAfrica1NAmerica* | *hspAfrica1NAmerica* | 1622001 | 51 | 38.8 | 1516 |
| 17 | AFR017 | GAM231Ai | Gambia | Unknown | *hspAfrica1WAfrica* | *hspAfrica1WAfrica* | 1625679 | 59 | 38.0 | 1508 |
| 18 | AFR018 | GAM239Bi | Gambia | Unknown | *hspAfrica1WAfrica* | *hspAfrica1WAfrica* | 1634668 | 70 | 38.0 | 1526 |
| 19 | AFR019 | GAM244Ai | Gambia | Unknown | *hspAfrica1WAfrica* | *hspAfrica1WAfrica* | 1598686 | 65 | 38.1 | 1507 |
| 20 | AFR020 | GAM245Ai | Gambia | Unknown | *hspAfrica1WAfrica* | *hspAfrica1WAfrica* | 1653916 | 50 | 39.3 | 1532 |
| 21 | AFR021 | GAM246Ai | Gambia | Unknown | *hspAfrica1WAfrica* | *hspAfrica1WAfrica* | 1669835 | 46 | 39.2 | 1568 |
| 22 | AFR022 | GAM249T | Gambia | Unknown | *hspAfrica1WAfrica* | *hspAfrica1WAfrica* | 1629080 | 40 | 39.2 | 1524 |
| 23 | AFR023 | GAM250T | Gambia | Unknown | *hspAfrica1NAmerica* | *hspAfrica1NAmerica* | 1584596 | 50 | 39.2 | 1473 |
| 24 | AFR024 | GAM254Ai | Gambia | Unknown | *hspAfrica1WAfrica* | *hspAfrica1WAfrica* | 1642173 | 48 | 38.3 | 1531 |
| 25 | AFR025 | GAM260ASi | Gambia | Unknown | *hspAfrica1WAfrica* | *hspAfrica1WAfrica* | 1634306 | 60 | 38.5 | 1521 |
| 26 | AFR026 | GAM260Bi | Gambia | Unknown | *hspAfrica1WAfrica* | *hspAfrica1WAfrica* | 1671995 | 55 | 38.9 | 1559 |
| 27 | AFR027 | GAM260BSi | Gambia | Unknown | *hspAfrica1WAfrica* | *hspAfrica1WAfrica* | 1579292 | 57 | 38.8 | 1487 |
| 28 | AFR028 | GAM263BFi | Gambia | Unknown | *hspAfrica1WAfrica* | *hspAfrica1WAfrica* | 1653267 | 49 | 39.0 | 1529 |
| 29 | AFR029 | GAM264Ai | Gambia | Unknown | *hspAfrica1WAfrica* | *hspAfrica1WAfrica* | 1608832 | 43 | 38.8 | 1504 |
| 30 | AFR030 | GAM265BSii | Gambia | Unknown | *hspAfrica1WAfrica* | *hspAfrica1WAfrica* | 1665501 | 56 | 38.7 | 1547 |
| 31 | AFR031 | GAM270ASi | Gambia | Unknown | *hspAfrica1WAfrica* | *hspAfrica1WAfrica* | 1652380 | 44 | 38.8 | 1539 |
| 32 | AFR032 | GAM71Ai | Gambia | Unknown | *hspAfrica1WAfrica* | *hspAfrica1WAfrica* | 1605866 | 71 | 37.8 | 1494 |
| 33 | AFR033 | GAM80Ai | Gambia | Unknown | *hspAfrica1WAfrica* | *hspAfrica1WAfrica* | 1642594 | 44 | 38.6 | 1512 |
| 34 | AFR034 | GAM83Bi | Gambia | Unknown | *hspAfrica1WAfrica* | *hspAfrica1WAfrica* | 1612179 | 50 | 38.6 | 1523 |
| 35 | AFR035 | GAM96Ai | Gambia | Unknown | *hspAfrica1WAfrica* | *hspAfrica1WAfrica* | 1662167 | 72 | 38.2 | 1543 |
| 36 | AFR036 | Gambia94_24 | Gambia | Unknown | *hspAfrica1WAfrica* | *hspAfrica1WAfrica* | 1712468 | 2 | 38.3 | 1588 |
| 37 | AFR037 | GAMchjs106B | Gambia | Unknown | *hspAfrica1WAfrica* | *hspAfrica1WAfrica* | 1582190 | 32 | 39.5 | 1462 |
| 38 | AFR038 | GAMchjs114i | Gambia | Unknown | *hspAfrica1WAfrica* | *hspAfrica1WAfrica* | 1617328 | 53 | 38.4 | 1512 |
| 39 | AFR039 | GAMchjs117Ai | Gambia | Unknown | *hspAfrica1WAfrica* | *hspAfrica1WAfrica* | 1617564 | 50 | 38.3 | 1491 |
| 40 | AFR040 | GAMchjs124i | Gambia | Unknown | *hspAfrica1WAfrica* | *hspAfrica1WAfrica* | 1617681 | 49 | 38.8 | 1503 |
| 41 | AFR041 | GAMchjs136i | Gambia | Unknown | *hspAfrica1WAfrica* | *hspAfrica1WAfrica* | 1660591 | 63 | 38.6 | 1549 |
| 42 | AFR042 | HP116Bi | Gambia | Unknown | *hspAfrica1WAfrica* | *hspAfrica1WAfrica* | 1652841 | 51 | 39.7 | 1533 |
| 43 | AFR043 | SA40A | SouthAfrica | Mpumalanga | *hpAfrica2* | *hpAfrica2* | 1645049 | 42 | 37.9 | 1535 |
| 44 | AFR044 | SA37A | SouthAfrica | Mpumalanga | *hspNEurope* | *hspSEurope* | 1599946 | 43 | 36.5 | 1496 |
| 45 | AFR045 | SA144A | SouthAfrica | Mpumalanga | *hpAfrica2* | *hpAfrica2* | 1629847 | 37 | 37.7 | 1612 |
| 46 | AFR047 | SA155C | SouthAfrica | Mpumalanga | *hpAfrica2* | *hpAfrica2* | 1663996 | 53 | 37.4 | 1563 |
| 47 | AFR048 | SA156C | SouthAfrica | Mpumalanga | *hspAfrica1SAfrica* | *hspAfrica1SAfrica* | 1698250 | 70 | 37.3 | 1584 |
| 48 | AFR050 | SA158C | SouthAfrica | Mpumalanga | *hspAfrica1SAfrica* | *hspAfrica1SAfrica* | 1613772 | 52 | 38.7 | 1492 |
| 49 | AFR051 | SA160C | SouthAfrica | Mpumalanga | *hpAfrica2* | *hpAfrica2* | 1657073 | 50 | 38.3 | 1569 |
| 50 | AFR052 | SA161C | SouthAfrica | Mpumalanga | *hspAfrica1NAmerica* | *hspAfrica1SAfrica* | 1688820 | 60 | 37.6 | 1582 |
| 51 | AFR053 | SA164C | SouthAfrica | Mpumalanga | *hspSEurope* | *hspSEurope* | 1612829 | 54 | 37.1 | 1506 |
| 52 | AFR054 | SA166A | SouthAfrica | Mpumalanga | *hpAfrica2* | *hpAfrica2* | 1598966 | 47 | 37.7 | 1549 |
| 53 | AFR055 | SA168C | SouthAfrica | Mpumalanga | *hspAfrica1SAfrica* | *hspAfrica1SAfrica* | 1646600 | 68 | 37.7 | 1558 |
| 54 | AFR056 | SA169C | SouthAfrica | Mpumalanga | *hpAfrica2* | *hpAfrica2* | 1661612 | 55 | 37.6 | 1575 |
| 55 | AFR057 | SA170C | SouthAfrica | Mpumalanga | *hspAfrica1SAfrica* | *hspAfrica1SAfrica* | 1611204 | 49 | 38.2 | 1508 |
| 56 | AFR058 | SA171C | SouthAfrica | Mpumalanga | *hspSEurope* | *hspSEurope* | 1611340 | 37 | 37.8 | 1502 |
| 57 | AFR059 | SA172C | SouthAfrica | Mpumalanga | *hpAfrica2* | *hpAfrica2* | 1657599 | 48 | 38.2 | 1528 |
| 58 | AFR060 | SA173C | SouthAfrica | Mpumalanga | *hspSEurope* | *hspSEurope* | 1656714 | 57 | 37.9 | 1543 |
| 59 | AFR061 | SA174A | SouthAfrica | Mpumalanga | *hpAfrica2* | *hpAfrica2* | 1651580 | 84 | 38.9 | 1544 |
| 60 | AFR062 | SA175A | SouthAfrica | Mpumalanga | *hpAfrica2* | *hpAfrica2* | 1611900 | 37 | 38.5 | 1489 |
| 61 | AFR063 | SA194C | SouthAfrica | Mpumalanga | *hpAfrica2* | *hpAfrica2* | 1566326 | 52 | 37.6 | 1470 |
| 62 | AFR064 | SA213C | SouthAfrica | Mpumalanga | *hspSEurope* | *hspSEurope* | 1618027 | 96 | 36.2 | 1498 |
| 63 | AFR065 | SA220C | SouthAfrica | Mpumalanga | *hspAfrica1SAfrica* | *hspAfrica1SAfrica* | 1665550 | 59 | 37.6 | 1541 |
| 64 | AFR066 | SA221C | SouthAfrica | Mpumalanga | *hspSEurope* | *hspSEurope* | 1625551 | 41 | 37.6 | 1515 |
| 65 | AFR067 | SA222C | SouthAfrica | Mpumalanga | *hpAsia2* | *hpAsia2* | 1630022 | 38 | 39.2 | 1553 |
| 66 | AFR068 | SA226A | SouthAfrica | Mpumalanga | *hspAfrica1SAfrica* | *hspAfrica1SAfrica* | 1704847 | 56 | 37.7 | 1603 |
| 67 | AFR069 | SA227C | SouthAfrica | Mpumalanga | *hspAfrica1SAfrica* | *hspAfrica1SAfrica* | 1664860 | 63 | 37.2 | 1552 |
| 68 | AFR070 | SA233C | SouthAfrica | Mpumalanga | *hpAfrica2* | *hpAfrica2* | 1581357 | 42 | 38.4 | 1470 |
| 69 | AFR071 | SA251C | SouthAfrica | Mpumalanga | *hpAfrica2* | *hpAfrica2* | 1631426 | 52 | 37.7 | 1516 |
| 70 | AFR073 | SA253C | SouthAfrica | Mpumalanga | *hpAfrica2* | *hpAfrica2* | 1645891 | 42 | 37.9 | 1530 |
| 71 | AFR074 | SA29A | SouthAfrica | Mpumalanga | *hpAfrica2* | *hpAfrica2* | 1620625 | 41 | 38.3 | 1693 |
| 72 | AFR075 | SA302C | SouthAfrica | Mpumalanga | *hspSEurope* | *hspSEurope* | 1571643 | 47 | 37.2 | 1473 |
| 73 | AFR076 | SA303C | SouthAfrica | Mpumalanga | *hpAfrica2* | *hpAfrica2* | 1638198 | 87 | 36.5 | 1517 |
| 74 | AFR077 | SA30C | SouthAfrica | Mpumalanga | *hspAfrica1SAfrica* | *hspAfrica1SAfrica* | 1663603 | 85 | 36.2 | 1568 |
| 75 | AFR078 | SA36C | SouthAfrica | Mpumalanga | *hpAfrica2* | *hpAfrica2* | 1639584 | 50 | 36.7 | 1533 |
| 76 | AFR079 | SA45C | SouthAfrica | Mpumalanga | *hspAfrica1SAfrica* | *hspAfrica1SAfrica* | 1640441 | 43 | 38.1 | 1523 |
| 77 | AFR080 | SA46C | SouthAfrica | Mpumalanga | *hspAfrica1SAfrica* | *hspAfrica1SAfrica* | 1607413 | 46 | 38.2 | 1636 |
| 78 | AFR082 | SA162A | SouthAfrica | Mpumalanga | *hspAfrica1SAfrica* | *hspAfrica1SAfrica* | 1666848 | 67 | 37.1 | 1557 |
| 79 | AFR083 | SA214A | SouthAfrica | Mpumalanga | *hspAfrica1SAfrica* | *hspAfrica1SAfrica* | 1630634 | 52 | 37.4 | 1522 |
| 80 | AFR084 | SA301A | SouthAfrica | Mpumalanga | *hspAfrica1SAfrica* | *hspAfrica1SAfrica* | 1657549 | 50 | 37.7 | 1570 |
| 81 | AFR085 | SA35C | SouthAfrica | Mpumalanga | *hspAfrica1SAfrica* | *hspAfrica1SAfrica* | 1658847 | 41 | 37.7 | 1559 |
| 82 | AFR086 | CC33C | SouthAfrica | CapeTown | *hspAfrica1SAfrica* | *hspAfrica1SAfrica* | 1659899 | 1 | 38.9 | 1569 |
| 83 | AFR087 | SouthAfrica20 | SouthAfrica | Johannesburgo | *hpAfrica2* | *hpAfrica2* | 1622903 | 1 | 38.6 | 1579 |
| 84 | AFR089 | SouthAfrica7 | SouthAfrica | Johannesburgo | *hpAfrica2* | *hpAfrica2* | 1679829 | 2 | 36.1 | 1570 |
| 85 | AFR090 | SA34A | SouthAfrica | Mpumalanga | *hpAfrica2* | *hpAfrica2* | 1644723 | 30 | 37.8 | 1534 |
| 86 | AFR091 | SA146A | SouthAfrica | Mpumalanga | *hspAfrica1SAfrica* | *hspAfrica1SAfrica* | 1673021 | 42 | 37.6 | 1567 |
| 87 | AFR092 | SA156A | SouthAfrica | Mpumalanga | *hspAfrica1SAfrica* | *hspAfrica1SAfrica* | 1699733 | 50 | 37.3 | 1612 |
| 88 | AFR093 | SA157A | SouthAfrica | Mpumalanga | *hspAfrica1SAfrica* | *hspAfrica1SAfrica* | 1670045 | 69 | 37.9 | 1559 |
| 89 | AFR094 | SA158A | SouthAfrica | Mpumalanga | *hspAfrica1SAfrica* | *hspAfrica1SAfrica* | 1614685 | 36 | 39.3 | 1570 |
| 90 | AFR095 | SA161A | SouthAfrica | Mpumalanga | *hspAfrica1SAfrica* | *hspAfrica1NAmerica* | 1653643 | 47 | 38.1 | 1599 |
| 91 | AFR096 | SA168A | SouthAfrica | Mpumalanga | *hspAfrica1SAfrica* | *hspAfrica1SAfrica* | 1641562 | 69 | 37.6 | 1535 |
| 92 | AFR097 | SA220A | SouthAfrica | Mpumalanga | *hspAfrica1SAfrica* | *hspAfrica1SAfrica* | 1663953 | 74 | 37.2 | 1541 |
| 93 | AFR098 | SA227A | SouthAfrica | Mpumalanga | *hspAfrica1SAfrica* | *hspAfrica1SAfrica* | 1661470 | 61 | 37.8 | 1674 |
| 94 | AFR099 | SA30A | SouthAfrica | Mpumalanga | *hspAfrica1SAfrica* | *hspAfrica1SAfrica* | 1670038 | 62 | 36.8 | 1582 |
| 95 | AFR100 | SA35A | SouthAfrica | Mpumalanga | *hspAfrica1SAfrica* | *hspAfrica1SAfrica* | 1658759 | 59 | 38.4 | 1559 |
| 96 | AFR101 | SA45A | SouthAfrica | Mpumalanga | *hspAfrica1SAfrica* | *hspAfrica1SAfrica* | 1638238 | 54 | 38.2 | 1524 |
| 97 | AS001 | Mandalay03 | Burma | Unknown | *hpAsia2* | *hpAsia2* | 1603532 | 29 | 39.5 | 1491 |
| 98 | AS002 | Mandalay13 | Burma | Unknown | *hspSEurope* | *hspSEurope* | 1630410 | 47 | 38.2 | 1521 |
| 99 | AS003 | Mandalay30 | Burma | Unknown | *hpAsia2* | *hpAsia2* | 1596925 | 45 | 38.3 | 1495 |
| 100 | AS004 | Mandalay38 | Burma | Unknown | *hspEAsia* | *hspEastAsia* | 1575000 | 44 | 37.5 | 1503 |
| 101 | AS005 | Mandalay46 | Burma | Unknown | *hspSEurope* | *hspSEurope* | 1635936 | 45 | 38.2 | 1533 |
| 102 | AS006 | Mandalay60 | Burma | Unknown | *hpAsia2* | *hpAsia2* | 1609429 | 44 | 38.5 | 1515 |
| 103 | AS007 | Myanmar51 | Burma | Unknown | *hpAsia2* | *hpAsia2* | 1633349 | 45 | 38.7 | 1544 |
| 104 | AS008 | Myanmar52 | Burma | Unknown | *hpAsia2* | *hpAsia2* | 1621631 | 31 | 38.8 | 1534 |
| 105 | AS009 | Myanmar66 | Burma | Unknown | *hpAsia2* | *hpAsia2* | 1554504 | 26 | 39.5 | 1454 |
| 106 | AS010 | Yangon132 | Burma | Unknown | *hspSAmerind* | *hspAmerind* | 1599407 | 47 | 38.0 | 1494 |
| 107 | AS011 | Yangon142 | Burma | Unknown | *hpAsia2* | *hpAsia2* | 1628522 | 45 | 38.4 | 1533 |
| 108 | AS012 | Yangon159 | Burma | Unknown | *hspSAmerind* | *hspEastAsia* | 1574936 | 36 | 37.7 | 1503 |
| 109 | AS013 | Yangon173 | Burma | Unknown | *hspEAsia* | *hspEastAsia* | 1560017 | 38 | 37.8 | 1484 |
| 110 | AS014 | Yangon179 | Burma | Unknown | *hpAsia2* | *hpAsia2* | 1609742 | 34 | 38.6 | 1520 |
| 111 | AS015 | Yangon188 | Burma | Unknown | *hpAsia2* | *hpAsia2* | 1617637 | 40 | 38.9 | 1503 |
| 112 | AS016 | Yangon190 | Burma | Unknown | *hpAsia2* | *hpAsia2* | 1636179 | 55 | 37.3 | 1528 |
| 113 | AS017 | Yangon202 | Burma | Unknown | *hpAsia2* | *hpAsia2* | 1608290 | 33 | 38.7 | 1495 |
| 114 | AS018 | Yangon222 | Burma | Unknown | *hspEAsia* | *hspEastAsia* | 1646000 | 50 | 37.5 | 1570 |
| 115 | AS019 | Yangon233 | Burma | Unknown | *hpAsia2* | *hpAsia2* | 1594143 | 40 | 38.8 | 1503 |
| 116 | AS020 | Yangon244 | Burma | Unknown | *hspEAsia* | *hspEastAsia* | 1574084 | 44 | 37.3 | 1512 |
| 117 | AS021 | D33 | China | Qingdao | *hspEAsia* | *hspEastAsia* | 1545148 | 64 | 38.3 | 1469 |
| 118 | AS022 | HLJ039 | China | Beijing | *hspEAsia* | *hspEastAsia* | 1609997 | 34 | 36.9 | 1549 |
| 119 | AS023 | HLJHP193 | China | Heilongjiang | *hspEAsia* | *hspEastAsia* | 1569327 | 10 | 36.3 | 1491 |
| 120 | AS024 | HLJHP253 | China | Heilongjiang | *hspEAsia* | *hspEastAsia* | 1589603 | 11 | 38.0 | 1518 |
| 121 | AS025 | HLJHP256 | China | Heilongjiang | *hspEAsia* | *hspEastAsia* | 1576405 | 14 | 38.0 | 1494 |
| 122 | AS026 | HLJHP271 | China | Heilongjiang | *hspEAsia* | *hspEastAsia* | 1588229 | 11 | 37.1 | 1518 |
| 123 | AS027 | NY97-102 | China | HongKong | *hspEAsia* | *hspEastAsia* | 1579459 | 51 | 44.8 | 1493 |
| 124 | AS028 | NY97-103 | China | HongKong | *hspEAsia* | *hspEastAsia* | 1603417 | 52 | 43.1 | 1524 |
| 125 | AS029 | NY97-18 | China | HongKong | *hspEAsia* | *hspEastAsia* | 1603538 | 54 | 40.5 | 1521 |
| 126 | AS030 | NY97-20 | China | HongKong | *hspEAsia* | *hspEastAsia* | 1657230 | 74 | 41.9 | 1538 |
| 127 | AS031 | NY97-29 | China | HongKong | *hspEAsia* | *hspEastAsia* | 1627971 | 72 | 40.8 | 1541 |
| 128 | AS032 | NY97-38 | China | HongKong | *hspEAsia* | *hspEastAsia* | 1578910 | 63 | 40.0 | 1504 |
| 129 | AS034 | YN1-100 | China | Yunnan | *hspEAsia* | *hspEastAsia* | 1579076 | 1 | 38.8 | 1502 |
| 130 | AS037 | YN1_91 | China | Yunnan | *hspEAsia* | *hspEastAsia* | 1609835 | 11 | 39.0 | 1532 |
| 131 | AS038 | YN1-92 | China | Yunnan | *hspEAsia* | *hspEastAsia* | 1618974 | 1 | 38.6 | 1566 |
| 132 | AS039 | YN1-99 | China | Yunnan | *hspEAsia* | *hspEastAsia* | 1606530 | 1 | 38.7 | 1533 |
| 133 | AS040 | YN3-21 | China | Yunnan | *hspEAsia* | *hspEastAsia* | 1572044 | 1 | 38.9 | 1580 |
| 134 | AS042 | YN3-77 | China | Yunnan | *hspEAsia* | *hspEastAsia* | 1577728 | 1 | 38.8 | 1506 |
| 135 | AS046 | YN4-134 | China | Yunnan | *hspEAsia* | *hspEastAsia* | 1624549 | 1 | 38.6 | 1584 |
| 136 | AS048 | YN4-83 | China | Yunnan | *hspEAsia* | *hspEastAsia* | 1588166 | 1 | 38.9 | 1560 |
| 137 | AS049 | YN4_84 | China | Yunnan | *hspEAsia* | *hspEastAsia* | 1633405 | 9 | 37.9 | 1559 |
| 138 | AS050 | FD506 | Malaysia | KualaLumpur | *hpEastAsia* | *hspEastAsia* | 1615241 | 105 | 38.9 | 1510 |
| 139 | AS051 | FD568 | Malaysia | KualaLumpur | *hpEastAsia* | *hspEastAsia* | 1610163 | 113 | 39.2 | 1499 |
| 140 | AS052 | FD577 | Malaysia | KualaLumpur | *hpEastAsia* | *hspEastAsia* | 1625905 | 74 | 38.4 | 1541 |
| 141 | AS053 | UM023 | Malaysia | KualaLumpur | *hpEastAsia* | *hspEastAsia* | 1623075 | 34 | 38.5 | 1530 |
| 142 | AS054 | UM038 | Malaysia | KualaLumpur | *hpEastAsia* | *hspEastAsia* | 1762049 | 44 | 38.5 | 1617 |
| 143 | AS055 | UM065 | Malaysia | KualaLumpur | *hpEastAsia* | *hspEastAsia* | 1586653 | 38 | 38.5 | 1466 |
| 144 | AS056 | UM066 | Malaysia | KualaLumpur | *hpEastAsia* | *hspEastAsia* | 1694163 | 34 | 38.4 | 1587 |
| 145 | AS057 | UM077 | Malaysia | KualaLumpur | *hpEastAsia* | *hspEastAsia* | 1619377 | 53 | 38.1 | 1518 |
| 146 | AS058 | UM085 | Malaysia | KualaLumpur | *hpEastAsia* | *hspEastAsia* | 1645062 | 50 | 38.2 | 1524 |
| 147 | AS059 | UM111 | Malaysia | KualaLumpur | *hpEastAsia* | *hspEastAsia* | 1663127 | 38 | 38.4 | 1552 |
| 148 | AS060 | 83 | Unknown | Unknown | *hpEastAsia* | *hspEastAsia* | 1617426 | 1 | 38.7 | 1558 |
| 149 | AS061 | Santal49 | India | WestBengal | *hpAsia2* | *hpAsia2* | 1610830 | 2 | 38.2 | 1510 |
| 150 | AS063 | India7 | India | Unknown | *hpAsia2* | *hpAsia2* | 1675918 | 1 | 38.9 | 1569 |
| 151 | AS064 | L7 | India | Ladakh | *hpAsia2* | *hspAmerind* | 1617826 | 1 | 38.8 | 1551 |
| 152 | AS065 | NAK7 | India | Hyderabad | *hpAsia2* | *hpAsia2* | 1586710 | 68 | 38.2 | 1482 |
| 153 | AS066 | Qsc35 | India | Unknown | *hpAsia2* | *hpAsia2* | 1654841 | 1 | 38.8 | 1628 |
| 154 | AS067 | FD430 | Malaysia | KualaLumpur | *hpAsia2* | *hpAsia2* | 1640355 | 129 | 39.4 | 1535 |
| 155 | AS068 | FD535 | India | KualaLumpur | *hpAsia2* | *hpAsia2* | 1671564 | 81 | 39.5 | 1561 |
| 156 | AS069 | NAB47 | India | BangaloreNorth | *hpAsia2* | *hpAsia2* | 1585921 | 107 | 38.9 | 1472 |
| 157 | AS070 | UM067 | India | KualaLumpur | *Unknown* | *hpAsia2* | 1680838 | 44 | 38.9 | 1568 |
| 158 | AS071 | Manado_1 | Indonesia | Sulawesi | *hspEAsia* | *hspEastAsia* | 1539815 | 38 | 38.0 | 1479 |
| 159 | AS072 | 35A | Japan | Unknown | *hspEAsia* | *hspEastAsia* | 1566655 | 1 | 38.9 | 1497 |
| 160 | AS073 | 98_10 | Japan | Unknown | *hspEAsia* | *hspEastAsia* | 1571772 | 51 | 38.3 | 1535 |
| 161 | AS074 | CPY1124 | Japan | Yamaguchi | *hspEAsia* | *hspEastAsia* | 1560199 | 12 | 39.0 | 1535 |
| 162 | AS075 | CPY1313 | Japan | Yamaguchi | *hspEAsia* | *hspEastAsia* | 1581437 | 5 | 38.8 | 1550 |
| 163 | AS076 | CPY1662 | Japan | Yamaguchi | *hspEAsia* | *hspEastAsia* | 1595824 | 9 | 38.2 | 1529 |
| 164 | AS077 | CPY1962 | Japan | Yamaguchi | *hspEAsia* | *hspEastAsia* | 1561561 | 8 | 38.0 | 1513 |
| 165 | AS078 | CPY3281 | Japan | Yamaguchi | *hspEAsia* | *hspEastAsia* | 1606528 | 8 | 38.8 | 1607 |
| 166 | AS079 | CPY6081 | Japan | Yamaguchi | *hspEAsia* | *hspEastAsia* | 1599100 | 9 | 38.8 | 1560 |
| 167 | AS080 | CPY6261 | Japan | Yamaguchi | *hspEAsia* | *hspEastAsia* | 1608996 | 5 | 38.1 | 1555 |
| 168 | AS081 | CPY6271 | Japan | Yamaguchi | *hspEAsia* | *hspEastAsia* | 1602120 | 8 | 37.8 | 1533 |
| 169 | AS082 | CPY6311 | Japan | Yamaguchi | *hspEAsia* | *hspEastAsia* | 1591163 | 8 | 38.2 | 1536 |
| 170 | AS083 | F16 | Japan | Unknown | *hspEAsia* | *hspEastAsia* | 1575399 | 1 | 38.9 | 1507 |
| 171 | AS084 | F30 | Japan | Unknown | *hspEAsia* | *hspEastAsia* | 1579693 | 2 | 36.5 | 1496 |
| 172 | AS085 | F32 | Japan | Unknown | *hspEAsia* | *hspEastAsia* | 1581461 | 2 | 37.8 | 1504 |
| 173 | AS086 | F57 | Japan | Unknown | *hspEAsia* | *hspEastAsia* | 1609006 | 1 | 38.7 | 1528 |
| 174 | AS087 | oki102 | Japan | Okinawa | *hspSAmerind* | *hspAmerind* | 1633212 | 1 | 38.8 | 1525 |
| 175 | AS088 | oki112 | Japan | Okinawa | *hspSAmerind* | *hspAmerind* | 1637925 | 1 | 38.8 | 1533 |
| 176 | AS089 | oki128 | Japan | Okinawa | *hspSAmerind* | *hspEastAsia* | 1553826 | 1 | 39.0 | 1492 |
| 177 | AS090 | oki154 | Japan | Okinawa | *hspSAmerind* | *hspEastAsia* | 1599700 | 1 | 38.8 | 1533 |
| 178 | AS091 | oki422 | Japan | Okinawa | *hspSAmerind* | *hspAmerind* | 1634852 | 1 | 38.8 | 1535 |
| 179 | AS092 | oki673 | Japan | Okinawa | *hspSAmerind* | *hspEastAsia* | 1595058 | 1 | 38.8 | 1530 |
| 180 | AS093 | oki828 | Japan | Okinawa | *hspSAmerind* | *hspEastAsia* | 1600345 | 1 | 38.8 | 1537 |
| 181 | AS094 | oki898 | Japan | Okinawa | *hspSAmerind* | *hspAmerind* | 1634875 | 1 | 38.8 | 1514 |
| 182 | AS095 | OK113 | Japan | Okinawa | *hpEastAsia* | *hspEastAsia* | 1616617 | 1 | 38.7 | 1537 |
| 183 | AS097 | OK144 | Japan | Okinawa | *Unknown* | *hspAmerind* | 1655983 | 1 | 38.9 | 1556 |
| 184 | AS101 | OK188 | Japan | Okinawa | *Unknown* | *hspEastAsia* | 1619751 | 1 | 38.6 | 1565 |
| 185 | AS102 | OK301 | Japan | Okinawa | *Unknown* | *hspEastAsia* | 1573576 | 2 | 35.7 | 1522 |
| 186 | AS103 | OK302 | Japan | Okinawa | *Unknown* | *hspEastAsia* | 1563266 | 1 | 38.9 | 1488 |
| 187 | AS104 | OK305 | Japan | Okinawa | *Unknown* | *hspEastAsia* | 1642131 | 1 | 38.6 | 1606 |
| 188 | AS105 | OK306 | Japan | Okinawa | *Unknown* | *hspEastAsia* | 1587201 | 1 | 38.9 | 1524 |
| 189 | AS106 | OK308 | Japan | Okinawa | *Unknown* | *hspEastAsia* | 1597265 | 1 | 38.8 | 1554 |
| 190 | AS107 | OK310 | Japan | Okinawa | *hpEastAsia* | *hspEastAsia* | 1595436 | 2 | 37.6 | 1511 |
| 191 | AS108 | OK311 | Japan | Okinawa | *Unknown* | *hspEastAsia* | 1583795 | 1 | 38.9 | 1513 |
| 192 | AS109 | OK312 | Japan | Okinawa | *Unknown* | *hspEastAsia* | 1567222 | 1 | 38.9 | 1649 |
| 193 | AS110 | OK313 | Japan | Okinawa | *Unknown* | *hspEastAsia* | 1608985 | 1 | 38.7 | 1535 |
| 194 | AS111 | OK314 | Japan | Okinawa | *Unknown* | *hspEastAsia* | 1652988 | 1 | 38.6 | 1572 |
| 195 | AS112 | OK316 | Japan | Okinawa | *Unknown* | *hspEastAsia* | 1768253 | 3 | 36.4 | 1638 |
| 196 | AS113 | OK317 | Japan | Okinawa | *Unknown* | *hspSWEuropeSouthAmerica* | 1649151 | 1 | 38.9 | 1572 |
| 197 | AS114 | 22 | Kuwait | KuwaitCity | *hspSEurope* | *hspSEurope* | 1650014 | 73 | 38.6 | 1535 |
| 198 | AS115 | 45 | Kuwait | KuwaitCity | *hspSEurope* | *hspSEurope* | 1642618 | 55 | 38.6 | 1514 |
| 199 | AS116 | 59 | Kuwait | KuwaitCity | *hspSEurope* | *hspSEurope* | 1667210 | 59 | 38.7 | 1539 |
| 200 | AS117 | FD662 | Malaysia | KualaLumpur | *hpAsia2* | *hpAsia2* | 1659453 | 66 | 39.2 | 1552 |
| 201 | AS118 | FD719 | Malaysia | KualaLumpur | *hpAsia2* | *hpAsia2* | 1642171 | 79 | 39.5 | 1554 |
| 202 | AS119 | FD423 | Malaysia | KualaLumpur | *hpAsia2* | *hpAsia2* | 1623128 | 115 | 39.6 | 1518 |
| 203 | AS120 | FD703 | Malaysia | KualaLumpur | *hpAsia2* | *hpAsia2* | 1676283 | 87 | 39.8 | 1551 |
| 204 | AS121 | UM084 | Malaysia | KualaLumpur | *hpAsia2* | *hpAsia2* | 1656826 | 34 | 39.1 | 1536 |
| 205 | AS122 | NP04 | Nepal | Unknown | *hpAsia2* | *hpAsia2* | 1608578 | 46 | 38.3 | 1507 |
| 206 | AS123 | NP05 | Nepal | Unknown | *hpAsia2* | *hpAsia2* | 1648417 | 49 | 38.2 | 1551 |
| 207 | AS124 | NP05-105 | Nepal | Unknown | *hpAsia2* | *hpAsia2* | 1590286 | 41 | 38.3 | 1499 |
| 208 | AS125 | NP05-107 | Nepal | Unknown | *hpAsia2* | *hpAsia2* | 1584839 | 42 | 38.5 | 1506 |
| 209 | AS126 | NP05-112 | Nepal | Unknown | *hpAsia2* | *hpAsia2* | 1571516 | 35 | 38.4 | 1480 |
| 210 | AS127 | NP05-121 | Nepal | Unknown | *hpAsia2* | *hpAsia2* | 1575953 | 32 | 39.3 | 1483 |
| 211 | AS128 | NP05-124 | Nepal | Unknown | *hpAsia2* | *hspSEurope* | 1619121 | 32 | 38.8 | 1523 |
| 212 | AS129 | NP05-227 | Nepal | Unknown | *hpAsia2* | *hspSEurope* | 1610308 | 44 | 38.9 | 1494 |
| 213 | AS130 | NP05-234 | Nepal | Unknown | *hpAsia2* | *hpAsia2* | 1608395 | 68 | 38.2 | 1511 |
| 214 | AS131 | NP05-250 | Nepal | Unknown | *hpAsia2* | *hpAsia2* | 1576527 | 35 | 38.6 | 1491 |
| 215 | AS132 | NP05-261 | Nepal | Unknown | *hspSEurope* | *hspSEurope* | 1623623 | 43 | 38.9 | 1525 |
| 216 | AS133 | NP05-266 | Nepal | Unknown | *hpAsia2* | *hpAsia2* | 1613063 | 47 | 38.5 | 1508 |
| 217 | AS134 | NP05-272 | Nepal | Unknown | *hpAsia2* | *hpAsia2* | 1632720 | 43 | 38.7 | 1545 |
| 218 | AS135 | NP05-278 | Nepal | Unknown | *hpAsia2* | *hpAsia2* | 1582831 | 43 | 38.4 | 1483 |
| 219 | AS136 | NP05-282 | Nepal | Unknown | *hspSEurope* | *hspSEurope* | 1573324 | 43 | 38.2 | 1467 |
| 220 | AS137 | 132 | Singapore | Unknown | *hpAsia2* | *hpAsia2* | 1631683 | 29 | 38.6 | 1541 |
| 221 | AS138 | 178 | Singapore | Unknown | *hspEAsia* | *hspEastAsia* | 1593673 | 26 | 38.2 | 1527 |
| 222 | AS139 | 241 | Singapore | Unknown | *hspEAsia* | *hspEastAsia* | 1544935 | 26 | 38.4 | 1462 |
| 223 | AS140 | 428 | Singapore | Unknown | *hspEAsia* | *hspEastAsia* | 1560870 | 29 | 38.4 | 1489 |
| 224 | AS141 | 1177 | Singapore | Unknown | *hspEAsia* | *hspEastAsia* | 1584044 | 27 | 38.7 | 1532 |
| 225 | AS142 | H30 | Singapore | Unknown | *hspEAsia* | *hspEastAsia* | 1604097 | 25 | 38.4 | 1528 |
| 226 | AS143 | H9 | Singapore | Unknown | *hspEAsia* | *hspEastAsia* | 1573907 | 29 | 38.3 | 1499 |
| 227 | AS144 | S380A | Singapore | Unknown | *hspEAsia* | *hspEastAsia* | 1586987 | 30 | 38.1 | 1505 |
| 228 | AS145 | S468A | Singapore | Unknown | *hspEAsia* | *hspEastAsia* | 1610344 | 27 | 38.6 | 1530 |
| 229 | AS146 | 51 | SouthKorea | Unknown | *hspEAsia* | *hspEastAsia* | 1589954 | 1 | 38.8 | 1513 |
| 230 | AS147 | 52 | SouthKorea | Unknown | *hspEAsia* | *hspEastAsia* | 1568826 | 1 | 38.9 | 1508 |
| 231 | AS148 | DU15 | SouthKorea | Seoul | *hspEAsia* | *hspEastAsia* | 1614411 | 1 | 38.7 | 1563 |
| 232 | AS149 | Hp238 | Taiwan | Unknown | *hspEAsia* | *hspEastAsia* | 1586473 | 1 | 38.7 | 1529 |
| 233 | AS150 | ML1 | Taiwan | Taipei | *hspEAsia* | *hspEastAsia* | 1629815 | 1 | 38.7 | 1610 |
| 234 | AS151 | ML2 | Taiwan | Taipei | *hspEAsia* | *hspEastAsia* | 1562125 | 1 | 38.9 | 1659 |
| 235 | AS152 | ML3 | Taiwan | Taipei | *hspEAsia* | *hspEastAsia* | 1635334 | 2 | 35.9 | 1637 |
| 236 | AS153 | Taiwan_47 | Taiwan | Unknown | *hspEAsia* | *hspEastAsia* | 1577473 | 58 | 38.2 | 1515 |
| 237 | AS154 | TW_235 | Taiwan | Unknown | *hspEAsia* | *hspEastAsia* | 1615482 | 19 | 38.7 | 1522 |
| 238 | AS155 | TW_265 | Taiwan | Unknown | *hspEAsia* | *hspEastAsia* | 1616398 | 25 | 38.9 | 1548 |
| 239 | AS156 | 8A3 | Unknown | Experimental | *hpEastAsia* | *hspEastAsia* | 1547179 | 44 | 39.0 | 1581 |
| 240 | AS157 | UM007 | Malaysia | Unknown | *hpEastAsia* | *hspEastAsia* | 1575854 | 72 | 38.1 | 1489 |
| 241 | AS158 | UM018 | Malaysia | Unknown | *hpAsia2* | *hpAsia2* | 1614546 | 72 | 38.8 | 1514 |
| 242 | AS159 | UM034 | Malaysia | Unknown | *hpEastAsia* | *hspEastAsia* | 1608430 | 65 | 38.1 | 1497 |
| 243 | AS160 | UM054 | Malaysia | Unknown | *hpAsia2* | *hpAsia2* | 1594474 | 81 | 38.7 | 1487 |
| 244 | AS161 | wls_5_14 | China | Zhejiang | *Unknown* | *hspEastAsia* | 1567825 | 67 | 38.1 | 1489 |
| 245 | AS162 | Ainu711 | Japan | Hokkaido | *hspAmerind* | *hspAmerind* | 1665512 | 50 | 38.0 | 1557 |
| 246 | AS163 | Ainu721 | Japan | Hokkaido | *hspAmerind* | *hspAmerind* | 1602314 | 47 | 38.5 | 1486 |
| 247 | AS164 | AinuN839 | Japan | Hokkaido | *hspAmerind* | *hspAmerind* | 1625420 | 44 | 39.4 | 1512 |
| 248 | AS165 | JPT1_552 | Japan | Okinawa | *hspMaori* | *hspEastAsia* | 1551269 | 67 | 39.1 | 1466 |
| 249 | AS166 | K15 | Japan | Hokkaido | *Unknown* | *hspEastAsia* | 1551453 | 26 | 38.7 | 1476 |
| 250 | AS167 | K16 | Japan | Hokkaido | *Unknown* | *hspEastAsia* | 1605921 | 42 | 37.6 | 1537 |
| 251 | AS168 | K17 | Japan | Hokkaido | *Unknown* | *hspEastAsia* | 1551153 | 32 | 38.6 | 1478 |
| 252 | AS169 | K21 | Japan | Hokkaido | *Unknown* | *hspEastAsia* | 1601277 | 34 | 38.2 | 1502 |
| 253 | AS170 | K22 | Japan | Hokkaido | *Unknown* | *hspEastAsia* | 1716397 | 31 | 38.7 | 1622 |
| 254 | AS171 | K23 | Japan | Hokkaido | *Unknown* | *hspEastAsia* | 1545937 | 32 | 37.4 | 1482 |
| 255 | AS172 | K24 | Japan | Hokkaido | *Unknown* | *hspEastAsia* | 1636901 | 40 | 38.4 | 1563 |
| 256 | AS173 | K25 | Japan | Hokkaido | *Unknown* | *hspEastAsia* | 1704131 | 49 | 38.2 | 1597 |
| 257 | AS174 | K26 | Japan | Hokkaido | *Unknown* | *hspEastAsia* | 1636410 | 42 | 38.5 | 1567 |
| 258 | AS175 | K27 | Japan | Hokkaido | *Unknown* | *hspEastAsia* | 1595263 | 30 | 38.3 | 1516 |
| 259 | AS176 | K28 | Japan | Hokkaido | *Unknown* | *hspEastAsia* | 1587752 | 36 | 38.0 | 1506 |
| 260 | AS177 | K29 | Japan | Hokkaido | *Unknown* | *hspEastAsia* | 1586927 | 31 | 38.0 | 1505 |
| 261 | AS178 | K30 | Japan | Hokkaido | *Unknown* | *hspEastAsia* | 1525716 | 22 | 39.0 | 1473 |
| 262 | AS179 | K32 | Japan | Hokkaido | *Unknown* | *hspEastAsia* | 1524698 | 25 | 38.6 | 1471 |
| 263 | AS180 | K33 | Japan | Hokkaido | *Unknown* | *hspEastAsia* | 1522513 | 31 | 38.9 | 1472 |
| 264 | AS181 | K34 | Japan | Hokkaido | *Unknown* | *hspEastAsia* | 1559976 | 22 | 38.9 | 1494 |
| 265 | AS182 | K35 | Japan | Hokkaido | *Unknown* | *hspEastAsia* | 1551184 | 37 | 38.3 | 1470 |
| 266 | AS183 | K36 | Japan | Hokkaido | *Unknown* | *hspEastAsia* | 1550771 | 33 | 38.6 | 1469 |
| 267 | AS184 | K37 | Japan | Hokkaido | *Unknown* | *hspEastAsia* | 1553375 | 39 | 38.5 | 1475 |
| 268 | AS185 | KR03B | Kyrgyzstan | Bishkek | *hspSiberia* | *hspAmerind* | 1637614 | 51 | 38.2 | 1516 |
| 269 | AS186 | KR16A | Kyrgyzstan | DzhalalAbad | *hspSiberia1* | *hspAmerind* | 1668885 | 45 | 37.3 | 1566 |
| 270 | AS187 | mong44 | Mongolia | Ulaangom | *hspSiberia2* | *hspAmerind* | 1610604 | 42 | 38.4 | 1505 |
| 271 | C001 | 1002 | Colombia | Cundinamarca | *Unknown* | *hspSWEuropeSouthAmerica* | 1609572 | 37 | 38.5 | 1503 |
| 272 | C002 | 1039 | Colombia | Cundinamarca | *Unknown* | *hspColombia* | 1665634 | 83 | 38.0 | 1548 |
| 273 | C003 | 1057 | Colombia | Cundinamarca | *Unknown* | *hspColombia* | 1681551 | 44 | 38.8 | 1573 |
| 274 | C004 | 1059 | Colombia | Cundinamarca | *Unknown* | *hspColombia* | 1628493 | 52 | 38.2 | 1517 |
| 275 | C005 | 1061 | Colombia | Cundinamarca | *Unknown* | *hspSWEuropeNorthAmerica* | 1644581 | 53 | 38.6 | 1536 |
| 276 | C006 | 1071 | Colombia | Cundinamarca | *Unknown* | *hspSWEuropeNorthAmerica* | 1658634 | 23 | 38.8 | 1555 |
| 277 | C007 | 1077 | Colombia | Cundinamarca | *Unknown* | *hspColombia* | 1655152 | 72 | 38.5 | 1544 |
| 278 | C008 | 1081 | Colombia | Cundinamarca | *Unknown* | *hspColombia* | 1648284 | 76 | 38.7 | 1521 |
| 279 | C009 | 1086 | Colombia | Cundinamarca | *Unknown* | *hspColombia* | 1606219 | 117 | 38.3 | 1465 |
| 280 | C010 | 1088 | Colombia | Cundinamarca | *Unknown* | *hspColombia* | 1648661 | 46 | 39.2 | 1527 |
| 281 | C011 | 1093 | Colombia | Cundinamarca | *Unknown* | *hspColombia* | 1643605 | 57 | 38.4 | 1532 |
| 282 | C012 | 1095 | Colombia | Cundinamarca | *Unknown* | *hspColombia* | 1669742 | 71 | 37.8 | 1566 |
| 283 | C013 | 1102 | Colombia | Cundinamarca | *Unknown* | *hspSWEuropeSouthAmerica* | 1611787 | 49 | 38.1 | 1494 |
| 284 | C014 | 2007 | Colombia | Cundinamarca | *Unknown* | *hspColombia* | 1678132 | 67 | 37.8 | 1560 |
| 285 | C015 | 2010 | Colombia | Cundinamarca | *Unknown* | *hspColombia* | 1669035 | 68 | 39.0 | 1532 |
| 286 | C016 | 2015 | Colombia | Cundinamarca | *Unknown* | *hspSWEuropeColombia* | 1606839 | 47 | 37.1 | 1497 |
| 287 | C017 | 2020 | Colombia | Boyaca | *hspSWEuropeColombia* | *hspSWEuropeColombia* | 1719718 | 149 | 39.3 | 1585 |
| 288 | C018 | 2021 | Colombia | Boyaca | *hspSWEuropeColombia* | *hspSWEuropeNorthAmerica* | 1678514 | 73 | 37.7 | 1571 |
| 289 | C019 | 2025 | Colombia | Boyaca | *hspSWEuropeColombia* | *hspColombia* | 1650159 | 61 | 37.7 | 1535 |
| 290 | C020 | 2027 | Colombia | Cundinamarca | *Unknown* | *hspColombia* | 1653213 | 81 | 38.2 | 1540 |
| 291 | C021 | 2029 | Colombia | Cundinamarca | *Unknown* | *hspColombia* | 1649365 | 49 | 38.7 | 1535 |
| 292 | C022 | 2036 | Colombia | Cundinamarca | *Unknown* | *hspColombia* | 1647948 | 60 | 38.5 | 1525 |
| 293 | C023 | 2040 | Colombia | Cundinamarca | *Unknown* | *hspColombia* | 1666338 | 91 | 38.0 | 1547 |
| 294 | C024 | 2047 | Colombia | Cundinamarca | *Unknown* | *hspColombia* | 1672229 | 52 | 37.8 | 1561 |
| 295 | C025 | 2061 | Colombia | Cundinamarca | *Unknown* | *hspSWEuropeSouthAmerica* | 1606714 | 58 | 38.4 | 1490 |
| 296 | C026 | 2065 | Colombia | Cundinamarca | *Unknown* | *hspColombia* | 1665508 | 61 | 38.1 | 1558 |
| 297 | C027 | 22003 | Colombia | Cundinamarca | *Unknown* | *hspColombia* | 1626407 | 35 | 39.0 | 1505 |
| 298 | C028 | 22013 | Colombia | Boyaca | *hspSWEuropeColombia* | *hspColombia* | 1660987 | 56 | 38.1 | 1548 |
| 299 | C029 | 22019 | Colombia | Santander | *hspAfrica1MiscAmerica* | *hspColombia* | 1633452 | 55 | 37.9 | 1525 |
| 300 | C030 | 22020 | Colombia | Boyaca | *hspSWEuropeColombia* | *hspSWEuropeColombia* | 1603665 | 58 | 38.4 | 1490 |
| 301 | C031 | 22021 | Colombia | Boyaca | *hspSWEuropeColombia* | *hspColombia* | 1621460 | 60 | 38.2 | 1512 |
| 302 | C032 | 22023 | Colombia | Boyaca | *hspSWEuropeColombia* | *hspSWEuropeColombia* | 1618831 | 37 | 38.1 | 1534 |
| 303 | C033 | 22025 | Colombia | Boyaca | *hspSWEuropeColombia* | *hspSWEuropeColombia* | 1636102 | 99 | 38.0 | 1510 |
| 304 | C034 | 22046 | Colombia | Boyaca | *hspSWEuropeColombia* | *hspColombia* | 1642160 | 51 | 38.1 | 1525 |
| 305 | C035 | 22087 | Colombia | Boyaca | *hspSWEuropeColombia* | *hspColombia* | 1658802 | 52 | 37.5 | 1539 |
| 306 | C036 | 22093 | Colombia | Boyaca | *hspSWEuropeColombia* | *hspColombia* | 1660420 | 92 | 38.4 | 1556 |
| 307 | C037 | 22095 | Colombia | Santander | *hspSWEuropeColombia* | *hspSWEuropeNorthAmerica* | 1670586 | 48 | 38.1 | 1551 |
| 308 | C038 | 22151 | Colombia | Cundinamarca | *Unknown* | *hspColombia* | 1674056 | 40 | 38.4 | 1543 |
| 309 | C039 | 22211 | Colombia | Cundinamarca | *Unknown* | *hspColombia* | 1679435 | 72 | 37.4 | 1549 |
| 310 | C040 | 22278 | Colombia | Cundinamarca | *Unknown* | *hspColombia* | 1615973 | 33 | 38.4 | 1512 |
| 311 | C041 | 22308 | Colombia | Cundinamarca | *Unknown* | *hspColombia* | 1695419 | 109 | 39.1 | 1555 |
| 312 | C042 | 22311 | Colombia | Cundinamarca | *hspSWEuropeColombia* | *hspColombia* | 1652825 | 58 | 37.5 | 1543 |
| 313 | C043 | 22312 | Colombia | Cundinamarca | *hspSWEuropeColombia* | *hspColombia* | 1634624 | 50 | 38.4 | 1525 |
| 314 | C044 | 22315 | Colombia | Boyaca | *hspSWEuropeColombia* | *hspColombia* | 1661814 | 61 | 37.7 | 1546 |
| 315 | C045 | 22317 | Colombia | Cundinamarca | *Unknown* | *hspColombia* | 1658120 | 64 | 37.0 | 1534 |
| 316 | C046 | 22322 | Colombia | Cundinamarca | *hspSWEuropeColombia* | *hspColombia* | 1656363 | 86 | 38.1 | 1539 |
| 317 | C047 | 22327 | Colombia | Cundinamarca | *hspSWEuropeColombia* | *hspSWEuropeColombia* | 1746488 | 245 | 39.0 | 1583 |
| 318 | C048 | 22331 | Colombia | Boyaca | *hspSWEuropeColombia* | *hspColombia* | 1673360 | 50 | 38.5 | 1552 |
| 319 | C049 | 22335 | Colombia | Boyaca | *hspSWEuropeColombia* | *hspColombia* | 1666250 | 88 | 38.9 | 1544 |
| 320 | C050 | 22336 | Colombia | Cundinamarca | *Unknown* | *hspColombia* | 1685604 | 114 | 38.7 | 1556 |
| 321 | C051 | 22337 | Colombia | Cundinamarca | *hspSWEuropeColombia* | *hspSWEuropeColombia* | 1685536 | 120 | 38.6 | 1574 |
| 322 | C052 | 22339 | Colombia | Cundinamarca | *hspSWEuropeColombia* | *hspSWEuropeColombia* | 1788959 | 265 | 39.3 | 1639 |
| 323 | C053 | 22341 | Colombia | Cundinamarca | *hspSWEuropeColombia* | *hspColombia* | 1683657 | 77 | 38.4 | 1544 |
| 324 | C054 | 22343 | Colombia | Cundinamarca | *Unknown* | *hspColombia* | 1760343 | 151 | 39.3 | 1640 |
| 325 | C055 | 22346 | Colombia | Cundinamarca | *hspSWEuropeColombia* | *hspSWEuropeSouthAmerica* | 1656254 | 93 | 39.1 | 1526 |
| 326 | C056 | 22347 | Colombia | Boyaca | *hspSWEuropeColombia* | *hspColombia* | 1652228 | 34 | 38.8 | 1531 |
| 327 | C057 | 22350 | Colombia | Cundinamarca | *Unknown* | *hspColombia* | 1757352 | 135 | 39.0 | 1642 |
| 328 | C058 | 22351 | Colombia | Cundinamarca | *hspSWEuropeColombia* | *hspColombia* | 1636502 | 75 | 39.2 | 1513 |
| 329 | C059 | 22352 | Colombia | Cundinamarca | *Unknown* | *hspColombia* | 1656538 | 46 | 38.8 | 1532 |
| 330 | C060 | 22360 | Colombia | Cundinamarca | *hspSWEuropeColombia* | *hspColombia* | 1688761 | 53 | 38.6 | 1574 |
| 331 | C061 | 22362 | Colombia | Tolima | *hspSWEuropeColombia* | *hspColombia* | 1664727 | 34 | 38.2 | 1575 |
| 332 | C062 | 22366 | Colombia | Cundinamarca | *hspAfrica1MiscAmerica* | *hspColombia* | 1641782 | 42 | 38.3 | 1528 |
| 333 | C063 | 22367 | Colombia | Boyaca | *hspSWEuropeColombia* | *hspColombia* | 1644660 | 45 | 39.3 | 1538 |
| 334 | C064 | 22368 | Colombia | Boyaca | *hspSWEuropeColombia* | *hspColombia* | 1652164 | 70 | 37.8 | 1532 |
| 335 | C065 | 22370 | Colombia | Boyaca | *hspSWEuropeColombia* | *hspColombia* | 1798914 | 254 | 40.0 | 1645 |
| 336 | C066 | 22371 | Colombia | Cundinamarca | *hspAfrica1MiscAmerica* | *hspSWEuropeNorthAmerica* | 1693405 | 152 | 38.2 | 1562 |
| 337 | C067 | 22377 | Colombia | Cundinamarca | *Unknown* | *hspSWEuropeNorthAmerica* | 1680132 | 82 | 38.0 | 1547 |
| 338 | C068 | 22378 | Colombia | Santander | *hspSWEuropeColombia* | *hspColombia* | 1655573 | 46 | 38.7 | 1547 |
| 339 | C069 | 22385 | Colombia | Santander | *hspSWEuropeColombia* | *hspSWEuropeSouthAmerica* | 1673830 | 54 | 38.3 | 1546 |
| 340 | C070 | 22386 | Colombia | Cundinamarca | *Unknown* | *hspSEurope* | 1642484 | 50 | 38.1 | 1570 |
| 341 | C071 | 22389 | Colombia | Cundinamarca | *hspSWEuropeColombia* | *hspSWEuropeColombia* | 1598649 | 42 | 37.3 | 1479 |
| 342 | C072 | 22390 | Colombia | Caldas | *hspSWEuropeMexico* | *hspColombia* | 1593210 | 56 | 39.1 | 1503 |
| 343 | C073 | 22393 | Colombia | Cundinamarca | *hspSWEuropeColombia* | *hspColombia* | 1755241 | 130 | 39.8 | 1620 |
| 344 | C074 | 22395 | Colombia | Cundinamarca | *Unknown* | *hspSWEuropeColombia* | 1765897 | 215 | 39.6 | 1622 |
| 345 | C075 | 22402 | Colombia | Cundinamarca | *hspEuropeColombia* | *hspColombia* | 1628208 | 48 | 38.2 | 1506 |
| 346 | C076 | 24004 | Colombia | Tolima | *hspAfrica1MiscAmerica* | *hspSWEuropeNorthAmerica* | 1664940 | 61 | 38.4 | 1547 |
| 347 | C077 | 24012 | Colombia | Cundinamarca | *Unknown* | *hspColombia* | 1598826 | 135 | 38.4 | 1496 |
| 348 | C078 | 24013 | Colombia | Cundinamarca | *Unknown* | *hspSWEurope* | 1646368 | 48 | 38.0 | 1542 |
| 349 | C079 | 26024 | Colombia | Meta | *hspSWEuropeColombia* | *hspColombia* | 1671004 | 43 | 38.6 | 1552 |
| 350 | C080 | 26083 | Colombia | Cundinamarca | *Unknown* | *hspColombia* | 1666814 | 47 | 38.6 | 1551 |
| 351 | C081 | 26084 | Colombia | Cundinamarca | *hspSWEuropeColombia* | *hspColombia* | 1651187 | 46 | 38.3 | 1527 |
| 352 | C082 | 26093 | Colombia | Cundinamarca | *hspSWEuropeColombia* | *hspColombia* | 1641735 | 53 | 38.5 | 1522 |
| 353 | C083 | 26100 | Colombia | Caqueta | *hspSWEuropeMexico* | *hspColombia* | 1654770 | 50 | 38.0 | 1535 |
| 354 | C084 | 3004 | Colombia | Cundinamarca | *Unknown* | *hspColombia* | 1687786 | 75 | 38.4 | 1566 |
| 355 | C085 | 3026 | Colombia | Cundinamarca | *Unknown* | *hspColombia* | 1678789 | 238 | 37.9 | 1538 |
| 356 | C086 | 3029 | Colombia | Cundinamarca | *Unknown* | *hspColombia* | 1645607 | 118 | 38.2 | 1520 |
| 357 | C087 | 3033 | Colombia | Cundinamarca | *Unknown* | *hspColombia* | 1626867 | 113 | 38.3 | 1511 |
| 358 | C088 | 3046 | Colombia | Cundinamarca | *Unknown* | *hspColombia* | 1677884 | 44 | 38.6 | 1558 |
| 359 | C089 | 3053 | Colombia | Cundinamarca | *Unknown* | *hspSWEuropeColombia* | 1610637 | 42 | 38.2 | 1519 |
| 360 | C090 | 3056 | Colombia | Cundinamarca | *Unknown* | *hspColombia* | 1630322 | 60 | 38.5 | 1520 |
| 361 | C091 | 3076 | Colombia | Cundinamarca | *Unknown* | *hspSWEuropeColombia* | 1648828 | 35 | 38.4 | 1550 |
| 362 | C092 | 3096 | Colombia | Cundinamarca | *Unknown* | *hspColombia* | 1665124 | 78 | 38.2 | 1554 |
| 363 | C093 | 3118 | Colombia | Cundinamarca | *Unknown* | *hspColombia* | 1638615 | 46 | 38.3 | 1527 |
| 364 | C094 | 3120 | Colombia | Cundinamarca | *Unknown* | *hspColombia* | 1638552 | 41 | 37.9 | 1512 |
| 365 | C095 | 3125 | Colombia | Cundinamarca | *Unknown* | *hspSWEuropeNorthAmerica* | 1643481 | 49 | 37.6 | 1538 |
| 366 | C096 | 3133 | Colombia | Cundinamarca | *Unknown* | *hspColombia* | 1684208 | 72 | 38.5 | 1588 |
| 367 | C097 | 3136 | Colombia | Cundinamarca | *Unknown* | *hspSWEuropeColombia* | 1605938 | 32 | 38.1 | 1505 |
| 368 | C098 | A033 | Colombia | Cundinamarca | *Unknown* | *hspAfrica1NAmerica* | 1672165 | 59 | 37.6 | 1537 |
| 369 | C099 | A037 | Colombia | Cundinamarca | *Unknown* | *hspColombia* | 1645991 | 50 | 38.8 | 1530 |
| 370 | C100 | A039 | Colombia | Cundinamarca | *Unknown* | *hspSWEuropeColombia* | 1597563 | 64 | 38.6 | 1474 |
| 371 | C101 | A077 | Colombia | Cundinamarca | *Unknown* | *hspColombia* | 1652075 | 55 | 38.6 | 1529 |
| 372 | C102 | CA22019 | Colombia | Santander | *hspAfrica1MiscAmerica* | *hspSWEuropeNorthAmerica* | 1663519 | 58 | 38.9 | 1521 |
| 373 | C103 | CA22095 | Colombia | Santander | *hspAfrica1MiscAmerica* | *hspSWEuropeColombia* | 1664226 | 197 | 39.8 | 1520 |
| 374 | C104 | CA22327 | Colombia | Santander | *hspSWEuropeColombia* | *hspColombia* | 1623028 | 61 | 37.9 | 1495 |
| 375 | C105 | CA22337 | Colombia | Cundinamarca | *hspSWEuropeColombia* | *hspSWEuropeColombia* | 1613682 | 35 | 38.7 | 1521 |
| 376 | C106 | CA22339 | Colombia | Cundinamarca | *hspSWEuropeColombia* | *hspSWEuropeSouthAmerica* | 1627265 | 56 | 38.9 | 1489 |
| 377 | C107 | CC22093 | Colombia | Cundinamarca | *hspSWEuropeColombia* | *hspSWEuropeColombia* | 1628294 | 85 | 39.7 | 1505 |
| 378 | C108 | CG22023 | Colombia | Boyaca | *hspSWEuropeColombia* | *hspColombia* | 1663913 | 44 | 38.7 | 1541 |
| 379 | C109 | CG22025 | Colombia | Boyaca | *hspSWEuropeColombia* | *hspColombia* | 1646335 | 42 | 38.3 | 1519 |
| 380 | C110 | CG22087 | Colombia | Cundinamarca | *hspSWEuropeColombia* | *hspColombia* | 1633543 | 29 | 38.6 | 1512 |
| 381 | C111 | CG22322 | Colombia | Cundinamarca | *hspSWEuropeColombia* | *hspSWEuropeColombia* | 1630058 | 63 | 38.8 | 1530 |
| 382 | C112 | CG22366 | Colombia | Cundinamarca | *hspAfrica1MiscAmerica* | *hspSWEuropeNorthAmerica* | 1662084 | 53 | 38.2 | 1542 |
| 383 | C113 | CG22385 | Colombia | Santander | *hspSWEuropeColombia* | *hspColombia* | 1667304 | 47 | 38.3 | 1540 |
| 384 | C114 | CM22013 | Colombia | Boyaca | *hspSWEuropeColombia* | *hspColombia* | 1617437 | 40 | 38.7 | 1509 |
| 385 | C115 | CM22046 | Colombia | Boyaca | *hspSWEuropeColombia* | *hspColombia* | 1659887 | 122 | 38.0 | 1533 |
| 386 | C116 | CM22315 | Colombia | Boyaca | *hspSWEuropeColombia* | *hspColombia* | 1645605 | 70 | 38.0 | 1534 |
| 387 | C117 | CM22331 | Colombia | Boyaca | *hspSWEuropeColombia* | *hspSWEuropeColombia* | 1596052 | 39 | 37.9 | 1484 |
| 388 | C118 | CM22341 | Colombia | Cundinamarca | *hspSWEuropeColombia* | *hspColombia* | 1667593 | 38 | 38.9 | 1557 |
| 389 | C119 | CM22346 | Colombia | Cundinamarca | *hspSWEuropeColombia* | *hspColombia* | 1624282 | 52 | 37.7 | 1518 |
| 390 | C120 | CM22347 | Colombia | Boyaca | *hspSWEuropeColombia* | *hspColombia* | 1690306 | 97 | 39.0 | 1549 |
| 391 | C121 | GCT27 | Colombia | Cauca'sValley | *Unknown* | *hspSWEuropeNorthAmerica* | 1643850 | 1 | 39.0 | 1539 |
| 392 | C122 | GCT43 | Colombia | Risaralda | *Unknown* | *hspSWEuropeNorthAmerica* | 1642466 | 1 | 39.0 | 1549 |
| 393 | C123 | GCT97 | Colombia | Tolima | *Unknown* | *hspSWEuropeNorthAmerica* | 1656646 | 1 | 39.0 | 1549 |
| 394 | C124 | NQ1671 | Colombia | Nariño | *hspAfrica1MiscAmerica* | *hspColombia* | 1626740 | 36 | 38.5 | 1515 |
| 395 | C126 | NQ1707 | Colombia | Nariño | *Unknown* | *hspColombia* | 1653263 | 80 | 38.5 | 1522 |
| 396 | C127 | NQ1712 | Colombia | Nariño | *Unknown* | *hspColombia* | 1572582 | 62 | 39.0 | 1493 |
| 397 | C128 | NQ315 | Colombia | Nariño | *hspEuropeColombia* | *hspColombia* | 1601938 | 57 | 38.6 | 1515 |
| 398 | C129 | NQ352 | Colombia | Nariño | *hspAfrica1MiscAmerica* | *hspColombia* | 1639151 | 61 | 38.9 | 1605 |
| 399 | C130 | NQ367 | Colombia | Nariño | *Unknown* | *hspColombia* | 1620442 | 76 | 38.5 | 1601 |
| 400 | C131 | NQ392 | Colombia | Nariño | *hspEuropeColombia* | *hspColombia* | 1653090 | 67 | 38.2 | 1546 |
| 401 | C132 | NQ4044 | Colombia | Nariño | *hspSWEuropeMexico* | *hspColombia* | 1728524 | 17 | 38.5 | 1648 |
| 402 | C133 | NQ4053 | Colombia | Nariño | *hspSWEuropeColombia* | *hspColombia* | 1652030 | 6 | 38.9 | 1559 |
| 403 | C134 | NQ4060 | Colombia | Nariño | *Unknown* | *hspColombia* | 1649200 | 55 | 38.8 | 1548 |
| 404 | C135 | NQ4076 | Colombia | Nariño | *hspSWEuropeColombia* | *hspColombia* | 1632709 | 4 | 39.2 | 1526 |
| 405 | C136 | NQ4099 | Colombia | Nariño | *hspSWEuropeColombia* | *hspColombia* | 1650644 | 6 | 39.0 | 1531 |
| 406 | C137 | NQ4110 | Colombia | Nariño | *hspSWEuropeMexico* | *hspSWEuropeColombia* | 1596647 | 2 | 39.2 | 1498 |
| 407 | C138 | NQ4161 | Colombia | Nariño | *hspAfrica1MiscAmerica* | *hspColombia* | 1639283 | 7 | 38.4 | 1560 |
| 408 | C139 | NQ4191 | Colombia | Nariño | *hspAfrica1MiscAmerica* | *hspColombia* | 1628609 | 40 | 39.0 | 1525 |
| 409 | C140 | NQ4200 | Colombia | Nariño | *hspSWEuropeColombia* | *hspColombia* | 1646737 | 14 | 37.0 | 1523 |
| 410 | C141 | NQ4216 | Colombia | Nariño | *hspSWEuropeColombia* | *hspColombia* | 1650078 | 12 | 38.3 | 1553 |
| 411 | C142 | NQ4228 | Colombia | Nariño | *hspSWEuropeColombia* | *hspColombia* | 1653281 | 6 | 38.9 | 1527 |
| 412 | C143 | PZ5004 | Colombia | Nariño | *hspAfrica1NAmerica* | *hspAfrica1NAmerica* | 1586499 | 259 | 39.3 | 1493 |
| 413 | C144 | PZ5026 | Colombia | Nariño | *hspEuropeColombia* | *hspColombia* | 1622716 | 224 | 39.0 | 1507 |
| 414 | C145 | PZ5056 | Colombia | Nariño | *hspEuropeColombia* | *hspColombia* | 1603341 | 298 | 39.3 | 1521 |
| 415 | C146 | PZ5080 | Colombia | Nariño | *hspEuropeColombia* | *hspColombia* | 1613577 | 243 | 39.1 | 1526 |
| 416 | C148 | PZ5005_3A3 | Colombia | Nariño | *hspAfrica1NAmerica* | *hspAfrica1NAmerica* | 1663098 | 51 | 38.7 | 1541 |
| 417 | C149 | PZ5006_3A3 | Colombia | Nariño | *hspAfrica1NAmerica* | *hspAfrica1NAmerica* | 1633699 | 53 | 38.3 | 1511 |
| 418 | C150 | PZ5009_3A2 | Colombia | Nariño | *hspSWEuropeColombia* | *hspColombia* | 1658224 | 53 | 37.7 | 1544 |
| 419 | C151 | PZ5016_3A3 | Colombia | Nariño | *hspSWEuropeColombia* | *hspAfrica1NAmerica* | 1618403 | 40 | 38.9 | 1506 |
| 420 | C152 | PZ5019_3A3 | Colombia | Nariño | *hspAfrica1MiscAmerica* | *hspAfrica1SAfrica* | 1673722 | 44 | 38.4 | 1550 |
| 421 | C153 | PZ5033_3A2 | Colombia | Nariño | *hspAfrica1MiscAmerica* | *hspColombia* | 1646537 | 60 | 38.1 | 1521 |
| 422 | C154 | PZ5086 | Colombia | Nariño | *hspEuropeColombia* | *hspColombia* | 1559914 | 244 | 38.9 | 1471 |
| 423 | C155 | SV328_2 | Colombia | Nariño | *hspSWEuropeColombia* | *hspColombia* | 1639757 | 56 | 38.3 | 1526 |
| 424 | C156 | SV340_2 | Colombia | Nariño | *hspAfrica1MiscAmerica* | *hspColombia* | 1626819 | 53 | 38.7 | 1512 |
| 425 | C157 | SV355_2 | Colombia | Nariño | *hspAfrica1MiscAmerica* | *hspColombia* | 1629307 | 39 | 39.5 | 1516 |
| 426 | C158 | SV380_1 | Colombia | Nariño | *hspAfrica1NAmerica* | *hspAfrica1NAmerica* | 1624379 | 40 | 38.5 | 1518 |
| 427 | C159 | SV397_2 | Colombia | Nariño | *hspAfrica1NAmerica* | *hspAfrica1NAmerica* | 1661366 | 47 | 39.1 | 1539 |
| 428 | C160 | SV449_1 | Colombia | Nariño | *hspSWEuropeColombia* | *hspColombia* | 1643791 | 41 | 39.1 | 1530 |
| 429 | C161 | 2006 | Colombia | Cundinamarca | *Unknown* | *hspColombia* | 1664276 | 319 | 38.2 | 1494 |
| 430 | C163 | 22345 | Colombia | Cundinamarca | *Unknown* | *hspColombia* | 1661685 | 43 | 38.0 | 1560 |
| 431 | C165 | SV376 | Colombia | Nariño | *Unknown* | *hspColombia* | 1678533 | 207 | 39.1 | 1560 |
| 432 | C166 | Hui1681 | Colombia | Amazon | *hspAmerind* | *hspAmerind* | 1504130 | 22 | 39.5 | 1438 |
| 433 | C167 | Hui1764 | Colombia | Amazon | *hspAmerind* | *hspAmerind* | 1604297 | 59 | 38.3 | 1542 |
| 434 | CAM001 | Nic01_C | Nicaragua | Managua | *hspSWEuropeHonduras* | *hspSWEuropeHonduras* | 1630930 | 41 | 38.2 | 1515 |
| 435 | CAM003 | Nic03_C | Nicaragua | Managua | *hspAfrica1Nicaragua* | *hspAfrica1Nicaragua* | 1611321 | 42 | 37.8 | 1492 |
| 436 | CAM004 | Nic04_C | Nicaragua | Managua | *hspSWEuropeHonduras* | *hspSWEuropeSouthAmerica* | 1685151 | 66 | 37.3 | 1559 |
| 437 | CAM005 | Nic05_C | Nicaragua | Managua | *hspSWEuropeHonduras* | *hspSWEuropeHonduras* | 1644906 | 32 | 37.7 | 1547 |
| 438 | CAM006 | Nic06_A | Nicaragua | Managua | *hspAfrica1Nicaragua* | *hspAfrica1Nicaragua* | 1670983 | 30 | 38.6 | 1554 |
| 439 | CAM007 | Nic07_C | Nicaragua | Managua | *hspAfrica1Nicaragua* | *hspAfrica1Nicaragua* | 1645731 | 26 | 39.2 | 1535 |
| 440 | CAM008 | Nic08_C | Nicaragua | Managua | *hspSEurope* | *hspSEurope* | 1574197 | 19 | 38.0 | 1495 |
| 441 | CAM009 | Nic09_C | Nicaragua | Managua | *hspSWEurope* | *hspSWEuropeSouthAmerica* | 1625652 | 33 | 38.4 | 1529 |
| 442 | CAM010 | Nic10_C | Nicaragua | Managua | *hspAfrica1Nicaragua* | *hspAfrica1Nicaragua* | 1655348 | 25 | 38.3 | 1550 |
| 443 | CAM011 | Nic11_C | Nicaragua | Managua | *hspAfrica1Nicaragua* | *hspAfrica1Nicaragua* | 1652673 | 57 | 38.4 | 1520 |
| 444 | CAM012 | Nic12_C | Nicaragua | Managua | *hspAfrica1Nicaragua* | *hspAfrica1Nicaragua* | 1677467 | 37 | 38.0 | 1561 |
| 445 | CAM013 | Nic13_C | Nicaragua | Managua | *hspAfrica1Nicaragua* | *hspAfrica1Nicaragua* | 1654238 | 36 | 37.6 | 1550 |
| 446 | CAM014 | Nic14_C | Nicaragua | Managua | *hspSWEurope* | *hspSWEuropeSouthAmerica* | 1677214 | 105 | 38.8 | 1580 |
| 447 | CAM015 | Nic15-C | Nicaragua | Managua | *hspSWEurope* | *hspSWEurope* | 1636264 | 32 | 38.2 | 1521 |
| 448 | CAM016 | Nic16_C | Nicaragua | Managua | *hspAfrica1Nicaragua* | *hspAfrica1Nicaragua* | 1649352 | 21 | 39.3 | 1535 |
| 449 | CAM017 | Nic17_C | Nicaragua | Managua | *hspSWEuropeMexico* | *hspSWEuropeSouthAmerica* | 1676425 | 47 | 37.9 | 1575 |
| 450 | CAM018 | Nic18_C | Nicaragua | Managua | *hspAfrica1Nicaragua* | *hspAfrica1Nicaragua* | 1645769 | 30 | 37.4 | 1516 |
| 451 | CAM019 | Nic19-C | Nicaragua | Managua | *hspAfrica1Nicaragua* | *hspAfrica1Nicaragua* | 1650301 | 21 | 38.4 | 1519 |
| 452 | CAM020 | Nic20-A | Nicaragua | Managua | *hspSWEurope* | *hspSWEuropeSouthAmerica* | 1601554 | 37 | 38.6 | 1495 |
| 453 | CAM021 | Nic20_C | Nicaragua | Managua | *hspAfrica1NAmerica* | *hspAfrica1NAmerica* | 1658954 | 42 | 38.2 | 1519 |
| 454 | CAM022 | Nic21_C | Nicaragua | Managua | *hspAfrica1Nicaragua* | *hspAfrica1Nicaragua* | 1654184 | 65 | 38.4 | 1541 |
| 455 | CAM023 | Nic22_A | Nicaragua | Managua | *hspSWEurope* | *hspSWEuropeSouthAmerica* | 1614655 | 75 | 39.0 | 1517 |
| 456 | CAM024 | Nic23_A | Nicaragua | Managua | *hspSWEuropeHonduras* | *hspAfrica1Nicaragua* | 1664714 | 52 | 38.2 | 1556 |
| 457 | CAM025 | Nic24_A | Nicaragua | Managua | *hspAfrica1Nicaragua* | *hspAfrica1Nicaragua* | 1644586 | 38 | 38.4 | 1514 |
| 458 | CAM026 | Nic25_A | Nicaragua | Managua | *hspAfrica1Nicaragua* | *hspAfrica1Nicaragua* | 1669942 | 50 | 38.6 | 1546 |
| 459 | CAM027 | Nic26_A | Nicaragua | Managua | *hspAfrica1Nicaragua* | *hspAfrica1Nicaragua* | 1649180 | 34 | 39.9 | 1529 |
| 460 | CAM028 | Nic27_A | Nicaragua | Managua | *hspAfrica1Nicaragua* | *hspAfrica1Nicaragua* | 1635965 | 37 | 38.5 | 1525 |
| 461 | CAM029 | Nic28_A | Nicaragua | Managua | *hspAfrica1Nicaragua* | *hspAfrica1Nicaragua* | 1644503 | 28 | 39.6 | 1519 |
| 462 | CAM030 | Nic29_A | Nicaragua | Managua | *hspAfrica1Nicaragua* | *hspAfrica1Nicaragua* | 1644168 | 39 | 39.8 | 1523 |
| 463 | CAM031 | Nic30_A | Nicaragua | Managua | *hspAfrica1Nicaragua* | *hspAfrica1Nicaragua* | 1669615 | 41 | 38.7 | 1543 |
| 464 | CAM032 | Nic31_A | Nicaragua | Managua | *hspSEurope* | *hspSWEuropeSouthAmerica* | 1595248 | 47 | 38.6 | 1498 |
| 465 | CAM033 | Nic32_A | Nicaragua | Managua | *hspAfrica1Nicaragua* | *hspAfrica1Nicaragua* | 1615475 | 51 | 38.0 | 1510 |
| 466 | CAM034 | ELS37 | ElSalvador | ruralregion | *hspSWEuropeHonduras* | *hspSWEuropeHonduras* | 1669876 | 2 | 37.4 | 1561 |
| 467 | CAM035 | Gt_04_059 | Guatemala | GuatemalaCity | *hspSWEuropeHonduras* | *hspSWEuropeHonduras* | 1671486 | 74 | 38.4 | 1556 |
| 468 | CAM036 | Gt_04_218 | Guatemala | Quetzaltenango | *hspSWEuropeMexico* | *hspSWEuropeSouthAmerica* | 1659996 | 21 | 39.1 | 1545 |
| 469 | CAM037 | HN_G240 | Honduras | Unknown | *hspAfrica1Nicaragua* | *hspAfrica1Nicaragua* | 1653842 | 57 | 37.5 | 1537 |
| 470 | CAM038 | HN_G242 | Honduras | Unknown | *hspSWEuropeHonduras* | *hspSWEuropeHonduras* | 1638023 | 46 | 38.5 | 1530 |
| 471 | CAM039 | HN_G246 | Honduras | Unknown | *hspAfrica1SAfrica* | *hspAfrica1SAfrica* | 1761945 | 238 | 39.4 | 1629 |
| 472 | CAM040 | HN_G248 | Honduras | Unknown | *hspSWEuropeHonduras* | *hspSWEuropeHonduras* | 1662158 | 37 | 37.7 | 1554 |
| 473 | CAM041 | HN_G249 | Honduras | Unknown | *hspSWEuropeHonduras* | *hspSWEuropeHonduras* | 1627027 | 35 | 40.1 | 1520 |
| 474 | CAM042 | HN_G250 | Honduras | Unknown | *hspSWEuropeHonduras* | *hspSWEuropeHonduras* | 1649537 | 36 | 38.1 | 1537 |
| 475 | CAM043 | HN_G251 | Honduras | Unknown | *hspSWEuropeHonduras* | *hspSWEuropeHonduras* | 1686922 | 69 | 37.7 | 1581 |
| 476 | CAM044 | HN_G255 | Honduras | Unknown | *hspSWEuropeHonduras* | *hspSWEuropeHonduras* | 1655180 | 47 | 37.7 | 1543 |
| 477 | CAM045 | HN_G257 | Honduras | Unknown | *hspSWEuropeHonduras* | *hspSWEuropeHonduras* | 1592316 | 24 | 38.7 | 1485 |
| 478 | CAM046 | HN_G258 | Honduras | Unknown | *hspAfrica1SAfrica* | *hspAfrica1SAfrica* | 1735043 | 220 | 38.7 | 1600 |
| 479 | CAM047 | HN_G260 | Honduras | Unknown | *hspSWEuropeHonduras* | *hspSWEuropeHonduras* | 1643151 | 269 | 39.0 | 1507 |
| 480 | CAM048 | HN_G262 | Honduras | Unknown | *hspSWEuropeHonduras* | *hspSWEuropeHonduras* | 1661382 | 46 | 37.4 | 1562 |
| 481 | CAM049 | HN_G265 | Honduras | Unknown | *hspAfrica1SAfrica* | *hspAfrica1SAfrica* | 1651709 | 44 | 39.5 | 1525 |
| 482 | CAM050 | HN_G266 | Honduras | Unknown | *hspSWEuropeHonduras* | *hspSWEuropeHonduras* | 1584704 | 38 | 37.9 | 1484 |
| 483 | CAM051 | HN_G267 | Honduras | Unknown | *hspAfrica1Nicaragua* | *hspAfrica1Nicaragua* | 1664986 | 72 | 37.3 | 1528 |
| 484 | CAM052 | HN_G274 | Honduras | Unknown | *hspSWEuropeHonduras* | *hspSWEuropeHonduras* | 1591116 | 28 | 37.9 | 1505 |
| 485 | CAM053 | HN_G277 | Honduras | Unknown | *hspSWEuropeHonduras* | *hspSWEuropeHonduras* | 1665595 | 40 | 38.1 | 1547 |
| 486 | CAM054 | HN_G278 | Honduras | Unknown | *hspSWEuropeHonduras* | *hspSWEuropeHonduras* | 1667060 | 48 | 37.6 | 1545 |
| 487 | CAM055 | HN_G280 | Honduras | Unknown | *hspAfrica1Nicaragua* | *hspAfrica1Nicaragua* | 1676356 | 54 | 37.1 | 1565 |
| 488 | CAM056 | HN_G285 | Honduras | Unknown | *hspAfrica1SAfrica* | *hspAfrica1SAfrica* | 1693697 | 35 | 37.9 | 1572 |
| 489 | CAM057 | HN_G288 | Honduras | Unknown | *hspSWEuropeHonduras* | *hspSWEuropeHonduras* | 1694674 | 229 | 39.2 | 1533 |
| 490 | CAM058 | HN_G300 | Honduras | Unknown | *hspSWEuropeHonduras* | *hspSWEuropeHonduras* | 1630928 | 42 | 38.0 | 1506 |
| 491 | CAM059 | HN_G307 | Honduras | Unknown | *hspSWEuropeHonduras* | *hspSWEuropeHonduras* | 1577133 | 24 | 38.8 | 1470 |
| 492 | CAM060 | HN_G310 | Honduras | Unknown | *hspSWEuropeHonduras* | *hspSWEuropeHonduras* | 1636129 | 62 | 38.6 | 1500 |
| 493 | CAM061 | HN_G315 | Honduras | Unknown | *hspSWEurope* | *hspSWEuropeSouthAmerica* | 1614237 | 31 | 38.5 | 1517 |
| 494 | CAM062 | HN_G328 | Honduras | Unknown | *hspSWEuropeHonduras* | *hspSWEuropeHonduras* | 1585876 | 33 | 38.4 | 1478 |
| 495 | CAM063 | HN_G331 | Honduras | Unknown | *hspSWEuropeHonduras* | *hspSWEuropeHonduras* | 1734997 | 150 | 38.6 | 1597 |
| 496 | CAM064 | HN_G339 | Honduras | Unknown | *hspSWEuropeHonduras* | *hspSWEuropeHonduras* | 1612625 | 70 | 37.6 | 1503 |
| 497 | CAM065 | HN_G341 | Honduras | Unknown | *hspSWEuropeHonduras* | *hspSWEuropeHonduras* | 1631808 | 44 | 38.1 | 1513 |
| 498 | CAM066 | HN_G346 | Honduras | Unknown | *hspAfrica1SAfrica* | *hspAfrica1SAfrica* | 1602868 | 38 | 37.7 | 1499 |
| 499 | EUR001 | 19027 | Belgium | Brussels | *hspSEurope* | *hspSEurope* | 1655142 | 60 | 38.5 | 1562 |
| 500 | EUR002 | 21580 | Belgium | Brussels | *hspSEurope* | *hspSEurope* | 1637810 | 41 | 39.1 | 1549 |
| 501 | EUR003 | 27935 | Belgium | Brussels | *hspSEurope* | *hspSEurope* | 1596878 | 42 | 38.9 | 1482 |
| 502 | EUR004 | 28861 | Belgium | Brussels | *hspSEurope* | *hspSEurope* | 1619541 | 47 | 39.3 | 1504 |
| 503 | EUR005 | 29009 | Belgium | Brussels | *hspSEurope* | *hspSEurope* | 1651445 | 46 | 38.9 | 1550 |
| 504 | EUR006 | 29373 | Belgium | Brussels | *hspNEurope* | *hspNEurope* | 1551759 | 31 | 39.6 | 1453 |
| 505 | EUR007 | 31181 | Belgium | Brussels | *hspSEurope* | *hspSEurope* | 1615744 | 43 | 39.0 | 1501 |
| 506 | EUR008 | 31235 | Belgium | Brussels | *hspSEurope* | *hspSEurope* | 1636948 | 60 | 38.1 | 1556 |
| 507 | EUR009 | 33375 | Belgium | Brussels | *hspSEurope* | *hspSEurope* | 1615951 | 51 | 38.9 | 1548 |
| 508 | EUR010 | 34320 | Belgium | Brussels | *hspSWEurope* | *hspSWEurope* | 1620829 | 44 | 39.0 | 1522 |
| 509 | EUR011 | 36166 | Belgium | Unknown | *hspSEurope* | *hspSEurope* | 1596627 | 44 | 39.1 | 1490 |
| 510 | EUR012 | 38185 | Belgium | Brussels | *hspSWEurope* | *hspSWEurope* | 1652406 | 71 | 38.4 | 1537 |
| 511 | EUR013 | 30950 | Belgium | Brussels | *hspSEurope* | *hspSEurope* | 1646302 | 36 | 38.6 | 1548 |
| 512 | EUR014 | 908 | France | Unknown | *hspAfrica1NAmerica* | *hspAfrica1NAmerica* | 1549666 | 1 | 39.3 | 1510 |
| 513 | EUR015 | 3697 | France | Bordeaux | *hspSWEurope* | *hspSWEurope* | 1616668 | 77 | 38.0 | 1518 |
| 514 | EUR016 | 3699 | France | Bordeaux | *hspSWEurope* | *hspSWEuropeNorthAmerica* | 1642798 | 48 | 38.6 | 1500 |
| 515 | EUR017 | 3738 | France | Bordeaux | *hspSWEurope* | *hspSWEurope* | 1595882 | 64 | 39.0 | 1482 |
| 516 | EUR018 | 3746 | France | Bordeaux | *hspSEurope* | *hspSEurope* | 1637877 | 63 | 38.2 | 1525 |
| 517 | EUR019 | 3754 | France | Bordeaux | *hpAsia2* | *hpAsia2* | 1605987 | 71 | 38.2 | 1494 |
| 518 | EUR020 | 3755 | France | Bordeaux | *hspAfrica1SAfrica* | *hspAfrica1SAfrica* | 1596484 | 54 | 39.1 | 1501 |
| 519 | EUR021 | 3800 | France | Bordeaux | *hspSWEurope* | *hspSWEuropeColombia* | 1601263 | 43 | 38.7 | 1500 |
| 520 | EUR022 | 3802 | France | Bordeaux | *hspAfrica1NAmerica* | *hspAfrica1WAfrica* | 1546782 | 64 | 38.7 | 1422 |
| 521 | EUR023 | 3824 | France | Bordeaux | *hspAfrica1MiscAmerica* | *hspSWEuropeNorthAmerica* | 1694860 | 90 | 38.9 | 1540 |
| 522 | EUR025 | 3843 | France | Bordeaux | *hspSEurope* | *hspSEurope* | 1490309 | 268 | 39.0 | 1366 |
| 523 | EUR026 | ANT170 | France | Bordeaux | *hspSWEurope* | *hspSWEurope* | 1634603 | 53 | 38.4 | 1524 |
| 524 | EUR027 | B25 | France | Bordeaux | *hspSWEurope* | *hspSWEurope* | 1602166 | 50 | 38.6 | 1480 |
| 525 | EUR028 | B29 | France | Bordeaux | *hspSEurope* | *hspSEurope* | 1589584 | 62 | 38.9 | 1490 |
| 526 | EUR029 | B30 | France | Bordeaux | *hspSEurope* | *hspSEurope* | 1597644 | 33 | 39.3 | 1490 |
| 527 | EUR030 | B31 | France | Bordeaux | *hspSWEurope* | *hspSWEuropeNorthAmerica* | 1678687 | 68 | 37.6 | 1550 |
| 528 | EUR031 | B35 | France | Bordeaux | *hspSEurope* | *hspSEurope* | 1633732 | 54 | 38.1 | 1555 |
| 529 | EUR032 | B38 | France | Unknown | *hspSEurope* | *hspSEurope* | 1576758 | 1 | 39.2 | 1501 |
| 530 | EUR033 | B40 | France | Bordeaux | *hspSEurope* | *hspSEurope* | 1600079 | 46 | 38.5 | 1493 |
| 531 | EUR034 | B41 | France | Bordeaux | *hspAfrica1NAmerica* | *hspAfrica1NAmerica* | 1684372 | 65 | 38.7 | 1564 |
| 532 | EUR035 | B43 | France | Bordeaux | *hspSWEurope* | *hspSWEurope* | 1640677 | 54 | 38.6 | 1513 |
| 533 | EUR036 | B44 | France | Bordeaux | *hspNEurope* | *hspNEurope* | 1619848 | 55 | 38.3 | 1538 |
| 534 | EUR037 | B45 | France | Unknown | *hspSWEurope* | *hspSWEurope* | 1602587 | 63 | 39.0 | 1566 |
| 535 | EUR038 | B47 | France | Bordeaux | *hspSEurope* | *hspSEurope* | 1595814 | 51 | 38.2 | 1486 |
| 536 | EUR039 | CHA185 | France | Bordeaux | *hspSEurope* | *hspSEurope* | 1635326 | 71 | 38.8 | 1518 |
| 537 | EUR040 | Fr-B58-M | France | Unknown | *hpEastAsia* | *hspEastAsia* | 1573221 | 46 | 38.2 | 1502 |
| 538 | EUR041 | Fr-G12-G | France | Unknown | *hspAfrica1MiscAmerica* | *hspSWEuropeNorthAmerica* | 1701531 | 55 | 38.1 | 1572 |
| 539 | EUR042 | GC23_HL | France | Bordeaux | *hspSEurope* | *hspSEurope* | 1629644 | 71 | 38.0 | 1547 |
| 540 | EUR043 | GC26_HL | France | Bordeaux | *hspSEurope* | *hspSEurope* | 1596021 | 51 | 38.9 | 1490 |
| 541 | EUR044 | GC30_HL | France | Bordeaux | *hspSEurope* | *hspSEurope* | 1609142 | 62 | 38.2 | 1522 |
| 542 | EUR045 | GC31_B | France | Bordeaux | *hspSEurope* | *hspSEurope* | 1656312 | 51 | 38.6 | 1546 |
| 543 | EUR046 | GC34-HL | France | Bordeaux | *hspSEurope* | *hspSEurope* | 1604342 | 53 | 39.0 | 1514 |
| 544 | EUR047 | GC52_HL | France | Bordeaux | *hspSWEuropeMexico* | *hspSWEurope* | 1641427 | 70 | 37.7 | 1540 |
| 545 | EUR048 | GC54_HL | France | Bordeaux | *hspSEurope* | *hspSEurope* | 1623691 | 50 | 38.6 | 1507 |
| 546 | EUR049 | GC62-HL-2 | France | Bordeaux | *hspSEurope* | *hspSEurope* | 1621557 | 75 | 39.4 | 1501 |
| 547 | EUR050 | GC65_HL | France | Bordeaux | *hspSEurope* | *hspSEurope* | 1668838 | 63 | 37.5 | 1551 |
| 548 | EUR051 | GC67_HL | France | Bordeaux | *hspSWEurope* | *hspSWEurope* | 1613756 | 56 | 38.4 | 1498 |
| 549 | EUR052 | GC69_HL | France | Bordeaux | *hspSEurope* | *hspSEurope* | 1601364 | 61 | 38.4 | 1494 |
| 550 | EUR053 | GRA247 | France | Bordeaux | *hspSWEurope* | *hspSWEurope* | 1687801 | 65 | 38.2 | 1571 |
| 551 | EUR054 | N6 | France | Unknown | *hspSEurope* | *hspSEurope* | 1657184 | 51 | 37.6 | 1639 |
| 552 | EUR055 | PHI092 | France | Bordeaux | *hspSEurope* | *hspSEurope* | 1662824 | 60 | 38.3 | 1549 |
| 553 | EUR056 | De_M53_M | Germany | Unknown | *hspSEurope* | *hspSEurope* | 1655282 | 88 | 38.4 | 1551 |
| 554 | EUR057 | P12 | Germany | Unknown | *hspSEurope* | *hspSEurope* | 1684038 | 2 | 36.9 | 1585 |
| 555 | EUR058 | SSR1 | Ireland | Dublin | *hspNEurope* | *hspNEurope* | 1607365 | 61 | 38.1 | 1508 |
| 556 | EUR059 | SSR12 | Ireland | Dublin | *hspSEurope* | *hspSEurope* | 1640508 | 84 | 37.6 | 1509 |
| 557 | EUR060 | SSR13 | Ireland | Dublin | *hspNEurope* | *hspNEurope* | 1617847 | 73 | 38.3 | 1513 |
| 558 | EUR061 | SSR14 | Ireland | Dublin | *hspNEurope* | *hspNEurope* | 1662723 | 70 | 38.6 | 1551 |
| 559 | EUR062 | SSR17 | Ireland | Dublin | *hspNEurope* | *hspNEurope* | 1580609 | 61 | 39.1 | 1485 |
| 560 | EUR063 | SSR2 | Ireland | Dublin | *hspNEurope* | *hspNEurope* | 1582107 | 68 | 38.7 | 1489 |
| 561 | EUR064 | SSR20 | Ireland | Dublin | *hspNEurope* | *hspNEurope* | 1610542 | 57 | 38.4 | 1498 |
| 562 | EUR065 | SSR22 | Ireland | Dublin | *hspNEurope* | *hspNEurope* | 1640253 | 66 | 38.7 | 1552 |
| 563 | EUR066 | SSR23 | Ireland | Dublin | *hspNEurope* | *hspNEurope* | 1644971 | 87 | 38.2 | 1530 |
| 564 | EUR067 | SSR3 | Ireland | Dublin | *hspNEurope* | *hspNEurope* | 1670700 | 79 | 38.2 | 1567 |
| 565 | EUR068 | SSR33 | Ireland | Dublin | *hspNEurope* | *hspNEurope* | 1681995 | 90 | 38.8 | 1554 |
| 566 | EUR069 | SSR40 | Ireland | Dublin | *hspNEurope* | *hspNEurope* | 1666663 | 86 | 38.1 | 1569 |
| 567 | EUR070 | SSR43 | Ireland | Dublin | *hspNEurope* | *hspSEurope* | 1538019 | 40 | 39.3 | 1441 |
| 568 | EUR071 | SSR5 | Ireland | Dublin | *hspNEurope* | *hspNEurope* | 1615339 | 60 | 39.0 | 1505 |
| 569 | EUR072 | SSR7 | Ireland | Dublin | *hspNEurope* | *hspNEurope* | 1654739 | 68 | 38.4 | 1543 |
| 570 | EUR073 | SSR8 | Ireland | Dublin | *hspNEurope* | *hspNEurope* | 1611252 | 84 | 38.7 | 1501 |
| 571 | EUR074 | SSR9 | Ireland | Dublin | *hspNEurope* | *hspNEurope* | 1703465 | 99 | 37.5 | 1609 |
| 572 | EUR075 | G27 | Italy | Unknown | *hspSEurope* | *hspSEurope* | 1663013 | 2 | 36.9 | 1575 |
| 573 | EUR076 | Lithuania75 | Lithuania | Unknown | *hspSEurope* | *hspSEurope* | 1640673 | 2 | 36.3 | 1553 |
| 574 | EUR077 | 1152_04 | Portugal | Unknown | *hspSWEurope* | *hspSWEurope* | 1588646 | 59 | 39.0 | 1501 |
| 575 | EUR078 | 1198_04 | Portugal | Unknown | *hspSWEurope* | *hspSWEurope* | 1618345 | 57 | 38.8 | 1491 |
| 576 | EUR079 | 173_00 | Portugal | Lisbon | *hspSWEurope* | *hspSWEurope* | 1573297 | 79 | 38.7 | 1462 |
| 577 | EUR080 | 1786_05 | Portugal | Unknown | *hspSWEurope* | *hspSWEurope* | 1619972 | 41 | 39.3 | 1528 |
| 578 | EUR081 | 207_99 | Portugal | Unknown | *hspSWEurope* | *hspSWEurope* | 1544449 | 47 | 38.5 | 1430 |
| 579 | EUR082 | 228_99 | Portugal | Unknown | *hspSWEurope* | *hspSWEurope* | 1619666 | 54 | 38.4 | 1512 |
| 580 | EUR083 | 499_02 | Portugal | Unknown | *hspSWEuropeMexico* | *hspSWEuropeSouthAmerica* | 1666743 | 73 | 38.3 | 1555 |
| 581 | EUR084 | 655_99 | Portugal | Unknown | *hspSWEurope* | *hspSWEurope* | 1615666 | 45 | 39.2 | 1492 |
| 582 | EUR085 | Pt1293U | Portugal | Unknown | *hspSWEurope* | *hspSWEurope* | 1663004 | 35 | 38.3 | 1562 |
| 583 | EUR086 | Pt-1846-U | Portugal | Unknown | *hspSWEurope* | *hspSWEurope* | 1642077 | 58 | 38.8 | 1534 |
| 584 | EUR087 | Pt1918U | Portugal | Unknown | *hspAfrica1SAfrica* | *hspAfrica1SAfrica* | 1635641 | 42 | 39.0 | 1523 |
| 585 | EUR088 | Pt21299RU | Portugal | Unknown | *hspSWEurope* | *hspSWEurope* | 1637161 | 42 | 38.9 | 1529 |
| 586 | EUR089 | Pt4472G | Portugal | Unknown | *hspSWEurope* | *hspSWEurope* | 1668431 | 50 | 38.0 | 1568 |
| 587 | EUR090 | Pt4481G | Portugal | Unknown | *hspSWEurope* | *hspSWEurope* | 1614566 | 60 | 38.7 | 1500 |
| 588 | EUR091 | Pt4497U | Portugal | Unknown | *hspAfrica1MiscAmerica* | *hspAfrica1SAfrica* | 1651496 | 84 | 38.2 | 1527 |
| 589 | EUR092 | Pt5322G | Portugal | Unknown | *hspSWEurope* | *hspSWEurope* | 1591902 | 38 | 37.8 | 1492 |
| 590 | EUR093 | Pt5771G | Portugal | Unknown | *hspAfrica1MiscAmerica* | *hspSWEuropeNorthAmerica* | 1716534 | 73 | 38.6 | 1590 |
| 591 | EUR094 | Pt_7739 | Portugal | SanSebastiao | *hspSWEurope* | *hspSWEurope* | 1665858 | 110 | 40.0 | 1529 |
| 592 | EUR095 | Pt_7757 | Portugal | SantaMariaFeira | *hspSWEurope* | *hspSWEurope* | 1618794 | 29 | 38.1 | 1515 |
| 593 | EUR096 | Pt_7854 | Portugal | AngraHeroismo | *hspSWEurope* | *hspSWEurope* | 1639643 | 45 | 37.2 | 1553 |
| 594 | EUR097 | Pt_7901 | Portugal | PontaDelgada | *hspSWEurope* | *hspSWEurope* | 1595218 | 47 | 39.4 | 1492 |
| 595 | EUR098 | Pt_8186 | Portugal | SanSebastiao | *hspSWEurope* | *hspSWEurope* | 1642281 | 42 | 38.3 | 1531 |
| 596 | EUR099 | Pt_8220 | Portugal | Setubal | *hspSWEurope* | *hspSWEurope* | 1576706 | 38 | 37.5 | 1478 |
| 597 | EUR100 | Pt_8259 | Portugal | Setubal | *hspSWEurope* | *hspSWEurope* | 1609636 | 22 | 39.0 | 1508 |
| 598 | EUR101 | Pt_8284 | Portugal | Setubal | *hspSWEurope* | *hspSWEurope* | 1631908 | 35 | 37.8 | 1539 |
| 599 | EUR102 | Pt_8311 | Portugal | Porto | *hspSWEurope* | *hspSWEuropeColombia* | 1592388 | 23 | 38.4 | 1496 |
| 600 | EUR103 | Pt_8323 | Portugal | VianaCastelo | *hspAfrica1SAfrica* | *hspAfrica1SAfrica* | 1640025 | 30 | 38.8 | 1546 |
| 601 | EUR104 | Pt_8360 | Portugal | Braga | *hspSWEurope* | *hspSWEurope* | 1636793 | 42 | 39.2 | 1532 |
| 602 | EUR105 | Pt_8376 | Portugal | Porto | *hspSWEurope* | *hspSWEurope* | 1578038 | 41 | 37.5 | 1473 |
| 603 | EUR106 | Pt_8427 | Portugal | Viseu | *hspSWEurope* | *hspSWEurope* | 1597251 | 26 | 38.4 | 1486 |
| 604 | EUR107 | Pt_8434 | Portugal | Viseu | *hspSWEurope* | *hspSWEurope* | 1645513 | 128 | 39.0 | 1523 |
| 605 | EUR108 | PtB89G | Portugal | Lisbon | *Unknown* | *hspSWEurope* | 1646346 | 44 | 38.4 | 1549 |
| 606 | EUR109 | PtB92G | Portugal | Lisbon | *Unknown* | *hspSWEurope* | 1648404 | 55 | 38.0 | 1552 |
| 607 | EUR110 | ES_10 | Spain | Madrid | *hspSWEurope* | *hspSWEurope* | 1636461 | 59 | 38.4 | 1528 |
| 608 | EUR111 | ES_11 | Spain | Madrid | *hspSWEurope* | *hspSWEurope* | 1628668 | 49 | 36.9 | 1511 |
| 609 | EUR112 | ES_12 | Spain | Madrid | *hspSEurope* | *hspSEurope* | 1674068 | 46 | 36.6 | 1569 |
| 610 | EUR113 | ES_13 | Spain | Madrid | *hspSWEurope* | *hspSWEurope* | 1545459 | 21 | 39.4 | 1462 |
| 611 | EUR114 | ES_15 | Spain | Madrid | *hspSWEurope* | *hspSWEurope* | 1586585 | 34 | 37.7 | 1496 |
| 612 | EUR115 | ES_25 | Spain | Madrid | *hspSWEurope* | *hspSWEurope* | 1617338 | 25 | 38.3 | 1506 |
| 613 | EUR116 | ES_26 | Spain | Madrid | *hspSWEurope* | *hspSWEurope* | 1612648 | 24 | 38.3 | 1515 |
| 614 | EUR117 | ES_31 | Spain | Madrid | *hspSWEurope* | *hspSWEurope* | 1600892 | 32 | 39.2 | 1501 |
| 615 | EUR118 | ES_46 | Spain | Madrid | *hspSWEurope* | *hspSWEurope* | 1645252 | 44 | 37.8 | 1557 |
| 616 | EUR119 | ES_54 | Spain | Madrid | *hspAfrica1MiscAmerica* | *hspSWEuropeNorthAmerica* | 1661373 | 52 | 37.6 | 1547 |
| 617 | EUR120 | ES_58 | Spain | Madrid | *hspSWEurope* | *hspSWEurope* | 1623725 | 26 | 38.8 | 1521 |
| 618 | EUR121 | ES_60 | Spain | Madrid | *hspSWEurope* | *hspSWEurope* | 1581875 | 21 | 38.2 | 1490 |
| 619 | EUR122 | ES_61 | Spain | Madrid | *hspAfrica1MiscAmerica* | *hspSWEuropeNorthAmerica* | 1669259 | 56 | 38.4 | 1551 |
| 620 | EUR123 | ES_610 | Spain | Madrid | *hspSWEurope* | *hspSWEurope* | 1622089 | 26 | 39.3 | 1530 |
| 621 | EUR124 | ES_68 | Spain | Madrid | *hspSWEurope* | *hspSWEurope* | 1588849 | 18 | 38.8 | 1480 |
| 622 | EUR125 | ES_7 | Spain | Madrid | *hspSWEurope* | *hspSWEurope* | 1576631 | 22 | 37.5 | 1493 |
| 623 | EUR126 | ES_76 | Spain | Madrid | *hspSWEurope* | *hspSWEurope* | 1621939 | 32 | 38.8 | 1512 |
| 624 | EUR127 | ES_DO | Spain | Madrid | *hspSWEurope* | *hspSWEurope* | 1618507 | 31 | 38.3 | 1524 |
| 625 | EUR128 | ES_SN | Spain | Madrid | *hspSWEuropeHonduras* | *hspSWEuropeSouthAmerica* | 1624670 | 33 | 38.1 | 1509 |
| 626 | EUR129 | HUP_B14 | Spain | Unknown | *hspSWEurope* | *hspSWEurope* | 1607584 | 2 | 37.8 | 1506 |
| 627 | EUR130 | HE_C1 | Sweden | Orebro | *hspNEurope* | *hspNEurope* | 1588871 | 75 | 38.3 | 1483 |
| 628 | EUR131 | HE_C18 | Sweden | Vasteras | *hspNEurope* | *hspNEurope* | 1635728 | 97 | 37.9 | 1519 |
| 629 | EUR132 | HE_C23 | Sweden | Vasteras | *hspNEurope* | *hspNEurope* | 1663959 | 100 | 37.8 | 1559 |
| 630 | EUR133 | HE_C30 | Sweden | Kalmar | *hspNEurope* | *hspNEurope* | 1657933 | 115 | 37.4 | 1555 |
| 631 | EUR134 | HE_C32 | Sweden | Goteborg | *hspNEurope* | *hspNEurope* | 1636505 | 83 | 37.9 | 1540 |
| 632 | EUR135 | HE_C33 | Sweden | Vasteras | *hspNEurope* | *hspNEurope* | 1593414 | 75 | 38.4 | 1488 |
| 633 | EUR136 | HE_C34 | Sweden | Vasteras | *hspNEurope* | *hspNEurope* | 1597206 | 107 | 38.1 | 1509 |
| 634 | EUR137 | HE_C38 | Sweden | Kalmar | *hspNEurope* | *hspNEurope* | 1619325 | 66 | 37.7 | 1528 |
| 635 | EUR138 | HE_C50 | Sweden | Goteborg | *hspNEurope* | *hspNEurope* | 1673940 | 46 | 38.6 | 1571 |
| 636 | EUR139 | HE_C52 | Sweden | Mora | *hspNEurope* | *hspNEurope* | 1601235 | 54 | 38.7 | 1505 |
| 637 | EUR140 | HE_C55 | Sweden | Kalmar | *hspNEurope* | *hspNEurope* | 1617593 | 125 | 38.4 | 1515 |
| 638 | EUR141 | HE_C66 | Sweden | Stockholm | *hspNEurope* | *hspNEurope* | 1667591 | 93 | 38.1 | 1546 |
| 639 | EUR142 | HE_NC1_1 | Sweden | Orebro | *hspNEurope* | *hspNEurope* | 1585003 | 116 | 38.2 | 1497 |
| 640 | EUR143 | HE_NC13_5 | Sweden | Kalmar | *hspNEurope* | *hspNEurope* | 1660984 | 225 | 38.7 | 1535 |
| 641 | EUR144 | HE_NC18_1 | Sweden | Vasteras | *hspSEurope* | *hspSEurope* | 1620332 | 150 | 37.4 | 1543 |
| 642 | EUR145 | HE_NC18_4 | Sweden | Vasteras | *hspNEurope* | *hspNEurope* | 1670041 | 100 | 38.3 | 1579 |
| 643 | EUR146 | HE_NC20_5 | Sweden | Kalmar | *hspNEurope* | *hspNEurope* | 1614952 | 136 | 38.1 | 1499 |
| 644 | EUR147 | HE_NC23_2a | Sweden | Vasteras | *hspNEurope* | *hspNEurope* | 1643502 | 76 | 38.0 | 1549 |
| 645 | EUR148 | HE_NC24_6 | Sweden | Kalmar | *hspNEurope* | *hspNEurope* | 1647215 | 74 | 37.8 | 1544 |
| 646 | EUR149 | HE_NC27_4 | Sweden | Kalmar | *hspNEurope* | *hspNEurope* | 1601303 | 82 | 38.2 | 1504 |
| 647 | EUR150 | HE_NC29_2 | Sweden | Orebro | *hspNEurope* | *hspNEurope* | 1592465 | 82 | 38.6 | 1503 |
| 648 | EUR151 | HE_NC30_2 | Sweden | Kalmar | *hspNEurope* | *hspNEurope* | 1555156 | 96 | 38.0 | 1462 |
| 649 | EUR152 | HE_NC30_3 | Sweden | Kalmar | *hspNEurope* | *hspNEurope* | 1617715 | 111 | 37.6 | 1514 |
| 650 | EUR153 | HE_NC32_4 | Sweden | Goteborg | *hspNEurope* | *hspNEurope* | 1625295 | 88 | 37.6 | 1521 |
| 651 | EUR154 | HE_NC32_5 | Sweden | Goteborg | *hspNEurope* | *hspNEurope* | 1599686 | 74 | 38.5 | 1490 |
| 652 | EUR155 | HE_NC36_4 | Sweden | Kalmar | *hspNEurope* | *hspNEurope* | 1614521 | 80 | 37.6 | 1527 |
| 653 | EUR156 | HE_NC38_2 | Sweden | Kalmar | *hspNEurope* | *hspNEurope* | 1616978 | 60 | 38.4 | 1528 |
| 654 | EUR157 | HE_NC38_4 | Sweden | Kalmar | *hspNEurope* | *hspNEurope* | 1643242 | 91 | 38.1 | 1550 |
| 655 | EUR158 | HE_NC38_5 | Sweden | Kalmar | *hspNEurope* | *hspNEurope* | 1613524 | 66 | 38.3 | 1507 |
| 656 | EUR159 | HE_NC47_5 | Sweden | Vasteras | *hspNEurope* | *hspNEurope* | 1642370 | 86 | 37.5 | 1542 |
| 657 | EUR160 | HE_NC5_3 | Sweden | Kalmar | *hspNEurope* | *hspNEurope* | 1591808 | 60 | 38.7 | 1484 |
| 658 | EUR161 | HE_NC55_1 | Sweden | Kalmar | *hspNEurope* | *hspNEurope* | 1630581 | 71 | 38.2 | 1531 |
| 659 | EUR162 | HE_NC55_2 | Sweden | Kalmar | *hspNEurope* | *hspNEurope* | 1600232 | 55 | 38.7 | 1505 |
| 660 | EUR163 | HE_NC60_1 | Sweden | Kalmar | *hspNEurope* | *hspNEurope* | 1627305 | 59 | 37.8 | 1547 |
| 661 | EUR164 | HE_NC60_3 | Sweden | Kalmar | *hspNEurope* | *hspNEurope* | 1545648 | 57 | 38.7 | 1462 |
| 662 | EUR165 | HE_NC61_4 | Sweden | Kalmar | *hspNEurope* | *hspNEurope* | 1598197 | 61 | 38.6 | 1528 |
| 663 | EUR166 | HE_NC9_1 | Sweden | Kalmar | *hspNEurope* | *hspNEurope* | 1573353 | 64 | 38.8 | 1487 |
| 664 | EUR167 | HPAG1 | Sweden | Kalixanda | *hspNEurope* | *hspNEurope* | 1605736 | 2 | 37.7 | 1506 |
| 665 | EUR168 | Sw577G | Sweden | Unknown | *hspNEurope* | *hspNEurope* | 1654799 | 48 | 38.4 | 1565 |
| 666 | EUR169 | SwA626G | Sweden | Unknown | *hspNEurope* | *hspNEurope* | 1676675 | 29 | 38.5 | 1597 |
| 667 | EUR170 | 444 | UK | Nottingham | *hspNEurope* | *hspNEurope* | 1613498 | 44 | 39.1 | 1516 |
| 668 | EUR171 | 448 | UK | Nottingham | *hspNEurope* | *hspNEurope* | 1632931 | 57 | 37.8 | 1543 |
| 669 | EUR172 | 456 | UK | Nottingham | *hspNEurope* | *hspNEurope* | 1599501 | 44 | 38.4 | 1514 |
| 670 | EUR173 | 518 | UK | Nottingham | *hspNEurope* | *hspNEurope* | 1623419 | 47 | 38.1 | 1559 |
| 671 | EUR174 | 638 | UK | Nottingham | *hspNEurope* | *hspNEurope* | 1583926 | 42 | 38.3 | 1480 |
| 672 | EUR175 | 26695_Tomb | UK | Unknown | *hspNEurope* | *hspNEurope* | 1667892 | 1 | 38.9 | 1575 |
| 673 | EUR176 | H3014 | UK | Coventry | *hspNEurope* | *hspNEurope* | 1641212 | 44 | 38.3 | 1544 |
| 674 | EUR177 | SW21C | UK | Swansea | *hspNEurope* | *hspNEurope* | 1620810 | 57 | 37.4 | 1520 |
| 675 | EUR178 | SW23Ci | UK | Swansea | *hspNEurope* | *hspSEurope* | 1684861 | 65 | 38.0 | 1580 |
| 676 | EUR179 | SW7C | UK | Swansea | *hspNEurope* | *hspNEurope* | 1643252 | 64 | 38.2 | 1549 |
| 677 | EUR180 | UKEN31U | UK | Unknown | *hspNEurope* | *hspNEurope* | 1642283 | 54 | 38.8 | 1547 |
| 678 | EUR181 | altai18 | Russia | Altai | *hpEurope* | *hspAmerind* | 1620701 | 67 | 38.9 | 1543 |
| 679 | EUR182 | altai45 | Russia | Altai | *hpEurope* | *hspAmerind* | 1648327 | 55 | 38.3 | 1546 |
| 680 | EUR183 | altai59 | Russia | Altai | *hspSiberia1* | *hspAmerind* | 1641363 | 65 | 37.9 | 1539 |
| 681 | EUR184 | BURYAT14 | Russia | UlanUlde | *hspAltai* | *hspAmerind* | 1613560 | 71 | 38.0 | 1516 |
| 682 | EUR185 | BURYAT19 | Russia | UlanUlde | *hspSiberia1* | *hspAmerind* | 1671230 | 77 | 37.5 | 1576 |
| 683 | EUR186 | BURYAT2 | Russia | UlanUlde | *hspAltai* | *hspAmerind* | 1557240 | 104 | 38.9 | 1451 |
| 684 | EUR187 | BURYAT27 | Russia | UlanUlde | *hspSiberia1* | *hspAmerind* | 1615584 | 59 | 38.5 | 1502 |
| 685 | EUR188 | BURYAT9 | Russia | UlanUlde | *hspAltai* | *hspAmerind* | 1537480 | 44 | 38.5 | 1452 |
| 686 | EUR189 | Chukchi03 | Russia | Bilibino | *hspAmerind* | *hspAmerind* | 1491039 | 56 | 40.1 | 1377 |
| 687 | EUR190 | Chukchi08 | Russia | Bilibino | *hspSiberia* | *hspAmerind* | 1676389 | 66 | 38.5 | 1562 |
| 688 | EUR191 | Chukchi09 | Russia | Bilibino | *hpEurope* | *hspSEurope* | 1625793 | 64 | 37.8 | 1506 |
| 689 | EUR192 | Chukchi30 | Russia | Bilibino | *hpEurope* | *hspSEurope* | 1654884 | 52 | 38.5 | 1559 |
| 690 | EUR193 | Even14 | Russia | Anavgay | *hspSiberia1* | *hspAmerind* | 1634954 | 92 | 36.5 | 1540 |
| 691 | EUR194 | Even33 | Russia | Anavgay | *hspAmerind* | *hspAmerind* | 1646471 | 67 | 37.6 | 1525 |
| 692 | EUR195 | Even42 | Russia | Anavgay | *hspAmerind* | *hspAmerind* | 1623036 | 60 | 38.0 | 1505 |
| 693 | EUR196 | Evenky01 | Russia | Yessei | *hspSiberia1* | *hspAmerind* | 1635640 | 89 | 38.1 | 1524 |
| 694 | EUR197 | Evenky05 | Russia | Tura | *hspAltai* | *hspAmerind* | 1583664 | 76 | 37.9 | 1461 |
| 695 | EUR198 | Evenky11 | Russia | Tutonga | *hspAmerind* | *hspAmerind* | 1639970 | 76 | 37.9 | 1516 |
| 696 | EUR199 | Evenky58 | Russia | Tura | *hspAmerind* | *hspAmerind* | 1645885 | 109 | 38.1 | 1532 |
| 697 | EUR200 | Evenky65 | Russia | Tura | *hspSiberia1* | *hspAmerind* | 1619445 | 88 | 38.3 | 1492 |
| 698 | EUR201 | Evenky73 | Russia | Chirinda | *hspSiberia1* | *hspAmerind* | 1623568 | 134 | 38.5 | 1512 |
| 699 | EUR202 | Keto60 | Russia | Sulomai | *hspKet* | *hspAmerind* | 1639355 | 93 | 38.1 | 1534 |
| 700 | EUR203 | Khanty04 | Russia | Shurishkari | *hspAmerind* | *hspAmerind* | 1610205 | 57 | 38.0 | 1496 |
| 701 | EUR204 | Khanty16 | Russia | Shurishkari | *hspUral* | *hspAmerind* | 1639656 | 61 | 38.6 | 1544 |
| 702 | EUR205 | Khanty17 | Russia | Shurishkari | *hspUral* | *hspAmerind* | 1667542 | 79 | 38.6 | 1562 |
| 703 | EUR206 | Khanty26 | Russia | Shurishkari | *hspUral* | *hspAmerind* | 1635606 | 54 | 38.2 | 1541 |
| 704 | EUR207 | Khanty27 | Russia | Shurishkari | *hspSiberia1* | *hspAmerind* | 1597145 | 47 | 38.6 | 1506 |
| 705 | EUR208 | Khanty30 | Russia | Shurishkari | *hspUral* | *hspAmerind* | 1626735 | 63 | 38.3 | 1525 |
| 706 | EUR209 | Khanty39 | Russia | Shurishkari | *hspUral* | *hspAmerind* | 1647468 | 57 | 37.2 | 1550 |
| 707 | EUR210 | Khanty40 | Russia | Shurishkari | *hspUral* | *hspAmerind* | 1585843 | 59 | 39.5 | 1492 |
| 708 | EUR211 | Khanty47 | Russia | Shurishkari | *hspSiberia2* | *hspAmerind* | 1522983 | 46 | 38.9 | 1426 |
| 709 | EUR212 | Khanty48 | Russia | Shurishkari | *hpEurope* | *hspSEurope* | 1602221 | 58 | 38.4 | 1492 |
| 710 | EUR213 | Koryak28 | Russia | Shurishkari | *hspSiberia1* | *hspAmerind* | 1558583 | 80 | 37.6 | 1449 |
| 711 | EUR214 | Koryak35 | Russia | Palana | *hspAmerind* | *hspAmerind* | 1637456 | 59 | 37.8 | 1537 |
| 712 | EUR215 | Koryak37 | Russia | Palana | *hspAmerind* | *hspAmerind* | 1551573 | 57 | 38.9 | 1432 |
| 713 | EUR216 | Nanai27 | Russia | Naichin | *hpEurope* | *hspSEurope* | 1662438 | 68 | 37.9 | 1552 |
| 714 | EUR217 | Nanai30 | Russia | Naichin | *hspSiberia2* | *hspAmerind* | 1644088 | 76 | 37.8 | 1552 |
| 715 | EUR218 | Nanai34 | Russia | Naichin | *hspAmerind* | *hspAmerind* | 1644459 | 72 | 37.2 | 1535 |
| 716 | EUR219 | Nanai40 | Russia | Naichin | *hspAmerind* | *hspAmerind* | 1671876 | 73 | 38.1 | 1537 |
| 717 | EUR220 | Nentsy28 | Russia | NovyPort | *hspUral* | *hspAmerind* | 1677487 | 65 | 37.6 | 1568 |
| 718 | EUR221 | Nentsy29 | Russia | NovyPort | *hspUral* | *hspAmerind* | 1622982 | 36 | 38.1 | 1521 |
| 719 | EUR222 | Nentsy32 | Russia | NovyPort | *hspUral* | *hspAmerind* | 1624378 | 43 | 38.1 | 1528 |
| 720 | EUR223 | Nentsy37 | Russia | NovyPort | *hspUral* | *hspAmerind* | 1669500 | 66 | 37.1 | 1548 |
| 721 | EUR224 | Nentsy38 | Russia | NovyPort | *hspUral* | *hspAmerind* | 1646872 | 61 | 37.9 | 1540 |
| 722 | EUR225 | Nentsy42 | Russia | NovyPort | *hpAsia2* | *hspAmerind* | 1587324 | 116 | 38.9 | 1508 |
| 723 | EUR226 | Nentsy65 | Russia | NovyPort | *hspUral* | *hspAmerind* | 1634949 | 51 | 37.6 | 1530 |
| 724 | EUR227 | Nentsy81 | Russia | NovyPort | *hspAmerind* | *hspAmerind* | 1601626 | 56 | 38.7 | 1490 |
| 725 | EUR228 | Nentsy90 | Russia | NovyPort | *hspUral* | *hspAmerind* | 1629897 | 68 | 38.2 | 1526 |
| 726 | EUR229 | Sakhalin26 | Russia | Nogliki | *hpEurope* | *hspAmerind* | 1632359 | 88 | 38.4 | 1516 |
| 727 | EUR230 | Sakhalin60 | Russia | Val | *hspAmerind* | *hspAmerind* | 1671335 | 104 | 38.1 | 1532 |
| 728 | EUR231 | Sakhalin89 | Russia | Val | *hpEurope* | *hspSEurope* | 1648493 | 73 | 38.5 | 1548 |
| 729 | EUR232 | TUVAB15 | Russia | Kyzyl | *hspSiberia2* | *hspAmerind* | 1637575 | 65 | 37.7 | 1544 |
| 730 | EUR233 | TUVAB18a | Russia | Kyzyl | *hspAltai* | *hspAmerind* | 1614459 | 44 | 38.9 | 1514 |
| 731 | EUR234 | Tuvac16 | Russia | Kyzyl | *hpEurope* | *hspSEurope* | 1608037 | 191 | 38.0 | 1506 |
| 732 | EUR235 | Tuvac41 | Russia | Kyzyl | *hspLadak* | *hspAmerind* | 1630169 | 62 | 38.0 | 1539 |
| 733 | EUR236 | Tuvac46 | Russia | Kyzyl | *hspSiberia2* | *hspAmerind* | 1627303 | 54 | 37.9 | 1526 |
| 734 | EUR237 | Tuvac70 | Russia | Kyzyl | *hspSiberia1* | *hspAmerind* | 1718395 | 65 | 37.6 | 1619 |
| 735 | EUR238 | Tuvac80 | Russia | Kyzyl | *hspSiberia2* | *hspAmerind* | 1698000 | 64 | 37.9 | 1589 |
| 736 | EUR239 | Ulchi51 | Russia | Ulchi | *hpEurope* | *hspSEurope* | 1599184 | 41 | 38.9 | 1503 |
| 737 | EUR240 | Ulchi52 | Russia | Ulchi | *hpEurope* | *hspSEurope* | 1637246 | 43 | 39.4 | 1527 |
| 738 | EUR241 | yak97 | Russia | Yakutsk | *hspSiberia1* | *hspAmerind* | 1626090 | 55 | 38.2 | 1540 |
| 739 | EUR242 | BON254 | Unknown | NorthernEurope | *hspSEurope* | *hspSEurope* | 1659275 | 49 | 38.1 | 1581 |
| 740 | EUR243 | A45 | Russia | Moscow | *hspSAmerind* | *hspAmerind* | 1643927 | 33 | 37.7 | 1553 |
| 741 | EUR246 | Iceman | Italia | Eastern_Italian_Alps | *hpEurope* | *hspNEurope* | 1667867 | 1 | 39.2 | 1415 |
| 742 | NAM001 | Aklavik117 | Canada | NorthwestTerritories | *hspNAmerind* | *hspAmerind* | 1636125 | 3 | 36.1 | 1530 |
| 743 | NAM002 | Aklavik86 | Canada | NorthwestTerritories | *hspNAmerind* | *hspAmerind* | 1507930 | 3 | 36.6 | 1412 |
| 744 | NAM003 | Ca_97_30 | Canada | Unknown | *hspNAmerind* | *hspAmerind* | 1535086 | 44 | 37.3 | 1447 |
| 745 | NAM004 | Ca_97_34 | Canada | Unknown | *hspNAmerind* | *hspAmerind* | 1522915 | 33 | 38.5 | 1421 |
| 746 | NAM005 | CA-A105 | Canada | Unknown | *hspNAmerind* | *hspSWEuropeNorthAmerica* | 1636198 | 44 | 38.4 | 1519 |
| 747 | NAM006 | CA-A167 | Canada | NorthwestTerritories | *hspAfrica1MiscAmerica* | *hspAmerind* | 1501639 | 35 | 38.9 | 1412 |
| 748 | NAM007 | CA-A17 | Canada | NorthwestTerritories | *hspNAmerind* | *hspAmerind* | 1499829 | 33 | 39.0 | 1412 |
| 749 | NAM008 | CA-A212 | Canada | NorthwestTerritories | *hspNAmerind* | *hspAmerind* | 1502507 | 33 | 39.2 | 1412 |
| 750 | NAM009 | CA-A44 | Canada | NorthwestTerritories | *hspNAmerind* | *hspAmerind* | 1516681 | 41 | 38.5 | 1426 |
| 751 | NAM010 | CA-A46 | Canada | NorthwestTerritories | *hspNAmerind* | *hspAmerind* | 1483326 | 44 | 38.6 | 1379 |
| 752 | NAM011 | CA-A52 | Canada | NorthwestTerritories | *hspNAmerind* | *hspAmerind* | 1522716 | 44 | 38.2 | 1430 |
| 753 | NAM012 | CA-A53 | Canada | NorthwestTerritories | *hspNAmerind* | *hspAmerind* | 1513431 | 33 | 38.6 | 1413 |
| 754 | NAM013 | CA-A54 | Canada | NorthwestTerritories | *hspNAmerind* | *hspAmerind* | 1515893 | 37 | 38.1 | 1415 |
| 755 | NAM014 | CA-A63 | Canada | NorthwestTerritories | *hspNAmerind* | *hspAmerind* | 1602555 | 53 | 37.9 | 1513 |
| 756 | NAM015 | CA-A7 | Canada | NorthwestTerritories | *hspNAmerind* | *hspAmerind* | 1499994 | 34 | 39.2 | 1411 |
| 757 | NAM016 | CA-A73 | Canada | NorthwestTerritories | *hspNAmerind* | *hspAmerind* | 1517865 | 32 | 38.8 | 1433 |
| 758 | NAM017 | CA-A78 | Canada | NorthwestTerritories | *hspNAmerind* | *hspAmerind* | 1523994 | 14 | 39.0 | 1425 |
| 759 | NAM018 | CA-A88 | Canada | NorthwestTerritories | *hspNAmerind* | *hspAmerind* | 1516420 | 39 | 38.6 | 1426 |
| 760 | NAM019 | CCHI33 | Canada | Unknown | *Unknown* | *hspAfrica1NAmerica* | 1659327 | 11 | 39.0 | 1540 |
| 761 | NAM020 | R018c | Canada | Alberta | *hspNEurope* | *hspNEurope* | 1646836 | 15 | 39.3 | 1555 |
| 762 | NAM021 | R030b | Canada | Alberta | *hspAfrica1SAfrica* | *hspAfrica1SAfrica* | 1610617 | 11 | 40.5 | 1512 |
| 763 | NAM022 | R036d | Canada | Alberta | *hspSEurope* | *hspSEurope* | 1640262 | 8 | 39.1 | 1542 |
| 764 | NAM023 | R037c | Canada | Alberta | *hspNEurope* | *hspNEurope* | 1614130 | 9 | 38.9 | 1528 |
| 765 | NAM024 | R038b | Canada | Alberta | *hspNEurope* | *hspNEurope* | 1632757 | 14 | 38.8 | 1521 |
| 766 | NAM025 | R046Wa | Canada | Alberta | *hspSEurope* | *hspSEurope* | 1585443 | 8 | 38.9 | 1502 |
| 767 | NAM026 | R055a | Canada | Alberta | *hspSEurope* | *hspSEurope* | 1631090 | 16 | 38.5 | 1539 |
| 768 | NAM027 | R32b | Canada | Alberta | *hspNEurope* | *hspNEurope* | 1582461 | 10 | 38.5 | 1504 |
| 769 | NAM028 | UMB-G1 | Canada | Unknown | *hspSWEurope* | *hspSWEurope* | 1571801 | 5 | 43.2 | 1471 |
| 770 | NAM029 | CR_15 | CostaRica | Unknown | *hspSWEuropeMexico* | *hspSWEuropeSouthAmerica* | 1648440 | 38 | 38.7 | 1536 |
| 771 | NAM030 | CR_16 | CostaRica | Unknown | *hspAfrica1NAmerica* | *hspAfrica1NAmerica* | 1597089 | 28 | 38.7 | 1488 |
| 772 | NAM031 | CR_9 | CostaRica | Unknown | *hspSWEuropeMexico* | *hspSWEuropeSouthAmerica* | 1679356 | 30 | 37.0 | 1545 |
| 773 | NAM033 | HpA_11 | USA | Cleveland | *hspSEurope* | *hspSEurope* | 1668342 | 1 | 38.8 | 1588 |
| 774 | NAM034 | HpA_14 | USA | Cleveland | *hspSEurope* | *hspSEurope* | 1594597 | 7 | 38.9 | 1502 |
| 775 | NAM035 | HpA_16 | USA | Cleveland | *hspAfrica1NAmerica* | *hspAfrica1NAmerica* | 1637800 | 11 | 38.6 | 1536 |
| 776 | NAM036 | HpA_17 | USA | Cleveland | *hspAfrica1NAmerica* | *hspAfrica1NAmerica* | 1636618 | 5 | 41.1 | 1532 |
| 777 | NAM037 | HpA_20 | USA | Cleveland | *hspAfrica1NAmerica* | *hspAfrica1NAmerica* | 1674600 | 10 | 38.9 | 1589 |
| 778 | NAM038 | HpA_26 | USA | Cleveland | *hspSWEurope* | *hspSEurope* | 1624194 | 9 | 37.2 | 1553 |
| 779 | NAM039 | HpA_27 | USA | Cleveland | *hspSEurope* | *hspSEurope* | 1647774 | 6 | 38.9 | 1581 |
| 780 | NAM040 | HpA_4 | USA | Cleveland | *hspAfrica1WAfrica* | *hspAfrica1NAmerica* | 1663998 | 8 | 39.2 | 1567 |
| 781 | NAM041 | HpA_5 | USA | Cleveland | *hspAfrica1NAmerica* | *hspAfrica1NAmerica* | 1635925 | 7 | 39.3 | 1543 |
| 782 | NAM042 | HpA_6 | USA | Cleveland | *hspAfrica1NAmerica* | *hspAfrica1NAmerica* | 1653378 | 12 | 39.1 | 1541 |
| 783 | NAM043 | HpA_8 | USA | Cleveland | *hspAfrica1NAmerica* | *hspAfrica1NAmerica* | 1640628 | 7 | 38.6 | 1529 |
| 784 | NAM044 | HpA_9 | USA | Cleveland | *hspSEurope* | *hspSEurope* | 1714927 | 17 | 37.7 | 1626 |
| 785 | NAM045 | HpH_1 | USA | Cleveland | *hspAfrica1NAmerica* | *hspAfrica1NAmerica* | 1682934 | 54 | 38.4 | 1553 |
| 786 | NAM046 | HpH_10 | USA | Cleveland | *hspAfrica1NAmerica* | *hspAfrica1NAmerica* | 1649818 | 32 | 38.0 | 1521 |
| 787 | NAM047 | HpH_11 | USA | Cleveland | *hspSEurope* | *hspSEurope* | 1657667 | 9 | 38.9 | 1555 |
| 788 | NAM048 | HpH_16 | USA | Cleveland | *hspAfrica1WAfrica* | *hspAfrica1WAfrica* | 1703959 | 8 | 37.7 | 1560 |
| 789 | NAM049 | HpH_18 | USA | Cleveland | *hspAfrica1NAmerica* | *hspAfrica1NAmerica* | 1760691 | 21 | 37.6 | 1667 |
| 790 | NAM050 | HpH_19 | USA | Cleveland | *hspAfrica1NAmerica* | *hspAfrica1NAmerica* | 1634366 | 7 | 38.6 | 1511 |
| 791 | NAM051 | HpH_21 | USA | Cleveland | *hspAfrica1NAmerica* | *hspAfrica1NAmerica* | 1623667 | 6 | 38.1 | 1509 |
| 792 | NAM052 | HpH_23 | USA | Cleveland | *hspAfrica1NAmerica* | *hspAfrica1NAmerica* | 1648064 | 9 | 39.1 | 1535 |
| 793 | NAM053 | HpH_24 | USA | Cleveland | *hspAfrica1NAmerica* | *hspAfrica1NAmerica* | 1670182 | 8 | 38.5 | 1561 |
| 794 | NAM054 | HpH_27 | USA | Cleveland | *hspSEurope* | *hspSEurope* | 1612044 | 5 | 39.0 | 1527 |
| 795 | NAM055 | HpH_28 | USA | Cleveland | *hspNEurope* | *hspNEurope* | 1625000 | 16 | 37.4 | 1537 |
| 796 | NAM056 | HpH_29 | USA | Cleveland | *hspAfrica1NAmerica* | *hspAfrica1NAmerica* | 1677607 | 16 | 38.2 | 1529 |
| 797 | NAM057 | HpH_3 | USA | Cleveland | *hspAfrica1NAmerica* | *hspAfrica1NAmerica* | 1712042 | 11 | 38.4 | 1554 |
| 798 | NAM058 | HpH_30 | USA | Cleveland | *hspAfrica1NAmerica* | *hspAfrica1NAmerica* | 1625395 | 11 | 38.6 | 1501 |
| 799 | NAM059 | HpH_34 | USA | Cleveland | *hspAfrica1NAmerica* | *hspAfrica1NAmerica* | 1627237 | 8 | 39.3 | 1529 |
| 800 | NAM060 | HpH_36 | USA | Cleveland | *hspAfrica1NAmerica* | *hspAfrica1NAmerica* | 1675907 | 16 | 38.6 | 1559 |
| 801 | NAM061 | HpH_4 | USA | Cleveland | *hspAfrica1NAmerica* | *hspAfrica1NAmerica* | 1665093 | 9 | 38.4 | 1559 |
| 802 | NAM062 | HpH_41 | USA | Cleveland | *hspAfrica1NAmerica* | *hspAfrica1NAmerica* | 1660567 | 11 | 38.8 | 1548 |
| 803 | NAM063 | HpH_42 | USA | Cleveland | *hspAfrica1NAmerica* | *hspAfrica1NAmerica* | 1702292 | 19 | 39.0 | 1568 |
| 804 | NAM064 | HpH_43 | USA | Cleveland | *hspSWEurope* | *hspSEurope* | 1607665 | 12 | 38.9 | 1501 |
| 805 | NAM065 | HpH_44 | USA | Cleveland | *hspAfrica1NAmerica* | *hspAfrica1NAmerica* | 1665840 | 16 | 38.4 | 1538 |
| 806 | NAM066 | HpH_45 | USA | Cleveland | *hspSEurope* | *hspSEurope* | 1655397 | 14 | 38.9 | 1560 |
| 807 | NAM067 | HpH_5b | USA | Cleveland | *hspAfrica1NAmerica* | *hspAfrica1NAmerica* | 1708533 | 11 | 38.6 | 1586 |
| 808 | NAM068 | HpH_6 | USA | Cleveland | *hspAfrica1MiscAmerica* | *hspSWEuropeNorthAmerica* | 1707985 | 10 | 37.7 | 1613 |
| 809 | NAM069 | HpH_9 | USA | Cleveland | *hspSEurope* | *hspSEurope* | 1637692 | 11 | 38.7 | 1530 |
| 810 | NAM070 | HpP_1 | USA | Cleveland | *hspAfrica1NAmerica* | *hspAfrica1NAmerica* | 1669637 | 6 | 38.9 | 1553 |
| 811 | NAM071 | HpP_11 | USA | Cleveland | *hspAfrica1NAmerica* | *hspAfrica1NAmerica* | 1689122 | 11 | 38.9 | 1605 |
| 812 | NAM072 | HpP_13 | USA | Cleveland | *hspAfrica1NAmerica* | *hspAfrica1NAmerica* | 1714414 | 10 | 37.9 | 1661 |
| 813 | NAM073 | HpP_15 | USA | Cleveland | *hspSEurope* | *hspSEurope* | 1652821 | 4 | 38.9 | 1560 |
| 814 | NAM074 | HpP_16 | USA | Cleveland | *hspSEurope* | *hspSEurope* | 1549132 | 5 | 39.1 | 1468 |
| 815 | NAM075 | HpP_2 | USA | Cleveland | *hspAfrica1NAmerica* | *hspAfrica1NAmerica* | 1693870 | 6 | 39.1 | 1570 |
| 816 | NAM076 | HpP_23 | USA | Cleveland | *hspSEurope* | *hspSEurope* | 1643525 | 3 | 38.7 | 1582 |
| 817 | NAM077 | HpP_25d | USA | Cleveland | *hspAfrica1WAfrica* | *hspAfrica1WAfrica* | 1657665 | 9 | 39.3 | 1543 |
| 818 | NAM078 | HpP_26 | USA | Cleveland | *hspAfrica1NAmerica* | *hspAfrica1NAmerica* | 1697203 | 9 | 38.6 | 1589 |
| 819 | NAM079 | HpP_28b | USA | Cleveland | *hspAfrica1NAmerica* | *hspAfrica1NAmerica* | 1619904 | 11 | 39.6 | 1522 |
| 820 | NAM080 | HpP_3 | USA | Cleveland | *hspAfrica1NAmerica* | *hspAfrica1NAmerica* | 1652009 | 10 | 39.3 | 1544 |
| 821 | NAM081 | HpP_30 | USA | Cleveland | *hspSEurope* | *hspSEurope* | 1639810 | 4 | 38.8 | 1540 |
| 822 | NAM082 | J166 | USA | Nashville | *hspSEurope* | *hspSEurope* | 1650561 | 1 | 38.9 | 1538 |
| 823 | NAM083 | J99 | USA | Nashville | *hspAfrica1NAmerica* | *hspAfrica1NAmerica* | 1643831 | 1 | 39.2 | 1504 |
| 824 | NAM084 | NYPt127 | USA | NewYorkCity | *hspAfrica1NAmerica* | *hspAfrica1NAmerica* | 1642811 | 43 | 39.5 | 1513 |
| 825 | NAM085 | NYPt130 | USA | NewYorkCity | *hspAfrica1NAmerica* | *hspAfrica1NAmerica* | 1679649 | 63 | 38.4 | 1561 |
| 826 | NAM086 | NYPt133 | USA | NewYorkCity | *hspAfrica1NAmerica* | *hspAfrica1NAmerica* | 1644857 | 76 | 39.0 | 1517 |
| 827 | NAM087 | NYPt154 | USA | NewYorkCity | *hspAfrica1NAmerica* | *hspAfrica1NAmerica* | 1643368 | 68 | 38.8 | 1524 |
| 828 | NAM088 | NYPt170 | USA | NewYorkCity | *hspSEurope* | *hspSEurope* | 1655109 | 62 | 38.8 | 1560 |
| 829 | NAM089 | NYPt186 | USA | NewYorkCity | *hspSWEurope* | *hspSWEurope* | 1605103 | 58 | 39.9 | 1502 |
| 830 | NAM090 | NYPt187 | USA | NewYorkCity | *hspAfrica1WAfrica* | *hspAfrica1NAmerica* | 1704164 | 70 | 38.1 | 1572 |
| 831 | NAM091 | NYPt189 | USA | NewYorkCity | *hspAfrica1NAmerica* | *hspAfrica1NAmerica* | 1636344 | 57 | 38.9 | 1527 |
| 832 | NAM092 | NYPt19 | USA | NewYorkCity | *hspSEurope* | *hspSEurope* | 1583029 | 42 | 39.2 | 1503 |
| 833 | NAM093 | NYPt273 | USA | NewYorkCity | *hspAfrica1NAmerica* | *hspAfrica1NAmerica* | 1627774 | 53 | 41.4 | 1502 |
| 834 | NAM094 | NYPt281 | USA | NewYorkCity | *hspSWEuropeMexico* | *hspSWEuropeSouthAmerica* | 1667182 | 79 | 37.5 | 1551 |
| 835 | NAM095 | NYPt292 | USA | NewYorkCity | *hspAfrica1WAfrica* | *hspAfrica1NAmerica* | 1682411 | 55 | 38.8 | 1556 |
| 836 | NAM096 | NYPt297 | USA | NewYorkCity | *hspSWEurope* | *hspSWEuropeSouthAmerica* | 1621890 | 90 | 39.1 | 1507 |
| 837 | NAM097 | NYPt310 | USA | NewYorkCity | *hspAfrica1NAmerica* | *hspAfrica1NAmerica* | 1647461 | 51 | 39.4 | 1524 |
| 838 | NAM098 | NYPt328 | USA | NewYorkCity | *hspNEurope* | *hspNEurope* | 1644337 | 76 | 40.1 | 1565 |
| 839 | NAM099 | NYPt64 | USA | NewYorkCity | *hspSEurope* | *hspSEurope* | 1620535 | 35 | 39.1 | 1525 |
| 840 | NAM100 | NYPt66 | USA | NewYorkCity | *hspNEurope* | *hspNEurope* | 1542137 | 40 | 40.6 | 1440 |
| 841 | NAM101 | NYPt72 | USA | NewYorkCity | *hspNEurope* | *hspNEurope* | 1572649 | 48 | 38.2 | 1468 |
| 842 | NAM102 | NYPt82 | USA | NewYorkCity | *hspAfrica1NAmerica* | *hspAfrica1NAmerica* | 1616991 | 53 | 39.4 | 1519 |
| 843 | NAM103 | NYPt94 | USA | NewYorkCity | *hspNEurope* | *hspNEurope* | 1636543 | 51 | 38.6 | 1550 |
| 844 | NAM104 | NYPt95 | USA | NewYorkCity | *hspAfrica1NAmerica* | *hspAfrica1NAmerica* | 1698903 | 41 | 39.0 | 1574 |
| 845 | NAM105 | HpP_4 | USA | Cleveland | *hspAfrica1WAfrica* | *hspAfrica1NAmerica* | 1693924 | 8 | 38.6 | 1564 |
| 846 | NAM106 | HpP_41 | USA | Cleveland | *hspAfrica1NAmerica* | *hspAfrica1WAfrica* | 1712112 | 9 | 38.7 | 1597 |
| 847 | NAM107 | HpP_62 | USA | Cleveland | *hspAfrica1MiscAmerica* | *hspSWEuropeNorthAmerica* | 1648336 | 12 | 38.9 | 1525 |
| 848 | NAM108 | HpP_74 | USA | Cleveland | *hspSEurope* | *hspSEurope* | 1622385 | 2 | 38.9 | 1523 |
| 849 | NAM109 | HpP_8 | USA | Cleveland | *hspAfrica1WAfrica* | *hspAfrica1NAmerica* | 1615626 | 7 | 40.5 | 1512 |
| 850 | NAM110 | ALA10 | USA | Unknown | *hspAmerind* | *hspAmerind* | 1541449 | 75 | 37.6 | 1437 |
| 851 | NAM111 | ALA15 | USA | Alaska | *hspAmerind* | *hspAmerind* | 1685475 | 169 | 38.8 | 1588 |
| 852 | NAM112 | Ala2 | USA | Alaska | *hspAmerind* | *hspAmerind* | 1584890 | 82 | 37.6 | 1479 |
| 853 | NAM113 | ALA3 | USA | Alaska | *hspAmerind* | *hspAmerind* | 1663030 | 55 | 38.4 | 1556 |
| 854 | NAM114 | inma11 | Canada | NorthwestTerritories | *hspAmerind* | *hspAmerind* | 1528746 | 61 | 38.2 | 1437 |
| 855 | NAM115 | inma14 | Canada | NorthwestTerritories | *hspAmerind* | *hspAmerind* | 1518584 | 44 | 38.9 | 1412 |
| 856 | NAM116 | inma15 | Canada | NorthwestTerritories | *hspAmerind* | *hspAmerind* | 1519353 | 67 | 38.3 | 1417 |
| 857 | NAM117 | 29CaP | Mexico | Unknown | *hspSWEurope* | *hspSWEuropeSouthAmerica* | 1667159 | 1 | 38.8 | 1600 |
| 858 | NAM118 | 7C | Mexico | Unknown | *hspSWEurope* | *hspSWEurope* | 1631276 | 2 | 37.8 | 1530 |
| 859 | NAM119 | CG_IMSS_2012 | Mexico | Unknown | *hspSWEurope* | *hspSWEuropeSouthAmerica* | 1599050 | 45 | 38.4 | 1581 |
| 860 | NAM121 | C_Mx_2010_12 | Mexico | MexicoCity | *hspSWEuropeMexico* | *hspSWEuropeSouthAmerica* | 1658253 | 1 | 39.0 | 1552 |
| 861 | NAM122 | C_Mx_2010_97 | Mexico | MexicoCity | *hspAfrica1MiscAmerica* | *hspAfrica1SAfrica* | 1624459 | 1 | 39.2 | 1557 |
| 862 | NAM123 | C_Mx_2011_145 | Mexico | MexicoCity | *hspAfrica1MiscAmerica* | *hspSWEuropeNorthAmerica* | 1692846 | 2 | 37.6 | 1583 |
| 863 | NAM124 | C_Mx_2010_3 | Mexico | MexicoCity | *hspSWEuropeMexico* | *hspSWEuropeSouthAmerica* | 1642343 | 1 | 39.1 | 1539 |
| 864 | NAM125 | G_Mx_2003_250 | Mexico | MexicoCity | *hspAfrica1MiscAmerica* | *hspSWEuropeNorthAmerica* | 1672280 | 1 | 38.9 | 1558 |
| 865 | NAM126 | G_Mx_2003_108 | Mexico | MexicoCity | *hspSWEuropeMexico* | *hspSWEuropeSouthAmerica* | 1686171 | 2 | 36.8 | 1574 |
| 866 | NAM127 | G_Mx_2005_108 | Mexico | MexicoCity | *hspSWEuropeHonduras* | *hspSWEuropeSouthAmerica* | 1654849 | 1 | 39.0 | 1557 |
| 867 | NAM128 | G_Mx_2005_337 | Mexico | MexicoCity | *hspSWEuropeMexico* | *hspSWEuropeNorthAmerica* | 1697806 | 1 | 38.9 | 1649 |
| 868 | NAM129 | G_Mx_2006_152 | Mexico | MexicoCity | *hspSWEuropeMexico* | *hspSWEuropeSouthAmerica* | 1613306 | 1 | 39.1 | 1512 |
| 869 | NAM130 | G_Mx_2006_46 | Mexico | MexicoCity | *hspSWEuropeHonduras* | *hspSWEuropeNorthAmerica* | 1636468 | 2 | 37.9 | 1634 |
| 870 | NAM131 | G_Mx_2006_583 | Mexico | MexicoCity | *hspSWEuropeMexico* | *hspSWEurope* | 1645512 | 1 | 39.1 | 1669 |
| 871 | NAM132 | C_Mx_2008_31 | Mexico | MexicoCity | *hspSWEuropeHonduras* | *hspSWEuropeSouthAmerica* | 1644819 | 2 | 38.0 | 1559 |
| 872 | NAM133 | C_Mx_2010_5 | Mexico | MexicoCity | *hspSWEuropeMexico* | *hspSWEuropeSouthAmerica* | 1645840 | 1 | 39.1 | 1541 |
| 873 | NAM134 | C_Mx_2010_8 | Mexico | MexicoCity | *hspSWEuropeHonduras* | *hspSWEuropeSouthAmerica* | 1706946 | 4 | 37.1 | 1675 |
| 874 | NAM135 | C_Mx_2011_69 | Mexico | MexicoCity | *hspSWEuropeMexico* | *hspSWEurope* | 1693680 | 2 | 34.5 | 1618 |
| 875 | NAM136 | G_Mx_2011_124 | Mexico | MexicoCity | *hspSWEuropeMexico* | *hspSWEuropeSouthAmerica* | 1661266 | 1 | 39.0 | 1597 |
| 876 | NAM137 | G_Mx_2011_147 | Mexico | MexicoCity | *hspSWEuropeMexico* | *hspSWEuropeSouthAmerica* | 1667329 | 1 | 38.9 | 1569 |
| 877 | NAM138 | C_Mx_2011_152 | Mexico | MexicoCity | *hspSWEuropeHonduras* | *hspSWEuropeSouthAmerica* | 1638427 | 2 | 38.1 | 1562 |
| 878 | NAM139 | C_Mx_2011_171 | Mexico | MexicoCity | *hspAfrica1MiscAmerica* | *hspSWEuropeNorthAmerica* | 1645502 | 1 | 39.1 | 1553 |
| 879 | NAM140 | G_Mx_2003_356 | Mexico | MexicoCity | *hspAfrica1MiscAmerica* | *hspSWEuropeNorthAmerica* | 1676345 | 38 | 38.7 | 1531 |
| 880 | NAM141 | G_Mx_2005_100 | Mexico | MexicoCity | *hspSWEuropeMexico* | *hspSWEuropeSouthAmerica* | 1696927 | 44 | 38.0 | 1542 |
| 881 | NAM142 | G_Mx_2005_104 | Mexico | MexicoCity | *hspAfrica1MiscAmerica* | *hspSWEuropeNorthAmerica* | 1682427 | 30 | 38.0 | 1540 |
| 882 | NAM143 | G_Mx_2010_64 | Mexico | MexicoCity | *hspAfrica1MiscAmerica* | *hspSWEuropeNorthAmerica* | 1703200 | 38 | 38.7 | 1562 |
| 883 | NAM144 | G_Mx_2005_109 | Mexico | MexicoCity | *hspSWEuropeMexico* | *hspSWEurope* | 1612662 | 63 | 37.8 | 1508 |
| 884 | NAM145 | C_Mx_2005_33 | Mexico | MexicoCity | *hspSWEuropeMexico* | *hspSWEuropeNorthAmerica* | 1730331 | 59 | 37.8 | 1606 |
| 885 | NAM146 | G_Mx_2005_70 | Mexico | MexicoCity | *hspSWEuropeMexico* | *hspSWEuropeSouthAmerica* | 1688787 | 31 | 38.1 | 1546 |
| 886 | NAM147 | G_Mx_2006_53 | Mexico | MexicoCity | *hspSWEuropeHonduras* | *hspSWEurope* | 1629399 | 22 | 38.4 | 1514 |
| 887 | NAM148 | C_Mx_2006_52 | Mexico | MexicoCity | *hspSWEuropeMexico* | *hspSWEuropeSouthAmerica* | 1669032 | 34 | 38.9 | 1546 |
| 888 | NAM149 | C_Mx_2006_577 | Mexico | MexicoCity | *hspSWEuropeMexico* | *hspSWEuropeSouthAmerica* | 1681991 | 26 | 38.6 | 1545 |
| 889 | NAM150 | C_Mx_2010_10 | Mexico | MexicoCity | *hspSWEuropeHonduras* | *hspSWEuropeSouthAmerica* | 1674444 | 76 | 37.8 | 1543 |
| 890 | NAM151 | C_Mx_2010_103 | Mexico | MexicoCity | *hspSWEuropeHonduras* | *hspSWEuropeSouthAmerica* | 1650514 | 37 | 38.1 | 1528 |
| 891 | NAM152 | C_Mx_2010_2 | Mexico | MexicoCity | *hspSWEuropeMexico* | *hspSWEuropeSouthAmerica* | 1661203 | 30 | 38.8 | 1529 |
| 892 | NAM153 | G_Mx_2011_131 | Mexico | MexicoCity | *hspAfrica1MiscAmerica* | *hspSWEuropeNorthAmerica* | 1703001 | 38 | 39.1 | 1566 |
| 893 | NAM154 | G_Mx_2011_41 | Mexico | MexicoCity | *hspSWEuropeMexico* | *hspSWEuropeSouthAmerica* | 1699853 | 40 | 38.8 | 1568 |
| 894 | NAM155 | G_Mx_2003_136 | Mexico | MexicoCity | *hspSWEuropeMexico* | *hspSWEurope* | 1657128 | 112 | 36.7 | 1522 |
| 895 | NAM156 | G_Mx_2005_107 | Mexico | MexicoCity | *hspSWEuropeHonduras* | *hspSWEuropeSouthAmerica* | 1716270 | 35 | 38.4 | 1576 |
| 896 | NAM157 | C_Mx_2006_356 | Mexico | MexicoCity | *hspAfrica1MiscAmerica* | *hspSWEuropeNorthAmerica* | 1690901 | 25 | 38.7 | 1540 |
| 897 | NAM158 | C_Mx_2006_4 | Mexico | MexicoCity | *hspSWEuropeMexico* | *hspSWEuropeHonduras* | 1677211 | 28 | 38.7 | 1543 |
| 898 | NAM159 | G_Mx_2006_513 | Mexico | MexicoCity | *hspAfrica1MiscAmerica* | *hspSWEuropeNorthAmerica* | 1768260 | 55 | 38.0 | 1646 |
| 899 | NAM160 | C_Mx_2006_664 | Mexico | MexicoCity | *hspAfrica1MiscAmerica* | *hspSWEuropeNorthAmerica* | 1686099 | 50 | 38.2 | 1551 |
| 900 | NAM161 | C_Mx_2006_677 | Mexico | MexicoCity | *hspSWEuropeMexico* | *hspSWEuropeSouthAmerica* | 1629650 | 37 | 38.6 | 1502 |
| 901 | NAM162 | C_Mx_2010_11 | Mexico | MexicoCity | *hspSWEuropeHonduras* | *hspSWEuropeSouthAmerica* | 1657306 | 77 | 37.7 | 1529 |
| 902 | NAM163 | G_Mx_2010_13 | Mexico | MexicoCity | *hspSWEuropeMexico* | *hspSWEurope* | 1708271 | 49 | 38.2 | 1560 |
| 903 | NAM164 | 2010-67 | Mexico | MexicoCity | *Unknown* | *hspSWEuropeNorthAmerica* | 1661412 | 136 | 37.7 | 1544 |
| 904 | NAM165 | G_Mx_2010_99 | Mexico | MexicoCity | *hspSWEuropeMexico* | *hspSWEuropeSouthAmerica* | 1605913 | 50 | 39.0 | 1492 |
| 905 | NAM166 | C_Mx_2010_100 | Mexico | MexicoCity | *hspSWEuropeMexico* | *hspSWEuropeSouthAmerica* | 1655754 | 49 | 38.8 | 1521 |
| 906 | NAM167 | M_Mx_2011_113 | Mexico | MexicoCity | *hspSWEuropeMexico* | *hspSWEurope* | 1598782 | 55 | 38.6 | 1486 |
| 907 | NAM168 | C_Mx_2011_117 | Mexico | MexicoCity | *hspSWEuropeMexico* | *hspSWEurope* | 1599458 | 37 | 38.6 | 1496 |
| 908 | NAM169 | C_Mx_2011_136 | Mexico | MexicoCity | *hspSWEuropeMexico* | *hspSWEuropeSouthAmerica* | 1641622 | 81 | 38.0 | 1525 |
| 909 | NAM170 | M_Mx_2003_230 | Mexico | MexicoCity | *hspSWEuropeHonduras* | *hspSWEuropeNorthAmerica* | 1642507 | 60 | 38.2 | 1518 |
| 910 | NAM171 | M_Mx_2005_106 | Mexico | MexicoCity | *hspSWEuropeHonduras* | *hspSWEuropeSouthAmerica* | 1607854 | 31 | 38.7 | 1488 |
| 911 | NAM172 | M_Mx_2005_115 | Mexico | MexicoCity | *hspSWEuropeMexico* | *hspSWEuropeSouthAmerica* | 1664338 | 86 | 38.4 | 1544 |
| 912 | NAM173 | M_Mx_2005_119 | Mexico | MexicoCity | *hspSWEuropeMexico* | *hspSWEuropeSouthAmerica* | 1645091 | 28 | 38.4 | 1509 |
| 913 | NAM174 | M_Mx_2005_152 | Mexico | MexicoCity | *hspSWEuropeMexico* | *hspSWEuropeSouthAmerica* | 1632933 | 33 | 38.5 | 1500 |
| 914 | NAM175 | M_Mx_2006_196 | Mexico | MexicoCity | *hspSWEuropeMexico* | *hspSWEuropeSouthAmerica* | 1647922 | 77 | 37.6 | 1519 |
| 915 | NAM176 | M_Mx_2006_276 | Mexico | MexicoCity | *hspAfrica1MiscAmerica* | *hspSWEuropeSouthAmerica* | 1707900 | 83 | 38.3 | 1585 |
| 916 | NAM177 | M_Mx_2006_427 | Mexico | MexicoCity | *hspSWEuropeHonduras* | *hspSWEuropeSouthAmerica* | 1651780 | 64 | 38.2 | 1523 |
| 917 | NAM178 | M_Mx_2006_416 | Mexico | MexicoCity | *hspAfrica1MiscAmerica* | *hspSWEuropeNorthAmerica* | 1732546 | 55 | 37.9 | 1598 |
| 918 | NAM179 | M_Mx_2006_690 | Mexico | MexicoCity | *hspSWEuropeHonduras* | *hspSWEuropeSouthAmerica* | 1649151 | 70 | 38.0 | 1528 |
| 919 | NAM180 | M_Mx_2008_34 | Mexico | MexicoCity | *hspSWEuropeMexico* | *hspSWEuropeSouthAmerica* | 1681292 | 44 | 37.8 | 1554 |
| 920 | NAM182 | 2010-82 | Mexico | Unknown | *Unknown* | *hspSWEuropeNorthAmerica* | 1667669 | 1 | 39.0 | 1589 |
| 921 | NAM184 | 2011-119 | Mexico | Unknown | *Unknown* | *hspSWEurope* | 1597743 | 1 | 39.1 | 1572 |
| 922 | NAM186 | 2011-80 | Mexico | Unknown | *Unknown* | *hspSWEuropeSouthAmerica* | 1641692 | 1 | 38.9 | 1625 |
| 923 | NAM191 | 2010-66 | Mexico | Unknown | *Unknown* | *hspSWEuropeSouthAmerica* | 1640634 | 1 | 38.9 | 1619 |
| 924 | NAM192 | 2011-20 | Mexico | Unknown | *Unknown* | *hspSWEurope* | 1640285 | 1 | 38.9 | 1540 |
| 925 | NAM193 | MCms1054 | Mexico | MexicoCity | *hspAfrica1MiscAmerica* | *hspSWEuropeNorthAmerica* | 1656578 | 117 | 38.0 | 1516 |
| 926 | NAM194 | MCms1055 | Mexico | MexicoCity | *hspSWEuropeMexico* | *hspSWEuropeSouthAmerica* | 1676870 | 63 | 39.0 | 1549 |
| 927 | NAM195 | MCms1063 | Mexico | MexicoCity | *hspAfrica1MiscAmerica* | *hspSWEuropeNorthAmerica* | 1692058 | 72 | 38.7 | 1547 |
| 928 | NAM196 | MCms1078 | Mexico | MexicoCity | *hspSWEuropeMexico* | *hspSWEuropeSouthAmerica* | 1649218 | 81 | 38.4 | 1518 |
| 929 | NAM197 | MCms1080 | Mexico | MexicoCity | *hspSEurope* | *hspSWEuropeSouthAmerica* | 1661740 | 91 | 38.5 | 1510 |
| 930 | NAM198 | MCms931 | Mexico | MexicoCity | *hspSEurope* | *hspSWEuropeSouthAmerica* | 1668681 | 96 | 38.8 | 1525 |
| 931 | NAM199 | MG_2003_107 | Mexico | MexicoCity | *hspAfrica1MiscAmerica* | *hspSWEuropeNorthAmerica* | 1757909 | 145 | 37.4 | 1619 |
| 932 | NAM200 | MG_2003_98 | Mexico | MexicoCity | *hspAfrica1MiscAmerica* | *hspSWEuropeNorthAmerica* | 1715886 | 85 | 38.3 | 1591 |
| 933 | NAM201 | MG_2005_98 | Mexico | MexicoCity | *hspSWEuropeMexico* | *hspSWEuropeSouthAmerica* | 1653883 | 82 | 38.4 | 1528 |
| 934 | NAM202 | MG_2006_407 | Mexico | MexicoCity | *hspSWEuropeMexico* | *hspSWEuropeSouthAmerica* | 1746012 | 89 | 38.3 | 1635 |
| 935 | NAM203 | MG_2006_479 | Mexico | MexicoCity | *hspSWEuropeHonduras* | *hspSWEuropeSouthAmerica* | 1607232 | 43 | 38.8 | 1493 |
| 936 | NAM204 | MGms13 | Mexico | MexicoCity | *hspSEurope* | *hspSWEuropeSouthAmerica* | 1653926 | 111 | 37.9 | 1516 |
| 937 | NAM205 | MGms15 | Mexico | MexicoCity | *hspSEurope* | *hspSWEuropeSouthAmerica* | 1692789 | 88 | 38.5 | 1576 |
| 938 | NAM206 | MGms167 | Mexico | MexicoCity | *hspAfrica1MiscAmerica* | *hspSWEuropeNorthAmerica* | 1680682 | 73 | 38.8 | 1521 |
| 939 | NAM207 | MGms176 | Mexico | MexicoCity | *hspSEurope* | *hspSWEuropeSouthAmerica* | 1672233 | 225 | 40.1 | 1519 |
| 940 | NAM208 | MGms2 | Mexico | MexicoCity | *hspSWEuropeMexico* | *hspSWEuropeSouthAmerica* | 1712465 | 102 | 38.1 | 1581 |
| 941 | NAM209 | MGms203 | Mexico | MexicoCity | *hspSWEuropeMexico* | *hspSWEuropeSouthAmerica* | 1732925 | 84 | 38.2 | 1600 |
| 942 | NAM210 | MGms23 | Mexico | MexicoCity | *hspSEurope* | *hspSWEuropeSouthAmerica* | 1664931 | 89 | 37.7 | 1547 |
| 943 | NAM211 | MGms44 | Mexico | MexicoCity | *hspAfrica1MiscAmerica* | *hspSWEuropeNorthAmerica* | 1635449 | 113 | 37.8 | 1496 |
| 944 | NAM212 | 2003_103 | Mexico | MexicoCity | *hspSWEuropeMexico* | *hspSWEuropeNorthAmerica* | 1673433 | 77 | 37.4 | 1578 |
| 945 | NAM213 | 2004_20 | Mexico | MexicoCity | *hspSWEuropeMexico* | *hspSWEuropeSouthAmerica* | 1628314 | 37 | 38.8 | 1521 |
| 946 | NAM214 | 2005_126 | Mexico | MexicoCity | *hspSWEuropeMexico* | *hspSWEuropeSouthAmerica* | 1628448 | 47 | 37.4 | 1504 |
| 947 | NAM215 | 2005_72 | Mexico | MexicoCity | *hspAfrica1MiscAmerica* | *hspSWEuropeNorthAmerica* | 1659834 | 56 | 37.5 | 1535 |
| 948 | NAM216 | 2006_103 | Mexico | MexicoCity | *hspAfrica1MiscAmerica* | *hspSWEuropeNorthAmerica* | 1674943 | 47 | 37.9 | 1561 |
| 949 | NAM217 | 2006_480 | Mexico | MexicoCity | *hspSWEuropeMexico* | *hspSWEuropeSouthAmerica* | 1624363 | 42 | 39.4 | 1502 |
| 950 | NAM218 | 2006_56 | Mexico | MexicoCity | *hspSWEuropeHonduras* | *hspSWEuropeSouthAmerica* | 1642196 | 47 | 38.6 | 1540 |
| 951 | NAM219 | 2012_26 | Mexico | MexicoCity | *hspSWEuropeMexico* | *hspSWEuropeSouthAmerica* | 1630210 | 34 | 38.7 | 1532 |
| 952 | NAM220 | 2003_84 | Mexico | MexicoCity | *hspSWEuropeMexico* | *hspSWEuropeSouthAmerica* | 1638214 | 53 | 38.1 | 1522 |
| 953 | NAM221 | 2004_02 | Mexico | MexicoCity | *hspSWEuropeMexico* | *hspSWEuropeSouthAmerica* | 1693055 | 77 | 37.6 | 1601 |
| 954 | NAM222 | 2017-20 | Mexico | Unknown | *Unknown* | *hspSWEuropeSouthAmerica* | 1667960 | 63 | 38.7 | 1550 |
| 955 | NAM223 | 2017-27 | Mexico | Unknown | *Unknown* | *hspSWEuropeSouthAmerica* | 1667459 | 68 | 38.0 | 1549 |
| 956 | NAM224 | 2017-38 | Mexico | Unknown | *Unknown* | *hspSWEuropeSouthAmerica* | 1640186 | 67 | 39.8 | 1510 |
| 957 | NAM225 | 2017-44 | Mexico | Unknown | *Unknown* | *hspSWEuropeNorthAmerica* | 1613848 | 61 | 37.6 | 1516 |
| 958 | NAM226 | 2017-54 | Mexico | Unknown | *Unknown* | *hspSWEuropeNorthAmerica* | 1621684 | 51 | 38.9 | 1532 |
| 959 | NAM227 | 2017-55 | Mexico | Unknown | *Unknown* | *hspSWEuropeNorthAmerica* | 1621385 | 51 | 39.1 | 1535 |
| 960 | NAM228 | 2017-58 | Mexico | Unknown | *Unknown* | *hspSWEuropeNorthAmerica* | 1672132 | 56 | 37.7 | 1546 |
| 961 | NAM229 | 2017-69 | Mexico | Unknown | *Unknown* | *hspSWEurope* | 1635024 | 41 | 37.7 | 1525 |
| 962 | NAM230 | 2017-78 | Mexico | Unknown | *Unknown* | *hspSWEurope* | 1651292 | 48 | 37.9 | 1548 |
| 963 | NAM231 | 2017-79 | Mexico | Unknown | *Unknown* | *hspSWEuropeNorthAmerica* | 1647657 | 54 | 37.6 | 1539 |
| 964 | NAM232 | 2017-94 | Mexico | Unknown | *Unknown* | *hspSWEuropeSouthAmerica* | 1658511 | 55 | 37.8 | 1526 |
| 965 | NAM233 | 2017-105 | Mexico | Unknown | *Unknown* | *hspSWEuropeSouthAmerica* | 1631163 | 53 | 38.6 | 1519 |
| 966 | NAM234 | 2017-107 | Mexico | Unknown | *Unknown* | *hspSWEuropeNorthAmerica* | 1627684 | 56 | 38.6 | 1536 |
| 967 | NAM235 | 2017-124 | Mexico | Unknown | *Unknown* | *hspSWEuropeSouthAmerica* | 1683835 | 68 | 38.2 | 1563 |
| 968 | NAM236 | 2017-143 | Mexico | Unknown | *Unknown* | *hspSWEuropeNorthAmerica* | 1672047 | 50 | 37.6 | 1548 |
| 969 | NAM237 | 2017-149 | Mexico | Unknown | *Unknown* | *hspSWEuropeSouthAmerica* | 1672961 | 98 | 36.8 | 1564 |
| 970 | NAM238 | 2017-151 | Mexico | Unknown | *Unknown* | *hspSWEuropeSouthAmerica* | 1671271 | 88 | 36.8 | 1563 |
| 971 | NAM239 | 2017-161 | Mexico | Unknown | *Unknown* | *hspSWEuropeNorthAmerica* | 1672121 | 63 | 37.8 | 1547 |
| 972 | NAM240 | 2017-177 | Mexico | Unknown | *Unknown* | *hspSWEuropeNorthAmerica* | 1647686 | 54 | 37.5 | 1539 |
| 973 | NAM241 | 2017-192 | Mexico | Unknown | *Unknown* | *hspSWEuropeNorthAmerica* | 1652213 | 67 | 39.5 | 1519 |
| 974 | NAM242 | 2017-198 | Mexico | Unknown | *Unknown* | *hspSWEuropeNorthAmerica* | 1616619 | 57 | 37.8 | 1519 |
| 975 | NAM243 | 2017-199 | Mexico | Unknown | *Unknown* | *hspSWEuropeNorthAmerica* | 1672216 | 52 | 37.6 | 1548 |
| 976 | NAM244 | 2018-03 | Mexico | Unknown | *Unknown* | *hspSWEuropeSouthAmerica* | 1648345 | 53 | 37.7 | 1533 |
| 977 | NAM245 | 2018-04 | Mexico | Unknown | *Unknown* | *hspSWEuropeSouthAmerica* | 1647880 | 45 | 37.4 | 1533 |
| 978 | NAM246 | 2018-05 | Mexico | Unknown | *Unknown* | *hspSWEuropeSouthAmerica* | 1648210 | 45 | 37.3 | 1533 |
| 979 | NAM247 | 2018-06 | Mexico | Unknown | *Unknown* | *hspSWEuropeSouthAmerica* | 1656719 | 56 | 39.1 | 1520 |
| 980 | NAM248 | 2019-1 | Mexico | Unknown | *Unknown* | *hspSWEuropeSouthAmerica* | 1656568 | 39 | 38.5 | 1536 |
| 981 | NAM249 | 2019-03 | Mexico | Unknown | *Unknown* | *hspSWEuropeSouthAmerica* | 1641332 | 83 | 38.2 | 1521 |
| 982 | NAM250 | 2019-05 | Mexico | Unknown | *Unknown* | *hspAfrica1SAfrica* | 1664731 | 45 | 38.1 | 1572 |
| 983 | NAM251 | 2019-06 | Mexico | Unknown | *Unknown* | *hspSWEuropeSouthAmerica* | 1696123 | 59 | 36.5 | 1565 |
| 984 | NAM252 | 2019-08 | Mexico | Unknown | *Unknown* | *hspSWEuropeSouthAmerica* | 1656630 | 37 | 38.7 | 1536 |
| 985 | NAM253 | 2004_137 | Mexico | Chihuahua | *hspAfrica1MiscAmerica* | *hspSWEuropeNorthAmerica* | 1642388 | 44 | 37.9 | 1534 |
| 986 | NAM254 | 2003_279 | Mexico | MilpaAlta | *hspSWEuropeMexico* | *hspSWEurope* | 1645938 | 36 | 38.1 | 1544 |
| 987 | NAM255 | 2003_292 | Mexico | MilpaAlta | *hspSWEuropeMexico* | *hspSWEuropeSouthAmerica* | 1644270 | 38 | 38.1 | 1541 |
| 988 | NAM256 | 2004_102 | Mexico | Chihuahua | *hspAfrica1MiscAmerica* | *hspSWEuropeNorthAmerica* | 1657958 | 37 | 38.0 | 1525 |
| 989 | NAM257 | 2004_125 | Mexico | Chihuahua | *hspSAmerind* | *hspAmerind* | 1571156 | 15 | 39.3 | 1487 |
| 990 | NAM258 | 2004_37 | Mexico | MilpaAlta | *hspSWEuropeHonduras* | *hspSWEuropeSouthAmerica* | 1651609 | 30 | 38.7 | 1542 |
| 991 | NAM259 | 2004_67 | Mexico | MilpaAlta | *hspSWEurope* | *hspSWEuropeColombia* | 1616544 | 35 | 37.3 | 1515 |
| 992 | NAM260 | 2004_40 | Mexico | MilpaAlta | *hspSWEurope* | *hspSWEurope* | 1586808 | 34 | 37.9 | 1486 |
| 993 | NAM261 | 2004_51 | Mexico | MilpaAlta | *hspAfrica1NAmerica* | *hspAfrica1NAmerica* | 1639801 | 26 | 38.8 | 1516 |
| 994 | NAM262 | 2004_210 | Mexico | MilpaAlta | *hspSWEurope* | *hspSWEurope* | 1636979 | 55 | 36.4 | 1543 |
| 995 | NAM263 | 2004_42 | Mexico | MilpaAlta | *hspSWEurope* | *hspSWEurope* | 1618576 | 39 | 38.1 | 1500 |
| 996 | NAM264 | 2003_368 | Mexico | Nayarit | *hspAfrica1MiscAmerica* | *hspSWEuropeNorthAmerica* | 1624523 | 38 | 38.6 | 1517 |
| 997 | NAM265 | 2004_84 | Mexico | Chihuahua | *hspSWEuropeHonduras* | *hspSWEuropeHonduras* | 1651980 | 33 | 37.7 | 1547 |
| 998 | NAM266 | 2002_14 | Mexico | MilpaAlta | *hspSAmerind* | *hspAmerind* | 1599680 | 23 | 38.0 | 1514 |
| 999 | NAM267 | Mx_04_130 | Mexico | Chihuahua | *hspSWEuropeMexico* | *hspSWEuropeSouthAmerica* | 1683669 | 53 | 36.0 | 1571 |
| 1000 | NAM268 | MM2006_106 | Mexico | MexicoCity | *hspAfrica1MiscAmerica* | *hspSWEuropeNorthAmerica* | 1768793 | 61 | 38.6 | 1633 |
| 1001 | NAM270 | MG2006_449 | Mexico | MexicoCity | *hspSWEuropeHonduras* | *hspSWEuropeSouthAmerica* | 1607095 | 79 | 38.6 | 1485 |
| 1002 | NAM271 | MEX-028ta | Mexico | Unknown | *Unknown* | *hspAmerind* | 1562280 | 1 | 39.0 | 1535 |
| 1003 | NAM272 | CAN_005 | Canada | NorthwestTerritories | *hspNAmerind* | *hspAmerind* | 1521597 | 50 | 37.6 | 1421 |
| 1004 | NAM273 | CAN_010 | Canada | NorthwestTerritories | *hspNAmerind* | *hspAmerind* | 1506179 | 88 | 38.4 | 1406 |
| 1005 | NAM274 | CAN_011 | Canada | NorthwestTerritories | *hspNAmerind* | *hspAmerind* | 1542813 | 85 | 36.9 | 1439 |
| 1006 | NAM275 | CAN_019 | Canada | NorthwestTerritories | *hspNAmerind* | *hspAmerind* | 1505987 | 46 | 37.6 | 1412 |
| 1007 | NAM276 | CAN_095 | Canada | NorthwestTerritories | *hspNAmerind* | *hspAmerind* | 1522475 | 56 | 37.6 | 1430 |
| 1008 | NAM277 | CAN_136 | Canada | NorthwestTerritories | *hspNAmerind* | *hspAmerind* | 1518472 | 50 | 37.5 | 1426 |
| 1009 | NAM278 | CAN_172 | Canada | NorthwestTerritories | *hspNAmerind* | *hspAmerind* | 1524747 | 73 | 37.1 | 1433 |
| 1010 | NAM279 | CAN_247 | Canada | NorthwestTerritories | *hspNAmerind* | *hspAmerind* | 1507187 | 41 | 38.6 | 1417 |
| 1011 | NAM280 | CAN_258 | Canada | NorthwestTerritories | *hspNAmerind* | *hspAmerind* | 1515148 | 36 | 38.4 | 1415 |
| 1012 | NAM281 | CAN_290 | Canada | NorthwestTerritories | *hspNAmerind* | *hspAmerind* | 1501432 | 41 | 37.6 | 1414 |
| 1013 | OCN001 | ausabrJ05 | Australia | Jigalong | *hspSAmerind* | *hpSahul* | 1518447 | 60 | 38.0 | 1405 |
| 1014 | OCN002 | BM012A | Australia | Perth | *hspNEurope* | *hspNEurope* | 1660425 | 1 | 38.9 | 1573 |
| 1015 | OCN003 | BM013A | Australia | Perth | *hspNEurope* | *hspSEurope* | 1604233 | 1 | 39.0 | 1503 |
| 1016 | OCN004 | Sahul64 | Australia | WesternAustralia | *hspNEurope* | *hpSahul* | 1644275 | 54 | 38.0 | 1546 |
| 1017 | OCN005 | PNG84A | PapuaNewGuinea | Goroka | *hspSAmerind* | *hpSahul* | 1531450 | 1 | 38.8 | 1487 |
| 1018 | OCN006 | Neuguinea102 | PapuaNewGuinea | Goroka | *hpSahul* | *hpSahul* | 1538571 | 45 | 38.7 | 1478 |
| 1019 | OCN007 | Neuguinea12 | PapuaNewGuinea | Goroka | *hpSahul* | *hpSahul* | 1542652 | 56 | 39.5 | 1473 |
| 1020 | OCN008 | Neuguinea13 | PapuaNewGuinea | Goroka | *hpSahul* | *hpSahul* | 1540070 | 43 | 38.5 | 1465 |
| 1021 | OCN009 | Neuguinea22 | PapuaNewGuinea | Goroka | *hpSahul* | *hpSahul* | 1542124 | 39 | 39.0 | 1468 |
| 1022 | OCN010 | Neuguinea25 | PapuaNewGuinea | Goroka | *hpSahul* | *hpSahul* | 1541461 | 44 | 38.7 | 1469 |
| 1023 | OCN011 | Neuguinea38 | PapuaNewGuinea | Goroka | *hpSahul* | *hpSahul* | 1534670 | 55 | 38.7 | 1467 |
| 1024 | OCN012 | Neuguinea42 | PapuaNewGuinea | Goroka | *hpSahul* | *hpSahul* | 1540553 | 41 | 38.3 | 1473 |
| 1025 | OCN013 | Neuguinea46 | PapuaNewGuinea | Goroka | *hpSahul* | *hpSahul* | 1517898 | 41 | 38.4 | 1456 |
| 1026 | OCN014 | Neuguinea51 | PapuaNewGuinea | Goroka | *hpSahul* | *hpSahul* | 1543154 | 51 | 39.1 | 1471 |
| 1027 | OCN015 | Neuguinea56 | PapuaNewGuinea | Goroka | *hpSahul* | *hpSahul* | 1541226 | 51 | 38.4 | 1466 |
| 1028 | OCN016 | Neuguinea63 | PapuaNewGuinea | Goroka | *hpSahul* | *hpSahul* | 1545399 | 37 | 39.5 | 1476 |
| 1029 | OCN017 | Neuguinea66 | PapuaNewGuinea | Goroka | *hpSahul* | *hpSahul* | 1540624 | 52 | 38.9 | 1480 |
| 1030 | OCN018 | Neuguinea70 | PapuaNewGuinea | Goroka | *hpSahul* | *hpSahul* | 1540735 | 42 | 38.6 | 1469 |
| 1031 | OCN019 | Neuguinea78 | PapuaNewGuinea | Goroka | *hpSahul* | *hpSahul* | 1543458 | 46 | 38.8 | 1478 |
| 1032 | OCN020 | Neuguinea87 | PapuaNewGuinea | Goroka | *hpSahul* | *hpSahul* | 1542452 | 38 | 39.3 | 1479 |
| 1033 | OCN021 | Neuguinea97 | PapuaNewGuinea | Goroka | *hpSahul* | *hpSahul* | 1547595 | 47 | 38.9 | 1478 |
| 1034 | OCN022 | HP97011 | Australia | Perth | *Unknown* | *hspSEurope* | 1552062 | 23 | 39.2 | 1480 |
| 1035 | OCN023 | HP98123 | Australia | Perth | *Unknown* | *hspSWEuropeAustralia* | 1634330 | 42 | 38.4 | 1531 |
| 1036 | OCN024 | HP98317 | Australia | Perth | *Unknown* | *hspSWEuropeAustralia* | 1602375 | 36 | 38.8 | 1495 |
| 1037 | OCN025 | HP98490 | Australia | Perth | *Unknown* | *hspSWEuropeAustralia* | 1687678 | 38 | 38.8 | 1590 |
| 1038 | OCN026 | HP99216 | Australia | Perth | *Unknown* | *hspAfrica1SAfrica* | 1628204 | 46 | 38.6 | 1523 |
| 1039 | OCN027 | HP99244 | Australia | Perth | *Unknown* | *hpAsia2* | 1612906 | 31 | 39.5 | 1525 |
| 1040 | OCN028 | HP99255 | Australia | Perth | *Unknown* | *hspSWEuropeAustralia* | 1564730 | 28 | 38.4 | 1448 |
| 1041 | OCN029 | HP99316 | Australia | Perth | *Unknown* | *hpAsia2* | 1660794 | 46 | 37.8 | 1562 |
| 1042 | OCN030 | HP99330 | Australia | Perth | *Unknown* | *hspAfrica1SAfrica* | 1596254 | 38 | 39.0 | 1493 |
| 1043 | OCN031 | HP99392 | Australia | Perth | *Unknown* | *hspSEurope* | 1631211 | 31 | 38.6 | 1544 |
| 1044 | OCN032 | HP99647 | Australia | Perth | *Unknown* | *hspSEurope* | 1600967 | 30 | 39.3 | 1476 |
| 1045 | OCN033 | HP99648 | Australia | Perth | *Unknown* | *hspAfrica1SAfrica* | 1622797 | 29 | 38.9 | 1519 |
| 1046 | OCN034 | HP99689 | Australia | Perth | *Unknown* | *hspSEurope* | 1615212 | 47 | 38.0 | 1507 |
| 1047 | OCN035 | HPAS14 | Australia | Perth | *Unknown* | *hspNEurope* | 1577919 | 36 | 39.0 | 1480 |
| 1048 | OCN036 | HPJ013 | Australia | Perth | *Unknown* | *hspNEurope* | 1662228 | 51 | 37.3 | 1547 |
| 1049 | OCN037 | HPJ024 | Australia | Perth | *Unknown* | *hpSahul* | 1684079 | 47 | 38.0 | 1594 |
| 1050 | OCN038 | HPJ025 | Australia | Perth | *Unknown* | *hpSahul* | 1682100 | 53 | 38.1 | 1583 |
| 1051 | OCN039 | HPJ040 | Australia | Perth | *Unknown* | *hspNEuropeAustralia* | 1647118 | 47 | 38.2 | 1530 |
| 1052 | OCN040 | HPJ050 | Australia | Perth | *Unknown* | *hpSahul* | 1682586 | 68 | 37.6 | 1580 |
| 1053 | OCN041 | HPJ099 | Australia | Perth | *Unknown* | *hspNEurope* | 1580798 | 28 | 38.0 | 1497 |
| 1054 | OCN042 | HPJ118 | Australia | Perth | *Unknown* | *hspNEuropeAustralia* | 1646903 | 42 | 38.3 | 1531 |
| 1055 | OCN043 | HPJ165 | Australia | Perth | *Unknown* | *hspNEurope* | 1694310 | 55 | 38.7 | 1581 |
| 1056 | OCN044 | ausabrAS41 | Australia | AliceSprings | *hpSahul* | *hpSahul* | 1516230 | 85 | 38.8 | 1396 |
| 1057 | OCN045 | ausabrAS48 | Australia | AliceSprings | *hpSahul* | *hpSahul* | 1516878 | 55 | 38.7 | 1414 |
| 1058 | OCN046 | ausabrAS49 | Australia | AliceSprings | *hpSahul* | *hpSahul* | 1517034 | 61 | 39.0 | 1411 |
| 1059 | OCN047 | ausabrJ11 | Australia | Jigalong | *hpSahul* | *hpSahul* | 1524904 | 54 | 38.7 | 1423 |
| 1060 | OCN048 | ausabrJ15 | Australia | Jigalong | *hpSahul* | *hpSahul* | 1526044 | 65 | 40.0 | 1423 |
| 1061 | OCN049 | ausabrJ20 | Australia | Jigalong | *hpSahul* | *hpSahul* | 1516098 | 39 | 39.9 | 1409 |
| 1062 | OCN050 | ausabrJ27 | Australia | Jigalong | *hpSahul* | *hpSahul* | 1552731 | 89 | 38.7 | 1439 |
| 1063 | OCN051 | ausabrp103 | Australia | Perth | *hpSahul* | *hpSahul* | 1507878 | 41 | 39.1 | 1417 |
| 1064 | OCN052 | ausabrp107 | Australia | Perth | *hpSahul* | *hpSahul* | 1645914 | 59 | 38.3 | 1543 |
| 1065 | OCN053 | ausabrP52 | Australia | Perth | *hpSahul* | *hpSahul* | 1508894 | 47 | 38.3 | 1408 |
| 1066 | OCN054 | ausabrP54 | Australia | Perth | *hpSahul* | *hpSahul* | 1525015 | 39 | 39.1 | 1428 |
| 1067 | OCN055 | ausabrP55 | Australia | Perth | *hpSahul* | *hpSahul* | 1655236 | 58 | 38.5 | 1545 |
| 1068 | OCN056 | ausabrp71 | Australia | Perth | *hpSahul* | *hpSahul* | 1658636 | 78 | 38.5 | 1554 |
| 1069 | OCN057 | ausabrp73 | Australia | Perth | *hpSahul* | *hpSahul* | 1602374 | 46 | 38.3 | 1505 |
| 1070 | OCN058 | ausabrp78 | Australia | Perth | *hpSahul* | *hpSahul* | 1643371 | 57 | 37.7 | 1541 |
| 1071 | OCN059 | ausabrp94 | Australia | Perth | *hpSahul* | *hpSahul* | 1610838 | 63 | 38.1 | 1516 |
| 1072 | OCN060 | auseurP207 | Australia | Perth | *hpSahul* | *hpSahul* | 1497470 | 27 | 38.9 | 1402 |
| 1073 | OCN061 | HP98285 | Australia | Perth | *Unknown* | *hpSahul* | 1513636 | 31 | 38.8 | 1414 |
| 1074 | OCN062 | HP99440 | Australia | Perth | *Unknown* | *hpSahul* | 1501251 | 26 | 38.3 | 1405 |
| 1075 | OCN063 | HP99511 | Australia | Perth | *Unknown* | *hpSahul* | 1529545 | 47 | 38.6 | 1431 |
| 1076 | OCN064 | HP99521 | Australia | Perth | *Unknown* | *hspAfrica1SAfrica* | 1624208 | 43 | 38.3 | 1522 |
| 1077 | OCN065 | HPAS23 | Australia | Perth | *Unknown* | *hpSahul* | 1519650 | 51 | 38.9 | 1427 |
| 1078 | OCN066 | HPJ003 | Australia | Perth | *Unknown* | *hpSahul* | 1518613 | 28 | 38.8 | 1406 |
| 1079 | OCN067 | HPJ022 | Australia | Perth | *Unknown* | *hpSahul* | 1524198 | 41 | 38.4 | 1426 |
| 1080 | OCN068 | HPJ023 | Australia | Perth | *Unknown* | *hpSahul* | 1524698 | 47 | 38.5 | 1411 |
| 1081 | OCN069 | HPJ055 | Australia | Perth | *Unknown* | *hpSahul* | 1529291 | 33 | 38.5 | 1431 |
| 1082 | OCN070 | HPJ056 | Australia | Perth | *Unknown* | *hpSahul* | 1531925 | 52 | 39.2 | 1424 |
| 1083 | OCN071 | HPJ057 | Australia | Perth | *Unknown* | *hpSahul* | 1523590 | 61 | 39.4 | 1414 |
| 1084 | OCN072 | HPJ071 | Australia | Perth | *Unknown* | *hpSahul* | 1524084 | 32 | 38.2 | 1429 |
| 1085 | OCN073 | HPJ098 | Australia | Perth | *Unknown* | *hpSahul* | 1504590 | 20 | 38.4 | 1404 |
| 1086 | OCN074 | HPJ117 | Australia | Perth | *Unknown* | *hpSahul* | 1516933 | 28 | 38.7 | 1415 |
| 1087 | OCN075 | HPJ119 | Australia | Perth | *Unknown* | *hpSahul* | 1537474 | 61 | 39.8 | 1428 |
| 1088 | OCN076 | HPJ148 | Australia | Perth | *Unknown* | *hpSahul* | 1529255 | 31 | 38.3 | 1439 |
| 1089 | OCN077 | HPJ156 | Australia | Perth | *Unknown* | *hpSahul* | 1565979 | 40 | 37.8 | 1459 |
| 1090 | OCN078 | HPJ207 | Australia | Perth | *Unknown* | *hpSahul* | 1527734 | 51 | 39.4 | 1431 |
| 1091 | OCN079 | HP00152 | Australia | Perth | *Unknown* | *hspNEuropeAustralia* | 1669430 | 78 | 38.1 | 1592 |
| 1092 | OCN080 | HP00192 | Australia | Perth | *Unknown* | *hspNEuropeAustralia* | 1625458 | 49 | 39.0 | 1515 |
| 1093 | OCN081 | HP00248 | Australia | Perth | *Unknown* | *hspNEuropeAustralia* | 1683463 | 36 | 37.9 | 1562 |
| 1094 | OCN082 | HP00260 | Australia | Perth | *Unknown* | *hspSWEuropeNorthAmerica* | 1597197 | 37 | 39.1 | 1495 |
| 1095 | OCN083 | HP01102 | Australia | Perth | *Unknown* | *hspAfrica1NAmerica* | 1634075 | 33 | 38.6 | 1513 |
| 1096 | OCN084 | HP01140 | Australia | Perth | *Unknown* | *hpSahul* | 1661792 | 61 | 38.4 | 1557 |
| 1097 | OCN085 | HP01193 | Australia | Perth | *Unknown* | *hpSahul* | 1617039 | 39 | 38.0 | 1521 |
| 1098 | OCN086 | HP01234 | Australia | Perth | *Unknown* | *hspSWEuropeAustralia* | 1658871 | 45 | 37.5 | 1546 |
| 1099 | OCN087 | HP01306 | Australia | Perth | *Unknown* | *hspNEuropeAustralia* | 1638947 | 48 | 38.4 | 1549 |
| 1100 | OCN088 | HP01316 | Australia | Perth | *Unknown* | *hpSahul* | 1657825 | 45 | 37.7 | 1550 |
| 1101 | OCN089 | HP01324 | Australia | Perth | *Unknown* | *hpSahul* | 1514250 | 27 | 38.1 | 1412 |
| 1102 | OCN090 | HP01330 | Australia | Perth | *Unknown* | *hspNEurope* | 1685767 | 44 | 38.6 | 1580 |
| 1103 | OCN091 | HP02140 | Australia | Perth | *Unknown* | *hspNEuropeAustralia* | 1634066 | 58 | 38.3 | 1530 |
| 1104 | OCN092 | HP03054 | Australia | Perth | *Unknown* | *hspNEuropeAustralia* | 1649552 | 44 | 37.9 | 1523 |
| 1105 | OCN093 | HP03127 | Australia | Perth | *Unknown* | *hpSahul* | 1647686 | 33 | 38.3 | 1544 |
| 1106 | OCN094 | HP03218 | Australia | Perth | *Unknown* | *hspSEurope* | 1643985 | 34 | 38.3 | 1533 |
| 1107 | OCN095 | HP04041 | Australia | Perth | *Unknown* | *hspNEuropeAustralia* | 1674734 | 39 | 38.7 | 1556 |
| 1108 | OCN096 | HP04042 | Australia | Perth | *Unknown* | *hspNEuropeAustralia* | 1601009 | 27 | 39.0 | 1499 |
| 1109 | OCN097 | HP04057 | Australia | Perth | *Unknown* | *hspSEurope* | 1582137 | 34 | 38.3 | 1458 |
| 1110 | OCN098 | HP04086 | Australia | Perth | *Unknown* | *hspSEurope* | 1622949 | 24 | 39.7 | 1538 |
| 1111 | OCN099 | HP04087 | Australia | Perth | *Unknown* | *hspNEurope* | 1625952 | 39 | 39.0 | 1515 |
| 1112 | OCN100 | HP05044 | Australia | Perth | *Unknown* | *hspSEurope* | 1632655 | 43 | 38.4 | 1517 |
| 1113 | OCN101 | HP06038 | Australia | Perth | *Unknown* | *hspSEurope* | 1638771 | 36 | 38.3 | 1546 |
| 1114 | OCN102 | HP06045 | Australia | Perth | *Unknown* | *hspAfrica1SAfrica* | 1645923 | 29 | 38.4 | 1533 |
| 1115 | OCN103 | HP06058 | Australia | Perth | *Unknown* | *hspNEurope* | 1647037 | 46 | 38.5 | 1551 |
| 1116 | OCN104 | HP06059 | Australia | Perth | *Unknown* | *hspAfrica1SAfrica* | 1666618 | 35 | 39.1 | 1558 |
| 1117 | OCN105 | HP07019 | Australia | Perth | *Unknown* | *hspSEurope* | 1584978 | 27 | 37.9 | 1479 |
| 1118 | OCN106 | HP07036 | Australia | Perth | *Unknown* | *hspSWEuropeAustralia* | 1639698 | 34 | 38.3 | 1530 |
| 1119 | OCN107 | HP08031 | Australia | Perth | *Unknown* | *hspAfrica1SAfrica* | 1630358 | 44 | 38.9 | 1521 |
| 1120 | OCN108 | HP08058 | Australia | Perth | *Unknown* | *hspSWEuropeAustralia* | 1547231 | 28 | 39.1 | 1442 |
| 1121 | OCN109 | HP08061 | Australia | Perth | *Unknown* | *hspSWEuropeAustralia* | 1646844 | 43 | 37.7 | 1541 |
| 1122 | OCN110 | HP08072 | Australia | Perth | *Unknown* | *hspAfrica1NAmerica* | 1629992 | 30 | 38.8 | 1513 |
| 1123 | OCN111 | HP08073 | Australia | Perth | *Unknown* | *hspSEurope* | 1614312 | 35 | 38.7 | 1508 |
| 1124 | OCN112 | HP08074 | Australia | Perth | *Unknown* | *hspSWEuropeAustralia* | 1569108 | 29 | 39.1 | 1477 |
| 1125 | OCN113 | HP09046 | Australia | Perth | *Unknown* | *hspAfrica1SAfrica* | 1626738 | 47 | 38.6 | 1513 |
| 1126 | OCN114 | HP11004 | Australia | Perth | *Unknown* | *hspSEurope* | 1575902 | 32 | 38.1 | 1490 |
| 1127 | OCN115 | HP11005 | Australia | Perth | *Unknown* | *hspNEurope* | 1575376 | 34 | 39.0 | 1490 |
| 1128 | OCN116 | HP11011 | Australia | Perth | *Unknown* | *hspSEurope* | 1650911 | 42 | 38.7 | 1560 |
| 1129 | OCN117 | HP11013 | Australia | Perth | *Unknown* | *hspSEurope* | 1619004 | 35 | 39.1 | 1522 |
| 1130 | OCN118 | HP11020 | Australia | Perth | *Unknown* | *hspNEurope* | 1615516 | 44 | 38.2 | 1538 |
| 1131 | OCN119 | HP11032 | Australia | Perth | *Unknown* | *hspNEurope* | 1623497 | 36 | 37.8 | 1553 |
| 1132 | OCN120 | HP11037 | Australia | Perth | *Unknown* | *hspSEurope* | 1653242 | 31 | 37.2 | 1546 |
| 1133 | OCN121 | HP11042 | Australia | Perth | *Unknown* | *hspSWEuropeAustralia* | 1643471 | 29 | 39.2 | 1551 |
| 1134 | OCN122 | HP11043 | Australia | Perth | *Unknown* | *hspSEurope* | 1633250 | 38 | 38.1 | 1528 |
| 1135 | OCN123 | HP11049 | Australia | Perth | *Unknown* | *hspSWEuropeAustralia* | 1552557 | 50 | 38.3 | 1446 |
| 1136 | OCN124 | HP11054 | Australia | Perth | *Unknown* | *hspNEurope* | 1584658 | 30 | 39.1 | 1479 |
| 1137 | OCN125 | HP11055 | Australia | Perth | *Unknown* | *hspAmerind* | 1587241 | 29 | 39.1 | 1481 |
| 1138 | OCN126 | HP11059 | Australia | Perth | *Unknown* | *hspEastAsia* | 1638916 | 38 | 37.0 | 1549 |
| 1139 | OCN127 | HP12002 | Australia | Perth | *Unknown* | *hspSWEuropeAustralia* | 1580451 | 21 | 39.1 | 1499 |
| 1140 | OCN128 | HP12014 | Australia | Perth | *Unknown* | *hspSEurope* | 1564796 | 31 | 38.7 | 1485 |
| 1141 | OCN129 | HP12019 | Australia | Perth | *Unknown* | *hspSEurope* | 1618733 | 34 | 38.7 | 1522 |
| 1142 | OCN130 | HP12020 | Australia | Perth | *Unknown* | *hspAfrica1NAmerica* | 1598666 | 33 | 39.1 | 1500 |
| 1143 | OCN131 | HP12036 | Australia | Perth | *Unknown* | *hspEastAsia* | 1603777 | 32 | 37.5 | 1529 |
| 1144 | OCN132 | HP12038 | Australia | Perth | *Unknown* | *hspSEurope* | 1654828 | 32 | 38.5 | 1550 |
| 1145 | OCN133 | HP12053 | Australia | Perth | *Unknown* | *hspSEurope* | 1615541 | 32 | 39.0 | 1519 |
| 1146 | OCN134 | HP12054 | Australia | Perth | *Unknown* | *hpAfrica2* | 1580448 | 32 | 38.7 | 1473 |
| 1147 | OCN135 | HP12059 | Australia | Perth | *Unknown* | *hspSEurope* | 1630575 | 35 | 38.2 | 1520 |
| 1148 | OCN136 | HP12060 | Australia | Perth | *Unknown* | *hspNEurope* | 1580522 | 38 | 38.8 | 1467 |
| 1149 | OCN137 | HP12064 | Australia | Perth | *Unknown* | *hspEastAsia* | 1575661 | 26 | 38.2 | 1496 |
| 1150 | OCN138 | HP12068 | Australia | Perth | *Unknown* | *hspSEurope* | 1609673 | 31 | 38.4 | 1516 |
| 1151 | OCN139 | HP12069 | Australia | Perth | *Unknown* | *hspEastAsia* | 1595644 | 36 | 38.2 | 1480 |
| 1152 | OCN140 | HP12070 | Australia | Perth | *Unknown* | *hspNEurope* | 1551236 | 38 | 38.6 | 1445 |
| 1153 | OCN141 | HP12073 | Australia | Perth | *Unknown* | *hpAfrica2* | 1606420 | 36 | 38.4 | 1495 |
| 1154 | OCN142 | HP12077 | Australia | Perth | *Unknown* | *hspSEurope* | 1598900 | 27 | 41.1 | 1495 |
| 1155 | OCN143 | HP12078 | Australia | Perth | *Unknown* | *hspEastAsia* | 1618888 | 34 | 37.7 | 1544 |
| 1156 | OCN144 | HP13005 | Australia | Perth | *Unknown* | *hspSWEuropeAustralia* | 1580020 | 38 | 38.8 | 1481 |
| 1157 | OCN145 | HP13007 | Australia | Perth | *Unknown* | *hspSEurope* | 1606819 | 36 | 38.5 | 1494 |
| 1158 | OCN146 | HP13009 | Australia | Perth | *Unknown* | *hspNEuropeAustralia* | 1654194 | 46 | 38.5 | 1549 |
| 1159 | OCN147 | HP13011 | Australia | Perth | *Unknown* | *hspAfrica1SAfrica* | 1597798 | 44 | 38.2 | 1490 |
| 1160 | OCN148 | HP13012 | Australia | Perth | *Unknown* | *hspEastAsia* | 1633006 | 28 | 38.5 | 1551 |
| 1161 | OCN149 | HP13013 | Australia | Perth | *Unknown* | *hspEastAsia* | 1661012 | 38 | 38.1 | 1577 |
| 1162 | OCN150 | HP13021 | Australia | Perth | *Unknown* | *hspSEurope* | 1577986 | 34 | 38.7 | 1502 |
| 1163 | OCN151 | HP13022 | Australia | Perth | *Unknown* | *hspNEurope* | 1594896 | 45 | 39.0 | 1492 |
| 1164 | OCN152 | HP13024 | Australia | Perth | *Unknown* | *hspNEurope* | 1641180 | 46 | 38.4 | 1540 |
| 1165 | OCN153 | HP13025 | Australia | Perth | *Unknown* | *hspNEurope* | 1627628 | 26 | 38.4 | 1527 |
| 1166 | OCN154 | HP13026 | Australia | Perth | *Unknown* | *hspAfrica1SAfrica* | 1676191 | 50 | 38.1 | 1554 |
| 1167 | OCN155 | HP13027 | Australia | Perth | *Unknown* | *hspNEurope* | 1585567 | 59 | 38.1 | 1505 |
| 1168 | OCN156 | HP13028 | Australia | Perth | *Unknown* | *hspSEurope* | 1649898 | 38 | 38.3 | 1569 |
| 1169 | OCN157 | HP13029 | Australia | Perth | *Unknown* | *hspNEurope* | 1564695 | 30 | 38.6 | 1467 |
| 1170 | OCN158 | HP13031 | Australia | Perth | *Unknown* | *hspAmerind* | 1588083 | 47 | 38.7 | 1492 |
| 1171 | OCN159 | HP13033 | Australia | Perth | *Unknown* | *hspNEurope* | 1648519 | 46 | 38.1 | 1540 |
| 1172 | OCN160 | HP13050 | Australia | Perth | *Unknown* | *hspSWEuropeAustralia* | 1580589 | 31 | 38.3 | 1484 |
| 1173 | OCN161 | HP13054 | Australia | Perth | *Unknown* | *hspSEurope* | 1578262 | 29 | 39.1 | 1481 |
| 1174 | OCN162 | HP13056 | Australia | Perth | *Unknown* | *hspNEurope* | 1588857 | 37 | 38.3 | 1486 |
| 1175 | OCN163 | HP13061 | Australia | Perth | *Unknown* | *hspAmerind* | 1706070 | 51 | 37.7 | 1620 |
| 1176 | OCN164 | HP13063 | Australia | Perth | *Unknown* | *hspEastAsia* | 1616602 | 24 | 39.2 | 1543 |
| 1177 | OCN165 | HP13064 | Australia | Perth | *Unknown* | *hspSEurope* | 1664798 | 46 | 38.5 | 1546 |
| 1178 | OCN166 | HP13068 | Australia | Perth | *Unknown* | *hspSWEuropeAustralia* | 1639150 | 24 | 38.2 | 1529 |
| 1179 | OCN167 | HP13072 | Australia | Perth | *Unknown* | *hspAfrica1SAfrica* | 1674524 | 39 | 38.2 | 1556 |
| 1180 | OCN168 | HP14021 | Australia | Perth | *Unknown* | *hspSWEuropeAustralia* | 1625282 | 34 | 38.0 | 1511 |
| 1181 | OCN169 | HP14023 | Australia | Perth | *Unknown* | *hspAfrica1SAfrica* | 1649904 | 29 | 38.6 | 1542 |
| 1182 | OCN170 | HP14031 | Australia | Perth | *Unknown* | *hspEastAsia* | 1565090 | 24 | 38.5 | 1487 |
| 1183 | OCN171 | HP14036 | Australia | Perth | *Unknown* | *hspNEuropeAustralia* | 1627696 | 46 | 38.1 | 1526 |
| 1184 | OCN172 | HP14039 | Australia | Perth | *Unknown* | *hspSWEuropeAustralia* | 1678260 | 1 | 38.7 | 1574 |
| 1185 | OCN173 | HP14048 | Australia | Perth | *Unknown* | *hspSEurope* | 1602417 | 42 | 38.4 | 1470 |
| 1186 | OCN174 | HP14050 | Australia | Perth | *Unknown* | *hspSWEuropeSouthAmerica* | 1583336 | 32 | 38.6 | 1465 |
| 1187 | OCN175 | HP14051 | Australia | Perth | *Unknown* | *hspEastAsia* | 1555159 | 24 | 39.1 | 1485 |
| 1188 | OCN176 | HP14052 | Australia | Perth | *Unknown* | *hspEastAsia* | 1551755 | 32 | 38.8 | 1487 |
| 1189 | OCN177 | HP14054 | Australia | Perth | *Unknown* | *hspSEurope* | 1581075 | 28 | 39.1 | 1492 |
| 1190 | OCN178 | HP14056 | Australia | Perth | *Unknown* | *hspEastAsia* | 1615732 | 28 | 38.7 | 1524 |
| 1191 | OCN179 | HP14065 | Australia | Perth | *Unknown* | *hspSEurope* | 1632634 | 34 | 38.7 | 1519 |
| 1192 | OCN180 | HP14069 | Australia | Perth | *Unknown* | *hspEastAsia* | 1619950 | 39 | 37.9 | 1539 |
| 1193 | OCN181 | HP15002 | Australia | Perth | *Unknown* | *hspEastAsia* | 1582835 | 27 | 39.0 | 1513 |
| 1194 | OCN182 | HP15003 | Australia | Perth | *Unknown* | *hspEastAsia* | 1598068 | 31 | 38.8 | 1515 |
| 1195 | OCN183 | HP15004 | Australia | Perth | *Unknown* | *hspEastAsia* | 1651366 | 24 | 37.7 | 1556 |
| 1196 | OCN184 | HP15005 | Australia | Perth | *Unknown* | *hspSWEuropeAustralia* | 1558820 | 44 | 38.7 | 1452 |
| 1197 | OCN185 | HP15006 | Australia | Perth | *Unknown* | *hspSEurope* | 1652308 | 36 | 38.5 | 1554 |
| 1198 | OCN186 | HP15011 | Australia | Perth | *Unknown* | *hspEastAsia* | 1652125 | 31 | 37.6 | 1562 |
| 1199 | OCN187 | HP15012 | Australia | Perth | *Unknown* | *hspSEurope* | 1600252 | 40 | 39.1 | 1509 |
| 1200 | OCN188 | HP15013 | Australia | Perth | *Unknown* | *hspSEurope* | 1564467 | 47 | 38.2 | 1496 |
| 1201 | OCN189 | HP15015 | Australia | Perth | *Unknown* | *hspEastAsia* | 1545257 | 37 | 38.6 | 1484 |
| 1202 | OCN190 | HP15018 | Australia | Perth | *Unknown* | *hspEastAsia* | 1544845 | 21 | 38.7 | 1472 |
| 1203 | OCN191 | HP15020 | Australia | Perth | *Unknown* | *hspEastAsia* | 1573363 | 27 | 38.9 | 1493 |
| 1204 | OCN192 | HP15022 | Australia | Perth | *Unknown* | *hspSEurope* | 1658050 | 53 | 38.5 | 1552 |
| 1205 | OCN193 | HP15025 | Australia | Perth | *Unknown* | *hspSEurope* | 1684227 | 38 | 38.2 | 1564 |
| 1206 | OCN194 | HP15026 | Australia | Perth | *Unknown* | *hspSEurope* | 1693128 | 43 | 38.6 | 1593 |
| 1207 | OCN195 | HP15027 | Australia | Perth | *Unknown* | *hspEastAsia* | 1582609 | 26 | 38.7 | 1494 |
| 1208 | OCN196 | HP15028 | Australia | Perth | *Unknown* | *hspSEurope* | 1631173 | 42 | 38.6 | 1538 |
| 1209 | OCN197 | HP15031 | Australia | Perth | *Unknown* | *hspEastAsia* | 1605490 | 38 | 38.6 | 1503 |
| 1210 | OCN198 | HP15032 | Australia | Perth | *Unknown* | *hspEastAsia* | 1555248 | 29 | 38.6 | 1468 |
| 1211 | OCN199 | HP15033 | Australia | Perth | *Unknown* | *hspSEurope* | 1571081 | 54 | 38.4 | 1481 |
| 1212 | OCN200 | HP15034 | Australia | Perth | *Unknown* | *hspAfrica1WAfrica* | 1628897 | 45 | 39.3 | 1512 |
| 1213 | OCN201 | HP15035 | Australia | Perth | *Unknown* | *hspSEurope* | 1605083 | 60 | 38.2 | 1505 |
| 1214 | OCN202 | HP15036 | Australia | Perth | *Unknown* | *hspEastAsia* | 1536706 | 43 | 38.4 | 1454 |
| 1215 | OCN203 | HP15039 | Australia | Perth | *Unknown* | *hspNEuropeAustralia* | 1588463 | 86 | 38.2 | 1483 |
| 1216 | OCN204 | HP15040 | Australia | Perth | *Unknown* | *hspEastAsia* | 1597365 | 19 | 38.9 | 1508 |
| 1217 | OCN205 | HP15044 | Australia | Perth | *Unknown* | *hspAmerind* | 1641358 | 46 | 38.8 | 1538 |
| 1218 | OCN206 | HP15050 | Australia | Perth | *Unknown* | *hspSEurope* | 1657276 | 42 | 38.9 | 1550 |
| 1219 | OCN207 | HP15051 | Australia | Perth | *Unknown* | *hspEastAsia* | 1610206 | 26 | 38.6 | 1529 |
| 1220 | OCN208 | HP15054 | Australia | Perth | *Unknown* | *hspSEurope* | 1610877 | 34 | 39.0 | 1509 |
| 1221 | OCN209 | HP15058 | Australia | Perth | *Unknown* | *hspSEurope* | 1549150 | 28 | 38.7 | 1459 |
| 1222 | OCN210 | HP15059 | Australia | Perth | *Unknown* | *hspAfrica1SAfrica* | 1660590 | 57 | 38.6 | 1525 |
| 1223 | OCN211 | HP15060 | Australia | Perth | *Unknown* | *hspSEurope* | 1620688 | 36 | 39.4 | 1515 |
| 1224 | OCN212 | HP15067 | Australia | Perth | *Unknown* | *hspEastAsia* | 1581081 | 25 | 38.8 | 1499 |
| 1225 | OCN213 | HP16001 | Australia | Perth | *Unknown* | *hpSahul* | 1701459 | 43 | 37.9 | 1595 |
| 1226 | OCN214 | HP16004 | Australia | Perth | *Unknown* | *hspEastAsia* | 1521290 | 21 | 39.1 | 1461 |
| 1227 | OCN215 | HP16008 | Australia | Perth | *Unknown* | *hspSWEurope* | 1648542 | 32 | 38.9 | 1548 |
| 1228 | OCN216 | ausabras47a | Australia | AliceSprings | *hpSahul* | *hpSahul* | 1579248 | 149 | 39.4 | 1453 |
| 1229 | SAM001 | Arg_53RA1 | Argentina | BuenosAires | *hspSWEurope* | *hspSWEurope* | 1625188 | 29 | 38.6 | 1511 |
| 1230 | SAM002 | Arg_110 | Argentina | Unknown | *hspSWEurope* | *hspSWEurope* | 1612882 | 43 | 38.8 | 1502 |
| 1231 | SAM003 | Arg_118EC2 | Argentina | BuenosAires | *hspSWEurope* | *hspSWEuropeSouthAmerica* | 1575110 | 46 | 37.4 | 1479 |
| 1232 | SAM004 | Arg_19RC1 | Argentina | BuenosAires | *hspSWEuropeMexico* | *hspSWEurope* | 1652983 | 44 | 38.4 | 1520 |
| 1233 | SAM005 | Arg_23UA1 | Argentina | BuenosAires | *hspAfrica1MiscAmerica* | *hspSWEuropeNorthAmerica* | 1695597 | 38 | 36.7 | 1565 |
| 1234 | SAM006 | Arg_34UA2 | Argentina | BuenosAires | *hspSWEuropeMexico* | *hspSWEuropeSouthAmerica* | 1640908 | 26 | 38.6 | 1512 |
| 1235 | SAM007 | Arg_38 | Argentina | Unknown | *hspSWEuropeMexico* | *hspSWEuropeSouthAmerica* | 1686585 | 41 | 38.9 | 1557 |
| 1236 | SAM008 | Arg_38RA2 | Argentina | BuenosAires | *hspSEurope* | *hspSEurope* | 1650898 | 25 | 38.8 | 1544 |
| 1237 | SAM009 | Arg_39UA2 | Argentina | BuenosAires | *hspSWEuropeMexico* | *hspSWEuropeSouthAmerica* | 1666392 | 52 | 37.8 | 1530 |
| 1238 | SAM010 | Arg_7RA1 | Argentina | BuenosAires | *hspSEurope* | *hspSEurope* | 1567871 | 23 | 38.7 | 1471 |
| 1239 | SAM011 | HPARG63 | Argentina | Unknown | *hspSWEuropeMexico* | *hspSWEuropeSouthAmerica* | 1667776 | 28 | 37.9 | 1553 |
| 1240 | SAM012 | HPARG8G | Argentina | Unknown | *hspSEurope* | *hspSEurope* | 1602229 | 47 | 38.2 | 1509 |
| 1241 | SAM013 | Bra_LPB32 | Brazil | BeloHorizonte | *hspSEurope* | *hspAfrica1NAmerica* | 1651604 | 45 | 38.0 | 1521 |
| 1242 | SAM014 | Bra_LPB33 | Brazil | BeloHorizonte | *hspAfrica1NAmerica* | *hspSWEurope* | 1584871 | 27 | 38.9 | 1475 |
| 1243 | SAM015 | Bra_LPB34 | Brazil | SantoAntoniodoGrana | *hspSWEurope* | *hspAfrica1NAmerica* | 1590058 | 30 | 39.0 | 1483 |
| 1244 | SAM016 | Bra_LPB35 | Brazil | BeloHorizonte | *hspAfrica1NAmerica* | *hspAfrica1NAmerica* | 1655842 | 35 | 38.0 | 1529 |
| 1245 | SAM017 | Bra_LPB37 | Brazil | RaulSoares | *hspAfrica1NAmerica* | *hspAfrica1NAmerica* | 1674158 | 26 | 38.8 | 1562 |
| 1246 | SAM018 | Bra_LPB40 | Brazil | BeloHorizonte | *hspAfrica1NAmerica* | *hspAfrica1NAmerica* | 1630354 | 35 | 39.2 | 1504 |
| 1247 | SAM019 | Bra_LPB41 | Brazil | BeloHorizonte | *hspAfrica1NAmerica* | *hspAfrica1NAmerica* | 1667369 | 35 | 39.0 | 1544 |
| 1248 | SAM020 | Bra_LPB42 | Brazil | BeloHorizonte | *hspAfrica1NAmerica* | *hspAfrica1SAfrica* | 1631908 | 45 | 37.8 | 1508 |
| 1249 | SAM021 | Bra_LPB43 | Brazil | BeloHorizonte | *hspAfrica1MiscAmerica* | *hspAfrica1NAmerica* | 1726318 | 115 | 38.3 | 1624 |
| 1250 | SAM022 | Bra_LPB44 | Brazil | Esmeraldas | *hspAfrica1NAmerica* | *hspAfrica1NAmerica* | 1597687 | 34 | 38.9 | 1478 |
| 1251 | SAM023 | CL_FE60 | Chile | Central | *hspSWEuropeMexico* | *hspSWEuropeSouthAmerica* | 1686794 | 40 | 36.8 | 1577 |
| 1252 | SAM024 | CL-FE70 | Chile | Central | *hspSWEuropeMexico* | *hspSWEuropeSouthAmerica* | 1594247 | 92 | 39.0 | 1463 |
| 1253 | SAM025 | CL_MASP | Chile | Central | *hspSWEurope* | *hspSWEuropeColombia* | 1615490 | 16 | 38.6 | 1521 |
| 1254 | SAM026 | Cuz20 | Peru | MonteCarmelo | *hspSAmerind* | *hspAmerind* | 1635449 | 1 | 38.9 | 1534 |
| 1255 | SAM027 | PeCan18 | Peru | Lima | *hspAfrica1MiscAmerica* | *hspSWEuropeNorthAmerica* | 1660685 | 1 | 39.0 | 1541 |
| 1256 | SAM028 | PeCan4 | Peru | Unknown | *hspSAmerind* | *hspAmerind* | 1638269 | 2 | 35.9 | 1531 |
| 1257 | SAM029 | Puno120 | Peru | Puno | *hspSAmerind* | *hspAmerind* | 1637762 | 2 | 37.4 | 1530 |
| 1258 | SAM030 | Puno135 | Peru | Puno | *hspSAmerind* | *hspAmerind* | 1646139 | 1 | 38.8 | 1537 |
| 1259 | SAM031 | Sat464 | Peru | Satipo | *hspSAmerind* | *hspAmerind* | 1567570 | 2 | 36.3 | 1462 |
| 1260 | SAM032 | Shi112 | Peru | Shimaa | *hspSAmerind* | *hspAmerind* | 1663456 | 1 | 38.8 | 1558 |
| 1261 | SAM033 | Shi169 | Peru | Shimaa | *hspSAmerind* | *hspAmerind* | 1616909 | 1 | 38.9 | 1515 |
| 1262 | SAM034 | Shi417 | Peru | Shimaa | *hspSAmerind* | *hspAmerind* | 1665719 | 1 | 38.8 | 1542 |
| 1263 | SAM035 | Shi470 | Peru | Shimaa | *hspSAmerind* | *hspAmerind* | 1608548 | 1 | 38.9 | 1517 |
| 1264 | SAM036 | SJM180 | Peru | Lima | *hspSWEuropeMexico* | *hspSWEuropeSouthAmerica* | 1658051 | 1 | 38.9 | 1521 |
| 1265 | SAM037 | v225d | Venezuela | Piaroa | *hspSAmerind* | *hspAmerind* | 1595604 | 2 | 35.9 | 1509 |
| 1266 | SAM038 | VE_HET009 | Venezuela | Caracas | *hspSWEurope* | *hspSWEurope* | 1630453 | 42 | 38.5 | 1522 |
| 1267 | SAM039 | VE_HET044 | Venezuela | Caracas | *hspSWEuropeMexico* | *hspSWEuropeSouthAmerica* | 1620359 | 29 | 38.3 | 1516 |
| 1268 | SAM040 | VE_HET046 | Venezuela | Caracas | *hspAfrica1MiscAmerica* | *hspSWEuropeNorthAmerica* | 1657590 | 36 | 37.6 | 1531 |
| 1269 | SAM041 | VE_HET052 | Venezuela | Caracas | *hspSWEurope* | *hspSWEurope* | 1668255 | 31 | 37.1 | 1574 |
| 1270 | SAM042 | VE_HET057 | Venezuela | Caracas | *hspSWEurope* | *hspSWEurope* | 1643379 | 45 | 38.9 | 1522 |
| 1271 | SAM043 | VE_HET084 | Venezuela | Caracas | *hspSWEuropeMexico* | *hspSWEuropeSouthAmerica* | 1764504 | 81 | 38.1 | 1624 |
| 1272 | SAM044 | VE_HET109 | Venezuela | Caracas | *hspSWEurope* | *hspSWEurope* | 1649942 | 39 | 37.6 | 1540 |
| 1273 | SAM045 | VE_HV055 | Venezuela | Caracas | *hspAfrica1MiscAmerica* | *hspAfrica1SAfrica* | 1598319 | 23 | 37.9 | 1482 |
| 1274 | SAM046 | VE_HV073 | Venezuela | Caracas | *hspSWEurope* | *hspSWEurope* | 1609082 | 54 | 37.7 | 1501 |
| 1275 | SAM047 | VE_HV076 | Venezuela | Caracas | *hspSWEurope* | *hspSWEurope* | 1656411 | 48 | 36.4 | 1553 |
| 1276 | SAM048 | VE_HV265 | Venezuela | Caracas | *hspSWEurope* | *hspSWEurope* | 1635921 | 43 | 37.0 | 1549 |

**Supplemental table 2.** *H. pylori* subpopulations identified in the 163 Colombian strains analyzed.

| **Colombian *H. pylori* subpopulations** | | **Number of strains** | | **Departments** | | | | | | | | | | |
| --- | --- | --- | --- | --- | --- | --- | --- | --- | --- | --- | --- | --- | --- | --- |
|  |  |  |  | **CU** | **NR** | **BY** | **TL** | **ST** | **CL** | **MT** | **CQ** | **CV** | **RS** | **AM** |
| *hspSEurope* | | 1 | | 1 |  |  |  |  |  |  |  |  |  |  |
| *hspSWEurope* | | 1 | | 1 |  |  |  |  |  |  |  |  |  |  |
| hspSWEuropeNorthAmerica | | 13 | | 6 |  | 1 | 2 | 2 |  |  |  | 1 | 1 |  |
| *hspSWEuropeSouthAmerica* | | 6 | | 5 |  |  |  | 1 |  |  |  |  |  |  |
| *hspSWEuropeColombia* | | 20 | | 13 | 1 | 5 |  | 1 |  |  |  |  |  |  |
| *hspColombia* | *hspColombiaNariño* | 112 | 14 |  | 14 |  |  |  |  |  |  |  |  |  |
|  | *hspColombiaAndesMountain* |  | 29 | 8 | 13 | 3 | 1 | 2 | 1 |  | 1 |  |  |  |
|  | *hspColombiaCundinamarca* |  | 69 | 49 | 1 | 16 |  | 2 |  | 1 |  |  |  |  |
| *hspAfrica1SAfrica* | | 1 | |  | 1 |  |  |  |  |  |  |  |  |  |
| *hspAfrica1NAmerica* | | 7 | | 1 | 6 |  |  |  |  |  |  |  |  |  |
| *hspAmerind* | | 2 | |  |  |  |  |  |  |  |  |  |  | 2 |
| Total | | 163 | | 84 | 36 | 25 | 3 | 8 | 1 | 1 | 1 | 1 | 1 | 2 |

CU: Cundinamarca

NR:Nariño

BY:Boyaca

TL:Tolima

ST:Santander

CL:Caldas

MT:Meta

CQ:Caqueta

CV:Cauca's Valley

RS:Risaralda

AM:Amazon

**Supplemental table 3.** Core genome positions of highest Fst values in *hspColombia* isolates compared with *hspSWEurope.*

| **Gene** | **SNP** | **Alleles** | | **Fst value** | **Amino acid change (Codon)** | **Sustitution class** |
| --- | --- | --- | --- | --- | --- | --- |
|  |  | **Reference** | **Alternative** |  |  |  |
| *HP0018* | 18007 | G | A | 0.678227 | *p.Ser359Asn* | Nonsynonymous |
| *HP0019* | 19187 | A | G | 0.66627 | *p.Asn270Asp* | Nonsynonymous |
| *HP0127* | 139203 | C | T | 0.622084 | *p.Thr266Val* | Nonsynonymous |
| *HP0127* | 139202 | A | G | 0.608849 | *p.Thr266Val* | Nonsynonymous |
| *HP0130* | 140740 | T | C | 0.635371 | *p.Ile96Val* | Nonsynonymous |
| *HP0175* | 182158 | G | A | 0.641939 | *p.Gln98Gln* | Synonymous |
| *HP0175* | 182168 | A | C | 0.641939 | *p.Asp101Asp* | Synonymous |
| *HP0175* | 182161 | A | G | 0.630515 | *p.Lys99Lys* | Synonymous |
| *HP0181* | 188183 | T | C | 0.500663 | *p.Leu69Leu* | Synonymous |
| *HP0252* | 262472 | G | A | 0.635204 | *p.Gly252Arg* | Nonsynonymous |
| *HP0252* | 262533 | A | G | 0.559515 | *p.Asn272Ser* | Nonsynonymous |
| *HP0252* | 262591 | A | G | 0.529252 | *p.Leu291Leu* | Synonymous |
| *HP0252* | 262576 | C | T | 0.525522 | *p.Ile286IIe* | Synonymous |
| *HP0269* | 279397 | C | T | 0.504857 | *p.Cys251Cys* | Synonymous |
| *HP0377* | 385622 | T | C | 0.674541 | *p.Phe95Leu* | Nonsynonymous |
| *HP0377* | 385885 | C | G | 0.615209 | *p.Ala182Ala* | Synonymous |
| *HP0377* | 385608 | C | A,T | 0.540745 | *p.Ser90His,Phe* | Nonsynonymous |
| *HP0377* | 385607 | T | C | 0.537553 | *p.Ser90His,Phe* | Nonsynonymous |
| *HP0415* | 428204 | G | A | 0.50346 | *p.Gly173Asn,Asp* | Nonsynonymous |
| *HP0486* | 510277 | A | G | 0.937532 | *p.Asp309Ser* | Nonsynonymous |
| *HP0486* | 510233 | C | T | 0.915792 | *p.Ser,Asn294Ser* | Nonsynonymous |
| *HP0486* | 510226 | A | G | 0.895824 | *p.Gln292Gly* | Nonsynonymous |
| *HP0486* | 510268 | C | T | 0.870624 | *p.Ala306Val* | Nonsynonymous |
| *HP0486* | 510212 | C | T | 0.824388 | *p.Tyr287Tyr* | Synonymous |
| *HP0486* | 510238 | G | T | 0.709831 | *p.Arg296Met* | Nonsynonymous |
| *HP0486* | 509477 | T | C | 0.654031 | *p.Ala42Ser* | Nonsynonymous |
| *HP0486* | 510245 | T | A,G | 0.612806 | *p.Ile298Met,Ile* | Nonsynonymous |
| *HP0486* | 510275 | T | C | 0.937532 | *p.Tyr308Tyr* | Synonymous |
| *HP0486* | 510276 | G | A | 0.937532 | *p.Asp309Ser* | Nonsynonymous |
| *HP0486* | 510224 | T | G | 0.904227 | *p.Gln292Gly* | Nonsynonymous |
| *HP0486* | 510272 | G | A | 0.904227 | *p.Leu307Leu* | Synonymous |
| *HP0486* | 510416 | T | C | 0.712921 | *p.Asp355Asp* | Synonymous |
| *HP0486* | 510251 | C | T | 0.709183 | *p.Asn300Asn* | Synonymous |
| *HP0486* | 510263 | C | T | 0.683289 | *p.Phe304Phe* | Synonymous |
| *HP0486* | 509475 | G | T | 0.678643 | *p.Ser,Ala42Ser* | Nonsynonymous |
| *HP0486* | 509474 | T | C | 0.654031 | *p.Phe41Phe* | Synonymous |
| *HP0486* | 510152 | C | T | 0.635819 | *p.Ile267IIe* | Synonymous |
| *HP0486* | 510170 | T | C | 0.595154 | *p.Leu273Leu* | Synonymous |
| *HP0517* | 545233 | A | G | 0.609185 | *p.Asp,Arg300Gly* | Nonsynonymous |
| *HP0517* | 545234 | A | G | 0.609185 | *p.Asp,Arg300Gly* | Nonsynonymous |
| *HP0517* | 545237 | G | T | 0.609185 | *p.Arg301Met* | Nonsynonymous |
| *HP0517* | 545227 | A | C | 0.528569 | *p.Arg,Lys297Gln* | Nonsynonymous |
| *HP0554* | 589650 | A | G | 0.687286 | *p.Ser295Gly* | Nonsynonymous |
| *HP0554* | 589649 | C | T | 0.635204 | *p.Ser295Gly* | Nonsynonymous |
| *HP0554* | 589660 | G | A | 0.568985 | *p.Arg298Lys* | Nonsynonymous |
| *HP0554* | 589661 | G | A | 0.568985 | *p.Arg298Lys* | Nonsynonymous |
| *HP0554* | 589664 | G | A | 0.561235 | *p.Arg299Arg* | Synonymous |
| *HP0554* | 589452 | G | A | 0.548769 | *p.Asp229Asn* | Nonsynonymous |
| *HP0558* | 592566 | C | T | 0.506588 | *p.Ile26Ile* | Synonymous |
| *HP0564* | 597177 | T | C | 0.625088 | *p.Ser51Ser* | Synonymous |
| *HP0686* | 735762 | A | G | 0.54699 | *p.Val462Ala* | Nonsynonymous |
| *HP0686* | 735764 | C | T | 0.54699 | *p.Ile463Ile* | Synonymous |
| *HP0706* | 760895 | A | C | 0.86328 | *p.Lys208Asn* | Nonsynonymous |
| *HP0706* | 760533 | C | T | 0.65096 | *p.Leu88Phe* | Nonsynonymous |
| *HP0706* | 760505 | T | C | 0.572761 | *p.Ala78Ser* | Nonsynonymous |
| *HP0706* | 760502 | T | C | 0.556793 | *p.His77His* | Synonymous |
| *HP0706* | 760503 | G | T | 0.556793 | *p.Ala78Ser* | Nonsynonymous |
| *HP0706* | 760755 | A | G | 0.548039 | *p.Ile162Val* | Nonsynonymous |
| *HP0706* | 760641 | A | G | 0.526858 | *p.Thr124Asp* | Nonsynonymous |
| *HP0706* | 760642 | C | A | 0.526858 | *p.Thr124Asp* | Nonsynonymous |
| *HP0706* | 760580 | T | A,G | 0.52683 | *p.Phe103Leu* | Nonsynonymous |
| *HP0788* | 844036 | G | A | 0.623103 | *p.Ala120Thr* | Nonsynonymous |
| *HP1012* | 1076440 | G | A | 0.589325 | *p.Arg299Lys* | Nonsynonymous |
| *HP1054* | 1117241 | T | C | 0.535516 | *p.Glu2Gly* | Nonsynonymous |
| *HP1055* | 1117712 | G | A | 0.717047 | *p.Ala163Val* | Nonsynonymous |
| *HP1055* | 1117671 | T | A | 0.663259 | *p.Ile177Phe* | Nonsynonymous |
| *HP1055* | 1117630 | A | G | 0.65924 | *p.Asp190Asp* | Synonymous |
| *HP1055* | 1117256 | T | C | 0.650959 | *p.Ter315Ter* | Synonymous |
| *HP1055* | 1117378 | T | C | 0.523366 | *p.Arg274Arg* | Synonymous |
| *HP1055* | 1117348 | T | G | 0.509373 | *p.Gln284His* | Nonsynonymous |
| *HP1056* | 1118775 | A | G | 0.511819 | *p.Ser92Ser* | Synonymous |
| *HP1340* | 1399617 | A | G | 0.63672 | *p.Ile29Val* | Nonsynonymous |
| *HP1487* | 1559900 | T | A,G | 0.540672 | *p.Arg209Arg* | Synonymous |
| *HP1487* | 1559824 | G | A | 0.520396 | *p.Leu235Phe* | Nonsynonymous |
| *HP1487* | 1559954 | G | A | 0.519794 | *p.Leu191Leu* | Synonymous |
| *HP1512* | 1586602 | G | A | 0.799812 | *p.Arg719Gln* | Nonsynonymous |
| *HP1512* | 1586601 | A | C | 0.788448 | *p.Arg719Gln* | Nonsynonymous |
| *HP1512* | 1586609 | G | A | 0.788448 | *p.Lys721Lys* | Synonymous |
| *HP1512* | 1586603 | G | A | 0.775948 | *p.Arg719Gln* | Nonsynonymous |
| *HP1512* | 1584617 | T | C | 0.53367 | *p.Ser57Ser* | Synonymous |
| *HP1565* | 1648838 | G | A | 0.584235 | *p.Thr66Ile* | Nonsynonymous |
| *HP1565* | 1648872 | T | C | 0.503984 | *p.Asn55Asp* | Nonsynonymous |
